# Divergent malignant evolution converges on shared immunosuppressive ecosystems through tumor-intrinsic TMED2–ADAR1-dependent complement signalling in multifocal hepatocellular carcinoma

**DOI:** 10.64898/2026.04.21.719788

**Authors:** Cheng Wu, Ke Si, Shuai Mao, Zehang Jiang, Douyue Li, Haipeng Zhu, Ningning Sun, Weiling Liang, Wenliang Zhang

## Abstract

**Background:** Multifocal hepatocellular carcinoma (HCC) comprises spatially separated tumor foci with distinct evolutionary trajectories, posing a major therapeutic challenge. Whether divergent lesions retain shared immune-evasion mechanisms beyond lesion-specific heterogeneity and offer cross-focal therapeutic opportunities remains unknown.

**Objective:** To resolve malignant and microenvironmental heterogeneity across tumor foci and identify conserved immunosuppressive programs underlying multifocal HCC progression.

**Design:** Matched tumour foci and peritumoural tissues from patients with multifocal HCC underwent integrated single-cell and spatial multi-omics profiling, including scFAST-seq (n=15), scATAC-seq (n=12), and Stereo-seq (n=14). In-house RNA-seq (n=13), public scRNA-seq (HRA001748) and RNA-seq datasets (TCGA-LIHC, OEP000321) were used for validation. Multiplex immunohistochemistry (n=96) and functional assays validated key findings.

**Results:** Multifocal HCC exhibited extensive inter-focal heterogeneity in malignant genomic, transcriptional, epigenetic, and evolutionary states, accompanied by diverse immune and stromal landscapes. Despite this divergence, spatially separated tumor foci converged on shared malignant states marked by elevated *TMED2* expression and conserved immunosuppressive ecosystems characterized by regulatory T-cell (Treg) and SPP1+/SPRED1+ macrophage enrichment, impaired cytotoxic CD8+ T-cell immunity, and recurrent stromal–vascular remodeling. Mechanistically, malignant hepatocytes engaged SPP1+/SPRED1+ macrophages through C3/C5 complement signaling across divergent tumor foci, while macrophages and Tregs contributed to CD8+ T-cell dysfunction through CD274–PDCD1, CD80/CD86–CTLA4/CD28, and MHC-I–CD8 interactions. A tumor-intrinsic TMED2–ADAR1-dependent A-to-I RNA editing program promoted C3/C5 complement expression and secretion, linking malignant heterogeneity to conserved tumor–immune signaling.

**Conclusions:** Despite divergent malignant evolution, multifocal HCC converges on shared immunosuppressive ecosystems. The TMED2–ADAR1– C3/C5 complement axis represents a conserved tumor–immune vulnerability that may enable therapeutic strategies targeting common immune-evasion mechanisms across heterogeneous tumor foci.

**Graphic Abstract:** 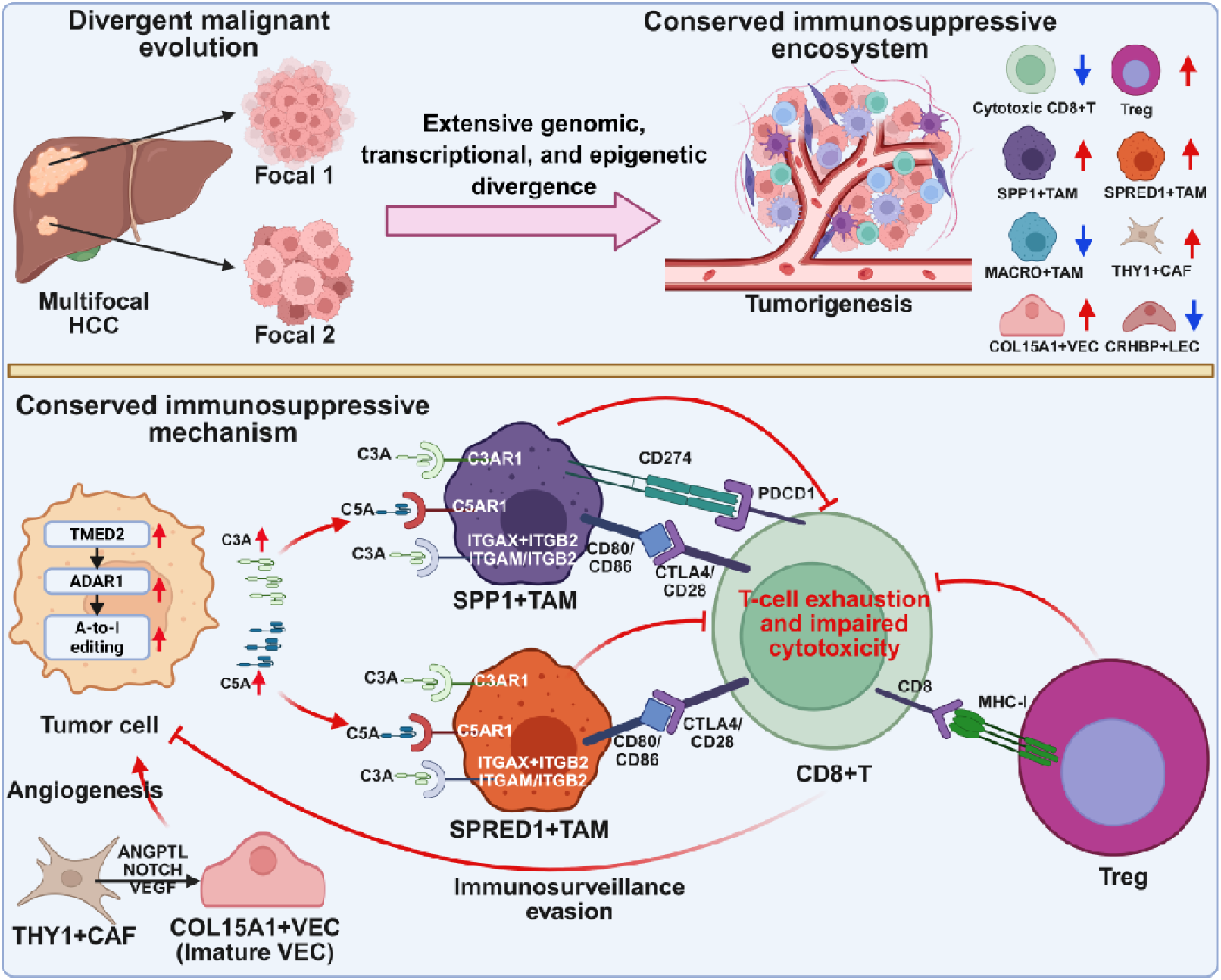

**WHAT IS ALREADY KNOWN ON THIS TOPIC:**

(1) Multifocal HCC comprises spatially separated tumor foci that can arise through distinct evolutionary trajectories, including intrahepatic metastasis and multicentric occurrence.
(2) Genomic, transcriptomic and immunological studies have demonstrated substantial molecular and cellular heterogeneity between tumor foci, but the malignant programs and microenvironmental states underlying this heterogeneity remain poorly defined.
(3) Whether evolutionarily divergent tumor foci retain shared oncogenic programs and conserved tumor–immune mechanisms that sustain immune suppression across multifocal disease remains unknown.

**WHAT THIS STUDY ADDS:**

(1) Integrated single-cell transcriptomic, epigenomic, and spatial profiling reveals extensive inter-focal heterogeneity in malignant genomic, transcriptional, and evolutionary states, accompanied by diverse remodeling of immune, stromal, and vascular compartments.
(2) Despite divergent malignant evolution, spatially separated tumor foci retain shared malignant states marked by recurrent *TMED2* expression and converge on conserved immunosuppressive ecosystems characterized by Treg and SPP1+/SPRED1+ TAM enrichment, impaired cytotoxic CD8+ T-cell immunity, and recurrent stromal–vascular remodeling.
(3) A previously uncharacterized tumor-intrinsic TMED2–ADAR1-dependent A-to-I RNA editing regulatory axis promotes C3/C5 complement signaling in malignant hepatocytes, linking malignant heterogeneity to conserved tumor– immune interactions and identifying a potential cross-focal therapeutic vulnerability.

**HOW THIS STUDY MIGHT AFFECT RESEARCH, PRACTICE OR POLICY:**

(1) The single cell and spatial multi-omic atlas generated in this study provides a valuable resource for investigating inter-focal heterogeneity and identifying shared oncogenic and immunosuppressive programs during multifocal HCC evolution.
(2) The persistence of shared oncogenic and immunosuppressive programs across evolutionarily distinct tumor foci suggests that targeting conserved tumour–immune interactions may complement lesion-specific therapeutic strategies and potentially overcome inter-focal heterogeneity.
(3) The TMED2–ADAR1–C3/C5 complement axis provides a mechanistically defined candidate for therapeutic intervention aimed at disrupting tumor– immune immunosuppression across multifocal HCC.

## INTRODUCTION

Hepatocellular carcinoma (HCC) remains one of the leading causes of cancer-related mortality worldwide^1^. Approximately 35–50% of patients present with multifocal disease at diagnosis^2, 3^, which is associated with increased recurrence, therapeutic resistance, and poor survival compared with unifocal tumors^4^. Multifocal HCC provides a unique clinical model to investigate how malignant evolution influences tumor microenvironment organization. Spatially separated tumor foci may arise through intrahepatic metastasis (IM), reflecting clonal dissemination from an ancestral tumor, or multicentric occurrence (MO), resulting from independent tumorigenic events in chronically injured liver tissue^5^. Such distinct evolutionary origins provide an opportunity to investigate how divergent malignant evolution shapes the cellular and molecular landscape of individual tumor foci and, importantly, whether distinct malignant states retain shared mechanisms of immune evasion.

Recent genomic and multi-omic studies have revealed extensive molecular diversity among spatially separated HCC lesions from the same patient. Whole-genome sequencing has uncovered distinct clonal architectures and mutational landscapes distinguishing IM and MO lesions^5^, while integrative analyses have identified inter-focal differences in HBV integration patterns, copy-number alterations, structural variants and driver alterations^6, 7^. Beyond genomic alterations, transcriptomic^8^, immunogenomic^9^, proteogenomic^10^, and microbiome^11^ studies have further revealed differences in malignant cell states, metabolic programs, immune landscapes, microbe– host interactions, and ecological niches across tumor foci. Single-cell RNA-seq studies have provided additional evidence of heterogeneous cellular states and immune compositions across IM and MO lesions^12^. Collectively, these studies support substantial molecular and cellular heterogeneity among spatially separated tumor foci. However, much of the existing evidence has been generated using bulk omics approaches, in which signals from diverse cell populations are averaged within each lesion. Consequently, the cellular states and spatial organization underlying inter-focal heterogeneity remain incompletely resolved, particularly with respect to how heterogeneous malignant populations interact with surrounding immune and stromal compartments.

Tumor evolution is traditionally viewed as a process in which progressive genetic and epigenetic diversification generates increasingly heterogeneous malignant populations^13^. Yet malignant cells evolve within complex tumor ecosystems in which immune, stromal, and vascular compartments both impose selective pressures and respond to tumor-derived signals^14, 15^. Therefore, divergent evolution may generate distinct malignant states without necessarily producing distinct mechanisms of immune escape. Evolutionarily separated tumor foci may retain convergent tumor–immune interactions that sustain immune evasion despite molecular divergence. Whether such evolutionary convergence of immune suppression exists across multifocal HCC remains largely unknown.

Defining such conserved immune-evasion programs is clinically important because lesion-specific alterations may limit the durability of therapies targeting individual malignant clones. Identifying vulnerabilities shared across evolutionarily distinct tumor foci could enable therapeutic strategies that overcome inter-focal heterogeneity. Here, we integrated cutting-edge single-cell and spatial multi-omics approaches to resolve malignant evolution, cellular states, spatial organization, and tumor–immune interactions across multifocal HCC. This integrated approach revealed extensive inter-focal divergence alongside conserved immunosuppressive signaling, providing a framework to distinguish lesion-specific cellular states from shared functional dependencies and to identify potential therapeutic vulnerabilities across multifocal disease.

## METHODS

### Overview of the patient cohort and study design

We enrolled patients with primary multifocal HCC undergoing surgical resection without prior anticancer treatment and collected paired tumor foci and adjacent tissues. Integrated single-cell full-length transcriptomics (scFAST-seq) (15 tumor/adjacent tissue samples), single-cell chromatin accessibility profiling (scATAC-seq) (12 samples), and high-resolution spatial transcriptomics (Stereo-seq) (14 samples), bulk RNA sequencing (13 samples) (**online supplemental Table 1**), public datasets (HRA001748^16^ cohort (n = 88); TCGA-LIHC (n = 466, UCSC Xena, https://xena.ucsc.edu/); OEP000321^17^ (n = 316)), and multiplex immunohistochemistry (mIHC) array (48 paried samples) (**online supplemental Table 2**) were applied to characterize malignant evolution and conserved tumor microenvironments. Multi-omics analyses were performed to define cellular heterogeneity, chromatin states, spatial organization, RNA editing profiles, copy number alterations, evolutionary trajectories, and intercellular communication. Functional validation, including gene perturbation, proliferation, migration, ELISA, and qPCR assays, was conducted to investigate the identified molecular mechanisms. For further details regarding the materials and methods, please refer to the **supplementary information** and the **online supplemental tables 1-7**.

## RESULTS

### Single-cell and spatial multi-omics delineate heterogeneous yet convergent cellular landscapes in multifocal HCC

To characterize inter-focal heterogeneity and uncover conserved oncogenic and immune-evasion programs across divergent tumor foci in multifocal HCC, we established an integrated single-cell and spatial multi-omics framework. We profiled matched tumor foci and peritumoral tissues from multifocal HCC patients using scFAST-seq (n=15), scATAC-seq (n=12), and Stereo-seq (n=14) (**figure 1a,b, online supplemental figure 1**). To further evaluate clinical relevance, we integrated our in-house bulk RNA-seq cohort (n=13) with large-scale public datasets, including scRNA-seq data from HRA001748 (n=88)^16^ and bulk RNA-seq from TCGA-LIHC (n=466) and OEP000321 (n=316)^17^ (**figure 1b**).

**Figure 1.**
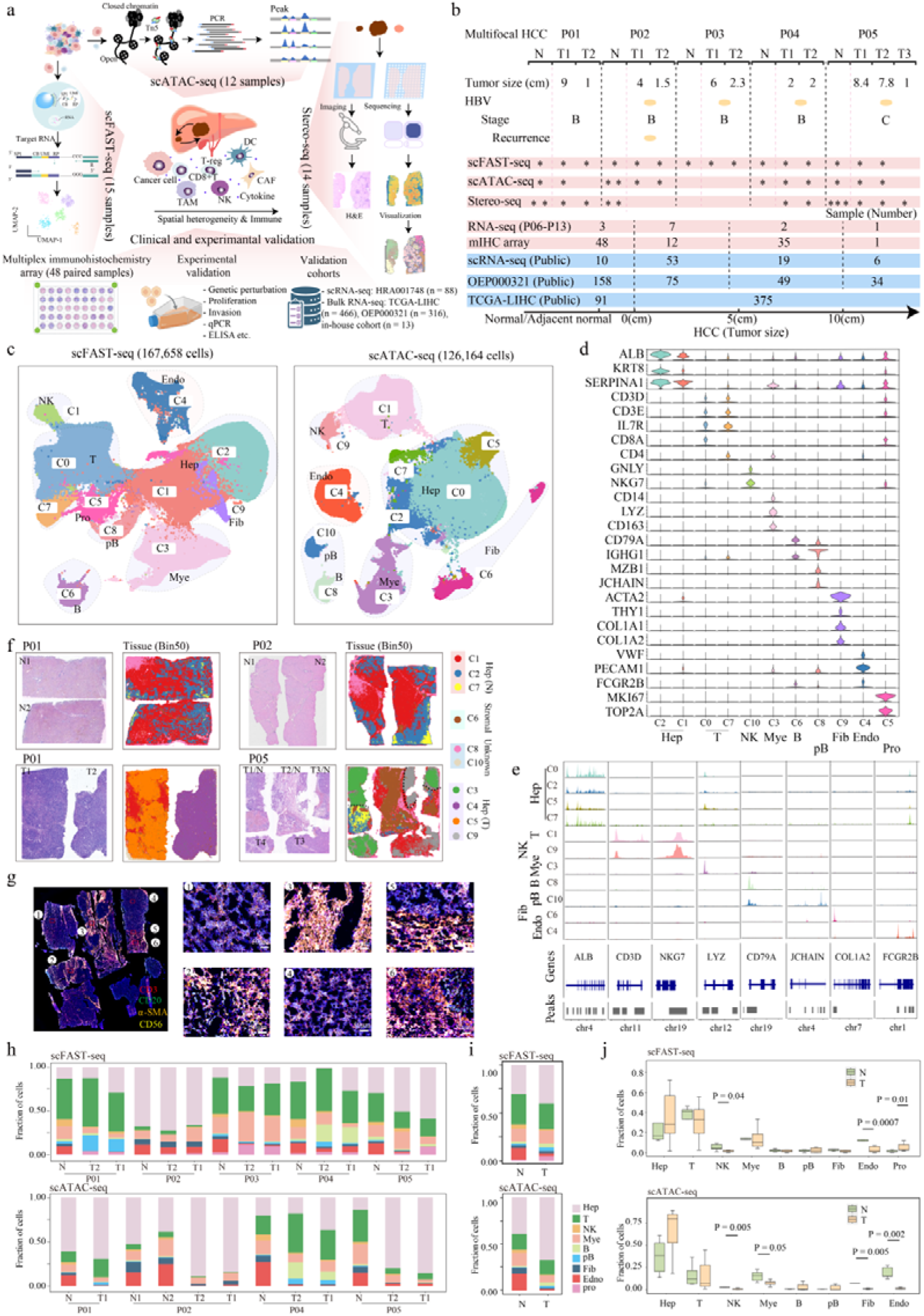
Integrated single-cell and spatial profiling of multifocal HCC. **(a)** Overview of the study design. **(b)** Diagram summarizing sample information contained in different sequencing datasets. **(c)** UMAP plot of cell lineages from scFAST-seq (left) and scATAC-seq (right) in multifocal HCC. **(d, e)** Violin plots showing marker gene expression across cell types in scFAST-seq **(d)** and scATAC-seq **(e)**. **(f)** H&E images and spatial cell-type annotation of representative adjacent normal (N) and tumor (T) tissues. **(g)** Multiplex immunohistochemistry validation of major cell populations and spatial organization of patient P05. **(h)** Stacked bar plots showing cell-type proportions across tumor foci from individual patients. **(i)** Stacked bar plots showing cell-type proportions in adjacent normal (N) verus tumor (T) tissues. **(j)** Comparison of major cell-type proportions between adjacent normal (N) and tumor (T) tissues.

scFAST-seq generated 167,658 high-quality single cells and resolved nine major cell populations, including hepatocytes (Hep), T cells, NK cells, myeloid cells (Mye), B cells, plasma cells (pB), fibroblasts (Fib), endothelial cells (Endo), and proliferating cells (Pro) (**figure 1c,d**). Complementary scATAC-seq profiling of 126,164 nuclei identified eight chromatin-accessibility-defined cell populations, which showed consistent annotation with transcriptomic profiles (**figure 1c,e**). Integrated analyses demonstrated high concordance across samples with minimal batch effects (**online supplemental figure 2a**), supporting the robustness of the multi-omic dataset. Spatial transcriptomic profiling using Stereo-seq (n=14, four chips) further mapped cellular organization within tumor tissues and provided spatial context for heterogeneous tumor ecosystems (**figure 1f**). H&E staining distinguished tumor, adjacent normal, and stromal regions, enabling spatial interpretation of molecular patterns (**figure 1f, online supplemental figure 2b**). Spatial clustering identified 14 distinct cell clusters representing diverse cellular states, including non-malignant hepatocytes (C1, C2, and C7, predominantly from P01_N1/N2 and P02_N1/N2), malignant hepatocytes (C3 mainly from P04_T2 and P05_T1/T3; C4 and C5 primarily from P01_T1/T2; C9 mostly from P05_T2), and stromal/immune-enriched regions (C6) (**figure 1f, online supplemental figure 2b**). Non-malignant hepatocytes exhibited *ALB* expression, whereas malignant hepatocyte regions displayed elevated *GPC3* expression. Immune and stromal regions were characterized by *CD3D*, *CD14*, and *ACTA2* expression, respectively (**online supplemental figure 2c,d**). These spatial assignments were further validated by mIHC (**figure 1g**).

Comparative analysis of cellular composition revealed extensive inter-focal heterogeneity among tumor lesions and matched peritumoral tissues (**figure 1h**). Notably, scATAC-seq showed higher hepatocyte representation than scFAST-seq, likely reflecting differences in cellular fragility and nuclear accessibility during sample processing (**figure 1h, online supplemental figure 2e**). Smaller tumor foci (e.g., P01_T2, P04_T2) exhibited increased T-cell infiltration compared with their corresponding larger lesions (P01_T1, P04_T1) (**figure 1h**), consistent with spatial observations from Stereo-seq (**online supplemental figure 2f**). Furthermore, HBV-associated, recurrent, and advanced-stage tumors displayed increased hepatocyte abundance accompanied by reduced T-cell infiltration, suggesting differential immune remodeling among tumor foci (**online supplemental figure 2g**).

Across the cohort, tumor foci were characterized by expansion of malignant hepatocyte and proliferative populations together with depletion of immune and stromal compartments compared with peritumoral tissues (**figure 1i,j**). This immune-desert phenotype was further supported by independent public single-cell HCC datasets (**online supplemental figure 3a–d**). Collectively, these findings demonstrate marked inter-focal heterogeneity alongside conserved malignant and tumor microenvironmental remodeling across multifocal HCC.

### Malignant hepatocytes exhibit inter-focal heterogeneity with conserved oncogenic programs across tumor foci in multifocal HC

To characterize the evolutionary landscape of malignant hepatocytes and determine whether divergent tumor foci share conserved oncogenic programs in multifocal HCC, we inferred large-scale CNVs from scFAST-seq profiles and resolved hepatocyte populations into six distinct clusters. Cluster C5, predominantly derived from matched peritumoral tissues, displayed a diploid CNV pattern and was classified as normal hepatocytes (Hep_N), whereas clusters C1–C4 and C6 from tumor lesions exhibited extensive CNV alterations and were annotated as malignant hepatocytes (Hep_M) (**figure 2a**). The CNV-based classification was further supported by concordant chromatin accessibility patterns from matched scATAC-seq datasets (**online supplemental figure 4a,b**), confirming the malignant identity of Hep_M populations.

**Figure 2.**
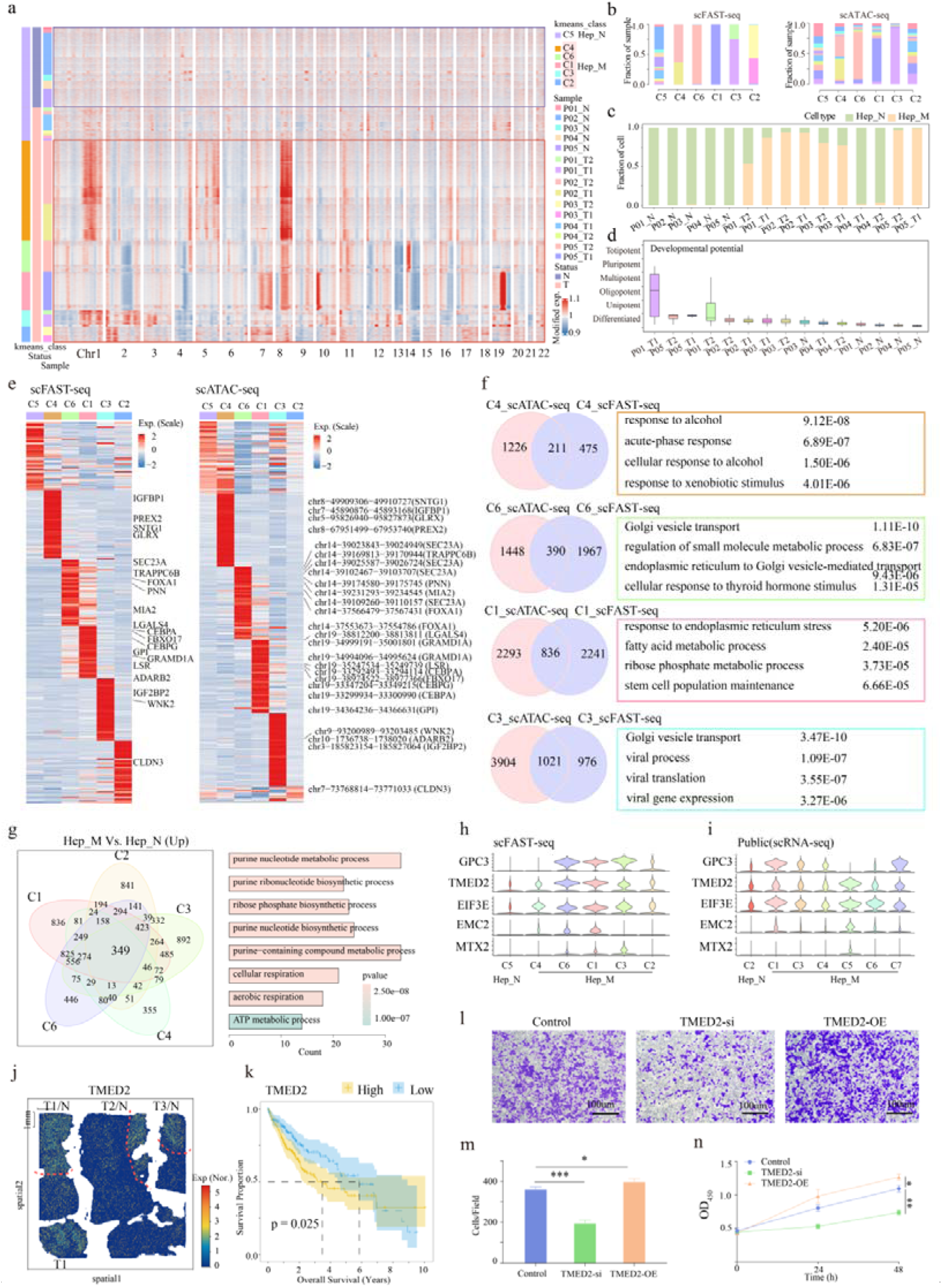
Inter-focal heterogeneity and shared oncogenic programs in malignant hepatocytes of multifocal HCC. **(a)** Heatmap illustrating large-scale CNVs across individual hepatocytes (rows), with red indicating amplifications and blue indicating deletions. Cluster C5 corresponds to normal hepatocytes (Hep_N), while clusters C1-C4, and C6 represent malignant hepatocytes (Hep_M). **(b)** Stacked bar charts showing the distribution of samples within each cluster in scRNA-seq (left) and scATAC-seq (right). **(c)** Stacked bar charts showing the proportions of Hep_M and Hep_N cells. **(d)** Boxplots depicting the differentiation potential of hepatocytes across different samples. **(e)** Heatmaps displaying differentially expressed genes (left) and differentially accessible chromatin peaks (right) across hepatocyte clusters. **(f)** Venn diagram showing the overlap between genes in scFAST-seq and scATAC-seq (peak corresponding genes), alongside functional enrichment analysis of overlapping genes. **(g)** Venn diagram representing the intersection of upregulated genes among Hep_M clusters, with corresponding enrichment analysis of shared genes. **(h,i)** Violin plots showing the expression levels of *GPC3*, *EIF3E*, *EMC2*, *TMED2*, and *MTX2* in scFAST-seq **(h)** and the public scRNA-seq dataset **(i)**. **(j)** Spatial expression patterns of *TMED2*. **(k)** Kaplan-Meier survival curves of *TMED2*. **(l,m)** Transwell assays were performed to determine the effect of *TMED2* overexpression or knockdown on the migration of HepG2 cells. **(n)** CCK-8 assays were performed to examine the effect of *TMED2* overexpression or knockdown on the proliferative ability of HepG2 cells.

Subclonal analysis revealed striking inter-focal segregation of malignant hepatocyte populations. Distinct Hep_M subclusters preferentially occupied different tumor foci, with C4 enriched in P02_T1/T2, C3 in P01_T1/T2, C2 in P03_T1/T2, and C1/C6 in P05_T1/T2, indicating patient- and lesion-specific evolutionary trajectories (**figure 2b**). The abundance of Hep_M populations also varied substantially among tumor foci (**figure 2c**). For example, P02 and P05 lesions exhibited extensive expansion of Hep_M populations, whereas P04 lesions contained relatively limited malignant hepatocyte populations. Within patient P01, the larger T1 lesion showed greater Hep_M expansion than the smaller T2 lesion, further demonstrating heterogeneity in malignant cell burden among anatomically distinct foci.

Despite these divergent cellular compositions, malignant hepatocyte populations exhibited shared functional states associated with aggressive tumor evolution. CytoTRACE^18^ analysis revealed enhanced differentiation potential of Hep_M populations from P01, P02, and P05, suggesting progenitor-like and more aggressive cellular states, whereas Hep_M cells from P04 retained molecular features closer to Hep_N populations (**figure 2d**). These molecular features were consistent with clinical characteristics, as P02 represented recurrent disease, P05 displayed extensive multifocal involvement, and P04 exhibited relatively limited lesions.

Integrated analysis of scFAST-seq and scATAC-seq demonstrated concordant transcriptional and chromatin accessibility programs across Hep_M subclusters (**figure 2e**). Functional enrichment analysis further revealed distinct biological states among malignant hepatocyte subclones (**figure 2f**). Cluster C4 cells showed enrichment of alcohol-associated response pathways, suggesting adaptation to alcohol-related liver injury, whereas C1 and C6 exhibited activation of Golgi vesicle transport and metabolic pathways, reflecting altered intracellular trafficking and metabolic remodeling. Notably, Cluster C3 uniquely displayed enrichment of viral response programs, consistent with HBV-associated tumor evolution.

At the population level, Hep_M cells shared a common malignant transcriptional program characterized by activation of biosynthetic and metabolic pathways and suppression of cell adhesion and immune-associated programs compared with Hep_N cells (C5) (**figure 2g, online supplemental figure 4c,d**). These conserved features were independently validated in public HCC single-cell datasets, which recapitulated the enrichment of malignant hepatocyte signatures and attenuation of adhesion- and immune-related pathways (**online supplemental figure 5**).

To identify core molecular programs underlying malignant hepatocyte states, we integrated multiple datasets and identified four consistently upregulated genes, including *TMED2*, *EIF3E*, *EMC2*, and *MTX2*, together with 18 downregulated genes, including *TAT*, *SLC22A1*, and *CYP2A6* (**figure 2h,i, online supplemental figure 6a**). Spatial transcriptomic analysis further demonstrated heterogeneous but recurrent expression patterns of these genes across tumor foci, linking malignant hepatocyte diversification with conserved molecular programs (**figure 2j, online supplemental figure 6b,c**). Among these candidates, high *TMED2* expression was strongly associated with poor patient survival, and similar prognostic associations were observed for *EIF3E* and *MTX2*, whereas reduced expression of *TAT* and *SLC22A1* correlated with unfavorable outcomes (**figure 2k, online supplemental figure 6d,e**). These prognostic relationships were consistently observed in both our multifocal HCC cohort and TCGA-LIHC dataset (**online supplemental figure 6f**).

Recent studies ^19–22^ have implicated *TMED2* and *EMC2* as oncogenic regulators in multiple cancers, but their roles in HCC remain unclear. Functional perturbation experiments demonstrated that depletion of *TMED2* or *EMC2* impaired HCC cell proliferation and migration, whereas *TMED2* overexpression enhanced malignant phenotypes (**figure 2l–n, online supplemental figure 6g–i**). However, the molecular mechanisms by which TMED2 and EMC2 promote HCC malignant progression remain to be elucidated. Collectively, these findings indicate that malignant hepatocytes across multifocal HCC display substantial phenotypic diversification while retaining conserved oncogenic programmes. We next sought to determine whether these divergent malignant states were shaped by distinct evolutionary trajectories.

### Clonal evolution drives divergent malignant trajectories and progressive adaptive states in multifocal HCC

To determine the evolutionary basis underlying inter-focal heterogeneity in multifocal HCC, we reconstructed the clonal trajectories of malignant hepatocytes across paired tumor foci using large-scale CNV inference (**figure 3a**). This analysis distinguished lesions arising through IM from those originating through MO, revealing distinct evolutionary routes that shape lesion-specific malignant phenotypes. Tumor foci with marked differences in size generally exhibited linear clonal evolution, whereas lesions of comparable size more frequently reflected independent multicentric origins (**figure 3a**). For example, in patient P01, the smaller T2 lesion (1 cm) acquired an early gain of chromosome 2p, whereas the larger T1 lesion (9 cm) retained this alteration while accumulating additional gains on chromosomes 2q and 3p, consistent with sequential clonal expansion (**figure 3a**). Similar stepwise evolutionary trajectories were observed in P02 and P03, supporting intrahepatic metastatic progression. In contrast, P05_T1 and P05_T2 shared only a subset of CNVs while harboring distinct chromosomal alterations (T2: chr2p, chr4q, chr14p; T1: chr3q, chr10p, chr19p, chr19q), consistent with parallel evolution from independent tumor origins (**figure 3a**). Collectively, these findings demonstrate that multifocal HCC evolves through both linear clonal expansion and independent multicentric evolution, generating substantial evolutionary diversity across tumor foci.

**Figure 3.**
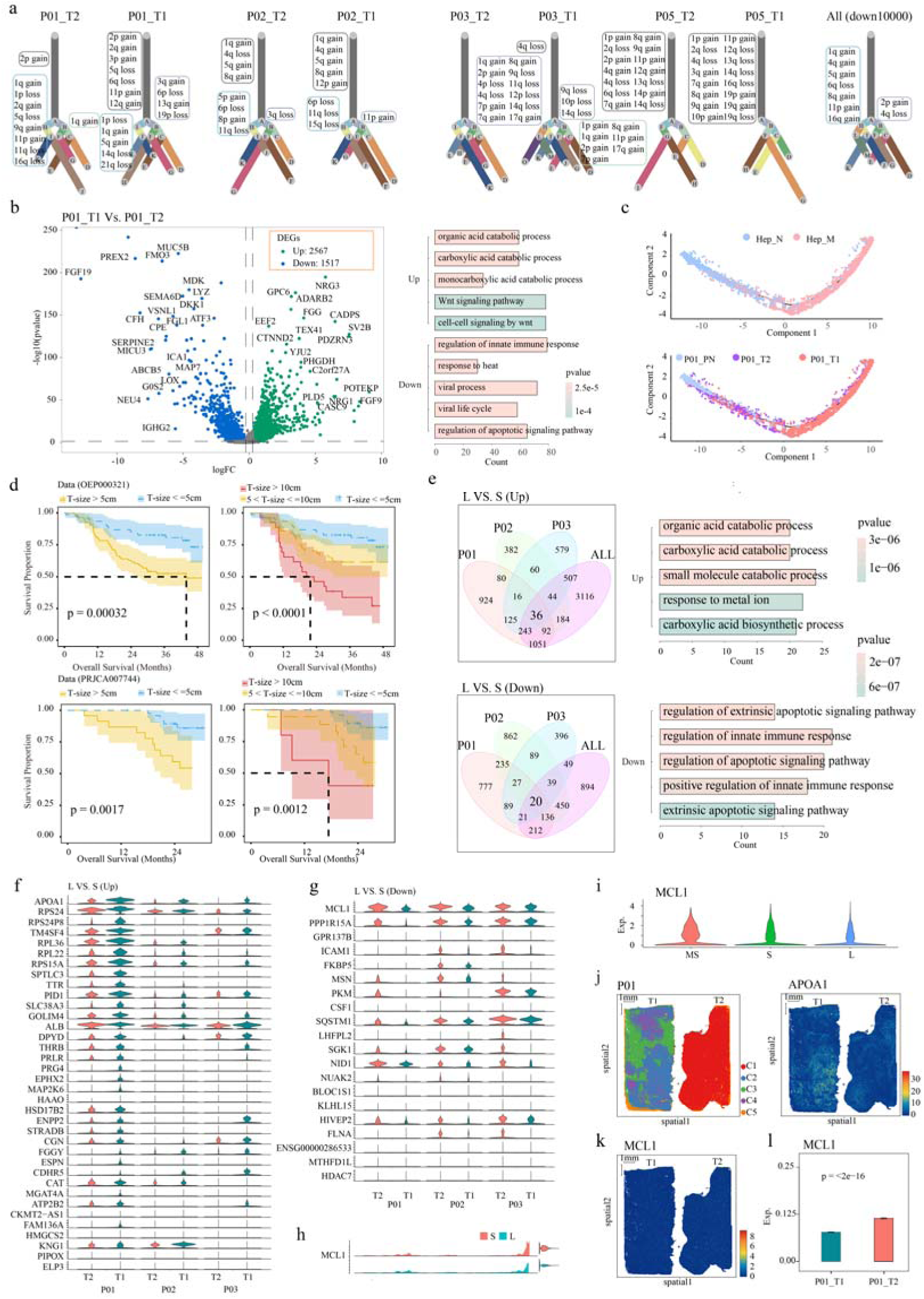
Clonal evolution and size-associated malignant reprogramming across tumor foci in multifocal HCC. **(a)** Clonality trees for each tumor focus, with branch lengths representing the number of cells in each subclone. Chromosomal gains and losses were inferred to reconstruct the clonal evolution. **(b)** Volcano plot showing differentially expressed genes between malignant cells from focus T1 and T2 in patient P01 (left); bar chart summarizing functional enrichment of upregulated and downregulated genes (right). **(c)** Development trajectory of hepatocytes in P01 inferred by monocle2, with cells colored by cell state (top) and sample origin (bottom). **(d)** Tumor focus size was positively correlated with patient survival in both the HRA001748 and OEP000321 datasets. **(e)** Venn diagram illustrating the overlap of differentially expressed genes (left) and corresponding functional enrichment analysis (right). **(f,g)** Violin plots of the upregulated **(f)** and downregulated **(g)** overlap genes. **(h)** Chromatin accessibility peaks of *MCL1* in small and large tumor foci based on scATAC-seq data. **(i)** Expression distribution of *MCL1* across micro-lesions (MS), small lesions (S), and large lesions (L) in the public scRNA-seq dataset. **(j–l)** The spatial expression pattern of *MCL1* in P01_T1 (large lession) and P01_T2 (small lesion) samples was examined using Stereo-seq.

Evolutionary divergence was accompanied by pronounced transcriptional reprogramming of malignant hepatocytes. In P01, the larger T1 lesion exhibited activation of catabolic processes and Wnt signaling together with suppression of immune response, antiviral defense, and apoptotic pathways, indicating progressive metabolic adaptation and immune evasion during tumor evolution (**figure 3b**). Pseudotime analysis further supported linear evolutionary progression, positioning malignant hepatocytes from the smaller P01_T2 lesion at an intermediate differentiation state between normal hepatocytes and the advanced P01_T1 lesion (**figure 3c**). Similar progressive transcriptional trajectories were observed in P02 and P03, where malignant cells from smaller lesions also occupied intermediate developmental states and displayed gradual activation of metabolic programs (**figure 3c, online supplemental figure 7a–d**). In contrast, P05 lesions followed distinct evolutionary paths: P05_T1 preferentially activated DNA replication and cell-cycle programs, whereas P05_T2 exhibited enrichment of apoptosis- and autophagy-related pathways, consistent with independent molecular evolution (**online supplemental figure 7e,f**). These observations indicate that distinct clonal origins give rise to divergent transcriptional states that further amplify inter-focal molecular heterogeneity.

To identify molecular programs associated with evolutionary progression, we compared malignant hepatocytes from larger and smaller tumor foci. Larger lesions consistently exhibited more aggressive molecular features and were associated with poorer clinical outcomes **(figure 3d)**. Malignant hepatocytes from advanced lesions showed enhanced activation of metabolic and biosynthetic programs, accompanied by suppression of apoptosis and innate immune pathways, indicating a progressive transition toward metabolically adaptive and immune-evasive states during tumor evolution **(figure 3e–g)**. These transcriptional changes were supported by concordant alterations in chromatin accessibility identified by scATAC-seq **(online supplemental figure 8a,b)** and independently validated in public HCC single-cell datasets **(online supplemental figure 8c)**.

Integrated transcriptomic and epigenomic analyses further identified 12 consistently upregulated and 10 downregulated genes associated with evolutionary progression, exhibiting coordinated changes in gene expression and chromatin accessibility **(online supplemental figure 8d–f)**. Among these candidates, the anti-apoptotic regulator *MCL1* showed preferential activation in smaller lesions, suggesting an early survival advantage during tumor initiation, whereas *APOA1* displayed the opposite pattern and was progressively reduced in larger tumors, consistent with spatial transcriptomic observations **(figure 3g–l)**. These findings support a stepwise adaptive transition, in which early malignant hepatocytes rely on apoptosis resistance, whereas advanced lesions acquire metabolic plasticity and immune-evasive properties to sustain tumor progression.

Collectively, these findings demonstrate that distinct clonal evolutionary trajectories generate divergent malignant states across tumor foci in multifocal HCC. During tumor progression, these evolutionary trajectories are accompanied by progressive adaptive reprogramming, characterized by metabolic plasticity, lineage diversification, and immune evasion. These observations suggest that multifocal HCC evolution involves both lesion-specific diversification and stepwise acquisition of aggressive malignant traits. Despite this extensive inter-focal molecular divergence, whether evolutionarily distinct tumor foci converge on shared immunosuppressive microenvironments remains unclear.

### T-cell immunosuppressive remodeling and conserved Treg-mediated MHC-I–CD8 interactions associated with CD8+ T-cell dysfunction in multifocal HCC

Given the malignant heterogeneity and evolutionary divergence across tumor foci, we next investigated whether multifocal HCC established a shared immunosuppressive T-cell ecosystem. We first characterized the T-cell landscape using scFAST-seq **(figure 4a**). Unsupervised clustering identified 15 transcriptionally distinct T-cell clusters, which were further annotated into 12 canonical T-cell subsets, including cytotoxic CD8+ T (C1), activated CD8+ T (C11), exhausted CD8+ T (C2), tissue-resident memory CD8+ T (Trm CD8+ T, C13), mucosal-associated invariant T (MAIT, C5), proliferative CD8+ T (Pro CD8+, C6 and C10), regulatory T cells (Tregs, C4), exhausted CD4+ T (C9), helper T (Th, C0), CCR4+ Th2 (C7), and proliferative CD4+ T (Pro CD4+, C14) **(figure 4a–c**). Three minor clusters (C3, C8, C12) lacking definitive lineage markers were designated as "Others". Cell identity was consistently supported by functional enrichment analyses and label transfer from matched scATAC-seq datasets (**figure S9a,b**).

**Figure 4.**
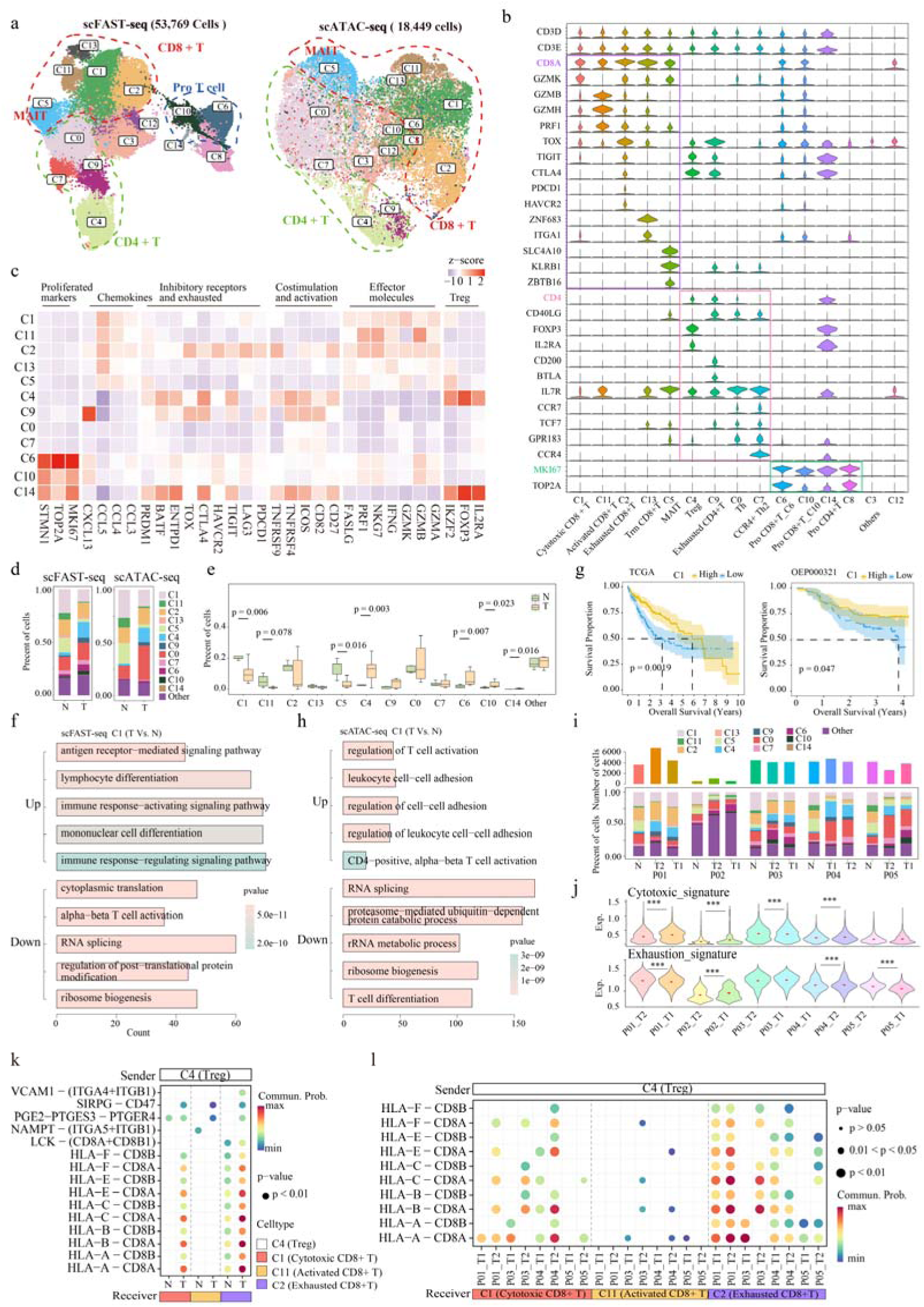
T-cell composition, functional states, and intercellular communication landscapes in multifocal HCC. **(a)** UMAP plots showing T cell lineages identified in scFAST-seq (left) and scATAC-seq (right). **(b)** Violin plots displaying the expression of subtype-specific marker genes across T cell subtypes. **(c)** Heatmap of scaled normalized gene expression within T cell clusters. **(d)** Stacked bar charts showing the proportion of T cell subtype contained in adjacent non-tumor and tumor samples from scFAST-seq (left) and scATAC-seq (right). **(e)** Boxplots showing the variation in T cell subtype proportions between adjacent non-tumor and tumor samples. **(f,h)** Functional enrichment of cytotoxic CD8+ T (C1) cell DEGs in tumor tissues identified by scFAST-seq **(f)** and scATAC-seq **(h)**. **(g)** Kaplan–Meier survival analysis of Cytotoxic CD8+ T cells (C1) in TCGA-LIHC and OEP000321 datasets. **(i)** Stacked bar charts showing T cell subtype composition across different patient samples. **(j)** Cytotoxic and exhaustion-related signature scores across tumor foci, compared using t-tests. **(k,l)** Receptor-ligand pairs showing significant differences between adjacent non-tumor and tumor tissues **(k)**, and across tumor foci **(l),** based on Treg (C4) and CD8+ T cell clusters. Dot size represents the p-value, and color reflects communication probability.

Next, we compared the distribution of T cell subtypes between tumor and adjacent non-tumor tissues (**figure 4d,e)**, and identified differentially expressed genes (**online supplemental figure 9c)**. Notably, cytotoxic CD8+ T cells (C1), activated cytotoxic CD8+ T cells (C11), and MAIT cells (C5) were significantly decreased in tumor tissues (**figure 4d,e**), reflecting impaired cytotoxic T lymphocyte (CTL) infiltration. Interestingly, despite their reduced presence, cytotoxic CD8+ T cells in tumors retained transcriptional activity in immune-related pathways such as lymphocyte differentiation, antigen receptor-mediated signaling, and immune response activation (**figure 4f**), suggesting partial preservation of effector potential. Similarly, residual MAIT cells exhibited immune activation signatures, suggesting that they may retain functional potential (**online supplemental figure 9d)**. Importantly, higher expression of the C1- and C5-associated gene signatures was significantly associated with better patient survival (**figure 4g, online supplemental figure 9e**), underscoring their potential role in orchestrating effective antitumor responses.

In contrast, immunosuppressive and dysfunctional T-cell populations were preferentially expanded in tumor tissues. Tregs (C4) were markedly enriched in tumor lesions, consistent with their role in suppressing antitumor immune responses (**figure 4d,e**). Proliferative CD8+ T cells (C6 and C10) and proliferative CD4+ T cells (C14) were also increased, potentially reflecting chronic antigen stimulation and dysfunctional immune activation (**figure 4d,e**). These alterations were consistently observed at both transcriptional and chromatin accessibility levels in matched scATAC-seq datasets (**figure 4h, online supplemental figure 9f-h**) and were further validated in independent single-cell datasets and bulk RNA-seq deconvolution analyses (**online supplemental figure 10, online supplemental figure 11**). Together, these findings demonstrate a broad remodeling of the T-cell compartment characterized by loss of cytotoxic immune populations and expansion of immunosuppressive or dysfunctional T-cell states in multifocal HCC.

Although the composition and magnitude of T-cell alterations varied among patients and tumor foci (**figure 4i,j, online supplemental figure 9h**), a common immunosuppressive pattern was observed across lesions. For example, P02 exhibited an immune-desert phenotype characterized by near-complete depletion of cytotoxic CD8+ T cells (C1), whereas P01 displayed intra-lesional differences in T-cell states, with the larger lesion (T1) retaining stronger cytotoxic activity and the smaller lesion (T2) showing increased exhaustion signatures. P05 exhibited uniformly reduced T-cell infiltration and limited cytotoxic activity across tumor foci, consistent with advanced multifocal disease (∼8 cm, stage C). These observations indicate that, despite variable immune compositions, multifocal HCC consistently develops an impaired cytotoxic immune environment.

To investigate potential mechanisms underlying T-cell dysfunction, we next analyzed intercellular communication among T-cell subsets (**figure 4k,l**). Tregs (C4) consistently emerged as the dominant immunosuppressive population across tumor lesions and exhibited extensive interactions with CD8+ T-cell subsets. Among these, both exhausted and cytotoxic CD8+ T cells received the strongest Treg-derived suppressive signals, whereas activated CD8+ T cells were comparatively less affected, suggesting a hierarchical program of T-cell suppression (**figure 4k,l**). Although MHC-I molecules (HLA-A/B/C) are classically recognized as mediators of antigen presentation to CD8+ T cells^23^, our findings suggest that tumor-associated Tregs may exploit MHC-I-related interactions to modulate CD8+ T-cell states within the immunosuppressive tumor microenvironment.

Importantly, although the strength of Treg-mediated MHC-I interactions varied among tumor foci, this interaction pattern was consistently observed across multifocal lesions (**figure 4l**). For instance, stronger MHC-I interactions in P01_T2 were accompanied by increased CD8+ T-cell exhaustion compared with P01_T1, whereas uniformly weak signaling in P05 was consistent with its immune-desert phenotype. Collectively, these findings identify a conserved Treg-mediated MHC-I–CD8 interaction module associated with impaired cytotoxic immunity in multifocal HCC, suggesting a potential mechanism by which Tregs contribute to CD8+ T-cell dysfunction across diverse tumor foci.

### SPP1+/SPRED1+ TAMs mediate conserved immunosuppressive interactions with exhausted CD8+ T cells through PD-L1 and CD80/CD86 signaling in multifocal HCC

Beyond T-cell-mediated suppression, we next explored whether myeloid populations represent additional conserved immunosuppressive components across divergent tumor foci. We therefore characterized the myeloid compartment using scFAST-seq. Unsupervised clustering identified 22,171 myeloid cells comprising ten transcriptionally distinct populations (**figure 5a**). Three major macrophage populations were defined, including tissue-resident MARCO+ macrophages (MARCO+ MACs, C6), SPP1+ tumor-associated macrophages (SPP1+ TAMs, C0), and SPRED1+ tumor-associated macrophages (SPRED1+ TAMs, C4), together with monocytes and multiple dendritic-cell subsets, including undifferentiated monocytes (Mono, C1), inactive/exhausted macrophages (C2), conventional dendritic cells (CD1C+ cDC2, C5; CLEC9A+ cDC1, C7), mature DCs (LAMP3+ mDCs, C9), plasmacytoid DCs (CLEC4C+ pDCs, C8), and a low-quality cluster (C3) (**figure 5b**). Cell identities were independently confirmed by matched scATAC-seq profiles (**figure 5a, online supplemental figure 12a**).

**Figure 5.**
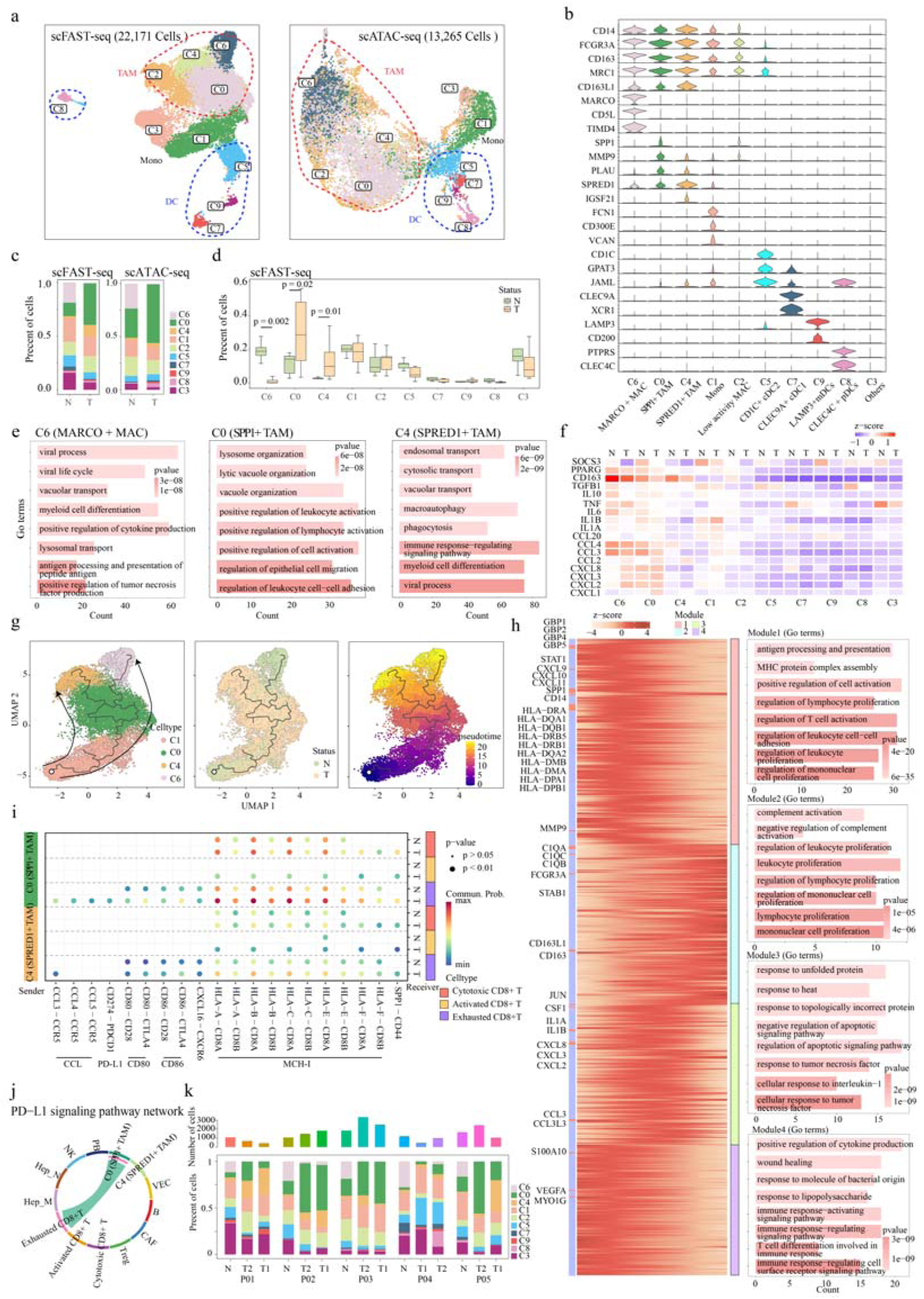
Characterization of myeloid cell states and immunosuppressive interactions across multifocal HCC. **(a)** UMAP plots showing the distribution of Mye cell lineages identified in scFAST-seq (left) and scATAC-seq (right). **(b)** Violin plot illustrating the expression of marker genes across Mye subtypes. **(c)** Stacked bar charts showing the proportions of Mye subtypes in adjacent non-tumor and tumor samples from scFAST-seq (left) and scATAC-seq (right). **(d)** Boxplots comparing the abundance of Mye subtypes between non-tumor and tumor tissues. **(e)** Functional enrichment analysis for clusters MARCO+ MACs (C6), SPP1+ TAMs (C0), and SPRED1+ TAMs (C4). **(f)** Heatmap displaying scaled normalized expression of chemokines, inflammatory genes, and immunosuppressive factors across different Mye subtypes. **(g)** Development trajectory of MARCO+ MACs (C6), SPP1+ TAMs (C0), SPRED1+ TAMs (C4), and undifferentiated monocytes (C1) inferred by monocle3. **(h)** Heatmap showing pseudotime-ordered differentially expressed genes, annotated with representative genes and associated GO terms for each trajectory branch. **(i)** Receptor–ligand interaction analysis between SPP1+ TAMs (C0)/SPRED1+ TAMs (C4) and CD8+ T cells, highlighting key signaling differences between tumor and adjacent non-tumor tissues. **(j)** The inferred PD-L1 signaling network. **(k)** Stacked bar charts illustrating the distribution of Mye subtypes across different tumor foci.

Compared with adjacent tissues, tumor lesions exhibited profound remodeling of the macrophage compartment, characterized by depletion of MARCO+ MACs (C6) and marked expansion of both SPP1+ TAMs (C0) and SPRED1+ TAMs (C4) (**figure 5 c,d, online supplemental figure 12b**). MARCO+ MACs (C6) retained transcriptional programs associated with antiviral responses and myeloid differentiation(**figure 5e and figure S12c**), consistent with homeostatic Kupffer-cell functions, acting as sentinels to maintain tissue homeostasis^24^. In contrast, SPP1+ TAMs (C0) displayed inflammatory and immunoregulatory features, including activation of lysosomal organization, leukocyte activation, and cell-adhesion pathways together with high expression of inflammatory chemokines, IL1A/B, IL10, and TGFB1 (**figure 5e, f, online supplemental figure 12c**). SPRED1+ TAMs (C4) preferentially activated endosomal trafficking and autophagy programs, suggesting complementary roles in stromal remodeling rather than inflammatory activation (**figure 5e, online supplemental figure 12c**). Similar macrophage remodeling was consistently observed in independent HCC scRNA-seq cohorts (**online supplemental figure 13**).

Trajectory reconstruction further demonstrated that tumor-associated macrophages arise through active reprogramming rather than selective expansion. In adjacent tissues, Mono (C1) differentiated predominantly toward homeostatic MARCO+ MACs (C6) through an intermediate SPP1+ state. In contrast, within tumors, Mono (C1) differentiation was redirected toward suppressive SPRED1+ TAMs through SPP1+ intermediates, indicating a tumor-driven macrophage fate switch **(figure 5g)**. Pseudotime-dependent gene module analysis further supported progressive acquisition of suppressive macrophage programs during differentiation **(figure 5h).**

We next investigated whether macrophage heterogeneity converges on conserved immunosuppressive communication programs. Despite marked differences in macrophage composition across lesions, SPP1+ and SPRED1+ TAMs consistently emerged as the dominant suppressive populations. Cell-cell communication analysis revealed extensive interactions between SPP1+/SPRED1+ TAMs and CD8+ T cells mediated by PD-L1 (CD274), CD80/CD86, MHC-I, and CCL signaling pathways, with substantially stronger signaling in tumor than adjacent tissues across patients and tumor foci (**figure 5i, online supplemental figure 14a-c**). Exhausted CD8+ T cells represented the principal recipients of TAM-derived suppressive signals, followed by cytotoxic CD8+ T cells, indicating preferential reinforcement of T-cell dysfunction during tumor evolution (**figure 5i**). Notably, SPP1+ TAMs exhibited particularly strong engagement of exhausted CD8+ T cells through the CD274–PDCD1 and CD80/CD86–CTLA4/CD28 axes (**figure 5j, online supplemental figure 14a,c**), highlighting these cells as major mediators of immune evasion and potential determinants of immunotherapy resistance. MHC-I-dependent interactions closely mirrored those observed in Tregs (**figure 4k,l**), suggesting that distinct suppressive cell populations converge on a common immunoregulatory program.

Although TAM abundance and signaling intensity varied considerably among individual lesions, the overall suppressive communication architecture remained highly conserved (**figure 5k, online supplemental figure 14b,c**). For example, P03 exhibited the highest myeloid infiltration (**figure 1h**), whereas SPP1+ TAMs predominated in P01_T2 and P05_T2 and SPRED1+ TAMs were preferentially enriched in the corresponding T1 lesions (**figure 5k**). These compositional differences were accompanied by quantitative variation in macrophage–T-cell signaling strength but did not alter the core suppressive network **(online supplemental figure 14c)**. Collectively, these findings demonstrate that SPP1+/SPRED1+ TAMs constitute a conserved immunosuppressive hub that transcends profound inter-focal heterogeneity, engaging cytotoxic and exhausted CD8+ T cells through PD-L1, CD80/CD86, and MHC-I signaling. Together with the Treg-mediated suppressive program identified above, these results establish that genetically and evolutionarily distinct tumor foci converge on a shared immunosuppressive microenvironments.

### Convergent expansion of THY1+ CAFs and COL15A1+ VECs accompanies inter-focal heterogeneity in stromal remodeling and angiogenesis in multifocal HCC

Stromal cells, including fibroblasts and endothelial cells, are key regulators of immune modulation, extracellular matrix (ECM) remodeling, and angiogenesis in the HCC microenvironment^25^. Analysis of 14,368 fibroblast (Fib) and endothelial (Endo) cells identified 12 transcriptionally distinct clusters (**figure 6a**). Fibroblast populations comprised THY1+ CAFs (C2), SPP1+ CAFs (C9), MYOCD+ CAFs (C7), and hepatic stellate cells (HSCs; C4), whereas endothelial cells included COL15A1+ (C6), SELE+ (C5), and SEMA3G+ (C3) vascular endothelial cells (VECs), together with CRHBP+ (C1/C8) and PROX1+ (C11) lymphatic endothelial cells (LECs) (**figure 6b**). Cluster C10 co-expressed fibroblast and endothelial markers, suggesting a transitional stromal population. Cell identities were independently confirmed by matched scATAC-seq data (**figure 6a, online supplemental figure 15a**).

**Figure 6.**
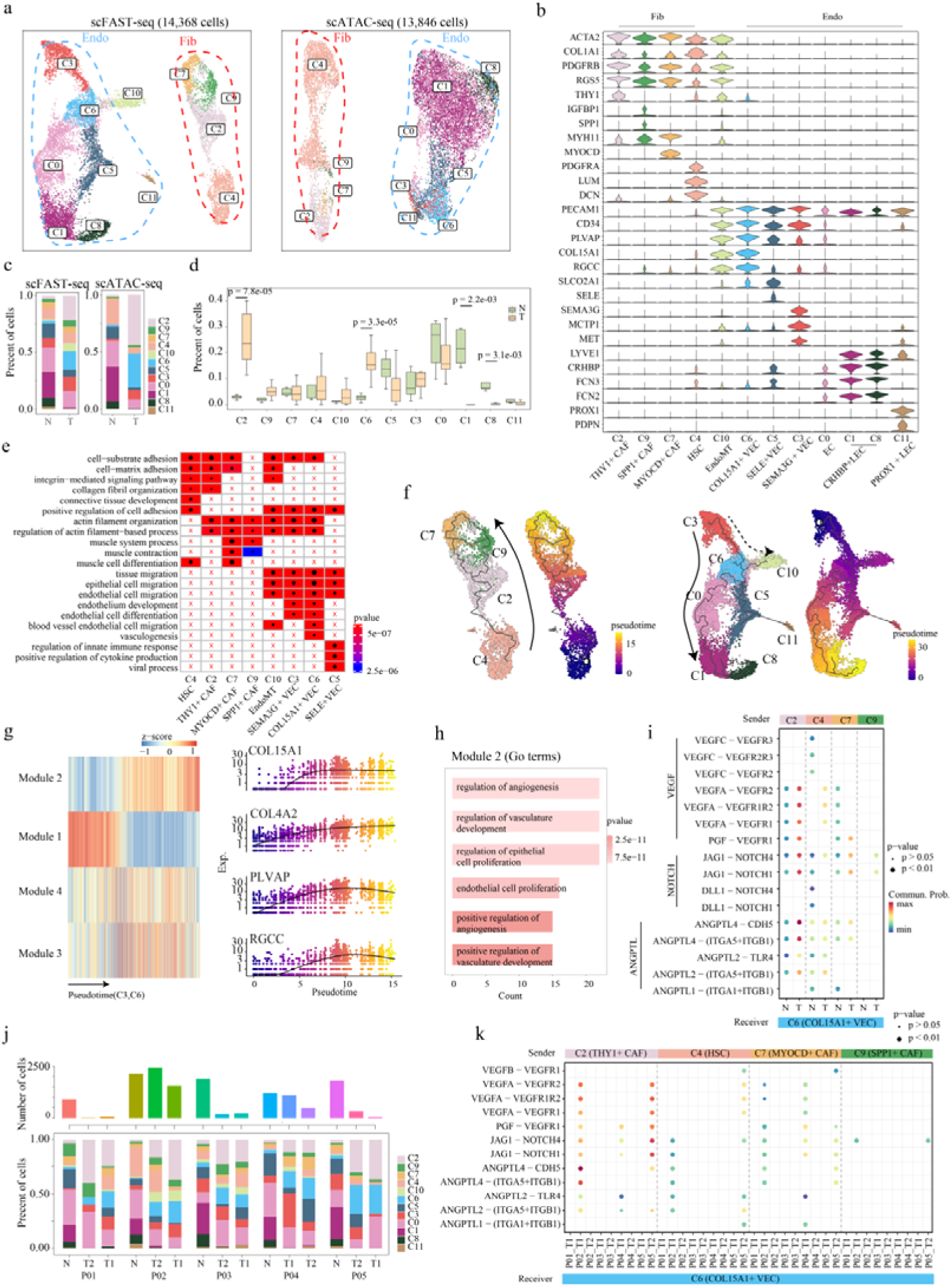
Fibroblast and endothelial cell states and stromal–vascular interactions across multifocal HCC. **(a)** UMAP plots displaying the distribution of fibroblast (Fib) and endothelial (Endo) cell lineages in scFAST-seq (left) and scATAC-seq (right). **(b)** Violin plots showing the expression of representative marker genes across Fib and Endo subtypes. **(c)** Stacked bar charts illustrating the proportions of Fib and Endo subtypes in adjacent non-tumor and tumor tissues, based on scFAST-seq (left) and scATAC-seq (right). **(d)** Boxplots comparing the abundance of Fib and Endo subtypes between adjacent non-tumor and tumor samples. **(e)** Functional enrichment analysis of identified cell clusters. “x” indicates no significant enrichment, and dot size reflects gene count. **(f)** Developmental trajectories of Fib (left) and Endo (right) cells inferred by Monocle3. **(g)** Heatmap showing differentially expressed genes modules arranged in pseudo temporal patterns (left). The expression dynamics of *COL15A1*, *COL4A2*, *PLVAP*, and *RGCC* in module 2 are highlighted. **(h)** Functional enrichment of genes in module 2. **(i)** Receptor– ligand interaction analysis between Fib subtypes and COL15A1+ VEC (C6), highlighting key signaling differences between tumor and adjacent non-tumor tissues. **(j)** Bar plots showing the distribution of fibroblast and endothelial subtypes across patients and lesions. **(k)** Receptor–ligand interaction analysis between Fib subtypes and COL15A1+ VEC (C6), highlighting key signaling differences between lesions.

Compared with adjacent tissues, tumors consistently exhibited expansion of THY1+ CAFs (C2) and COL15A1+ VECs (C6), accompanied by depletion of CRHBP+ LECs (C1, C8) (**figure 6c,d, online supplemental figure 15b–g**). THY1+ CAFs (C2) preferentially activated cell–matrix adhesion, actin cytoskeleton organization, and vesicle transport programs, consistent with active ECM remodeling (**figure 6e, online supplemental figure 16a**). In parallel, COL15A1+ VECs (C6) displayed proliferative and angiogenic signatures characteristic of immature endothelial cells (**figure 6e, online supplemental figure 16b**), whereas loss of CRHBP+ LECs (C1 and C8) suggested impaired lymphatic function and immune cell trafficking (**online supplemental figure 16c**). These findings indicate a conserved stromal shift toward matrix remodeling and angiogenesis across multifocal HCC.

Trajectory analysis further revealed coordinated stromal reprogramming during tumor progression. Fibroblasts differentiated from HSCs through THY1+ CAFs (C2) before diverging into SPP1+ CAFs (C9) and MYOCD+ CAFs (C7), consistent with progressive ECM deposition and stromal activation **(figure 6f)**. Endothelial cells followed a developmental trajectory from SEMA3G+ VECs (C3) to immature COL15A1+ VECs (C6) and subsequently to mature SELE+ VECs (C5), with a pseudotime-associated gene module (including *COL15A1*, *COL4A2*, *PLVAP*, and *RGCC*) supporting vascular remodeling and angiogenesis **(figure 6g,h)**.

Cell–cell communication analysis demonstrated that THY1+ CAFs (C2) preferentially interacted with COL15A1+ VECs (C6) through ANGPTL, NOTCH, and VEGF signaling pathways, indicating coordinated regulation of stromal remodeling and neovascularization (**figure 6i**). Although CAF and VEC composition, together with signaling intensity, varied substantially among patients and tumor foci (**figure 6j,k, online supplemental figure 16d**), the expansion of THY1+ CAFs and COL15A1+ VECs was consistently observed across lesions. These findings identify a conserved stromal remodeling program characterized by coordinated fibroblast–endothelial interactions, while highlighting pronounced inter-focal heterogeneity in angiogenic signaling that may contribute to lesion-specific progression and therapeutic responses.

### Malignant hepatocytes exhibit conserved C3/C5 complement signaling with SPP1+/SPRED1+ TAMs across tumor foci in multifocal HCC

During multifocal HCC progression, effector immune and stromal populations progressively declined, whereas immunosuppressive and tumor-promoting cell populations expanded (**online supplemental figure 17a**). These trends were consistently reproduced in scATAC-seq and independent public scRNA-seq datasets despite marked inter- and intra-patient heterogeneity (**online supplemental figure 17a,b**). For example, P01 retained relatively T cell-rich lesions, whereas P02 was dominated by malignant hepatocytes and stromal cells, illustrating the diverse immune landscapes across multifocal lesions.

Having established heterogeneous cellular states accompanied by recurrent remodeling patterns across malignant, immune, and stromal compartments, we next investigated how these distinct populations communicate to shape conserved immunosuppressive ecosystems. We systematically mapped intercellular communication networks across tumor foci. Among all cell populations, Hep_M emerged as a dominant signaling hub, exhibiting extensive interactions with CD8+ T cells, SPP1+/SPRED1+ TAMs, and CAFs (**online supplemental figure 18a,b**). A tumor-enriched, non-classical PVR/NECTIN2 immune checkpoint axis was activated in Hep_M cells, engaging TIGIT on exhausted CD8+ T cells and Tregs and CD226 on activated CD8+ T cells. This signaling program displayed marked inter-patient variation, with enhanced activation primarily observed in tumor foci from P01 and P05 (**figure 7a,b, online supplemental figure 19a–c**).

**Figure 7.**
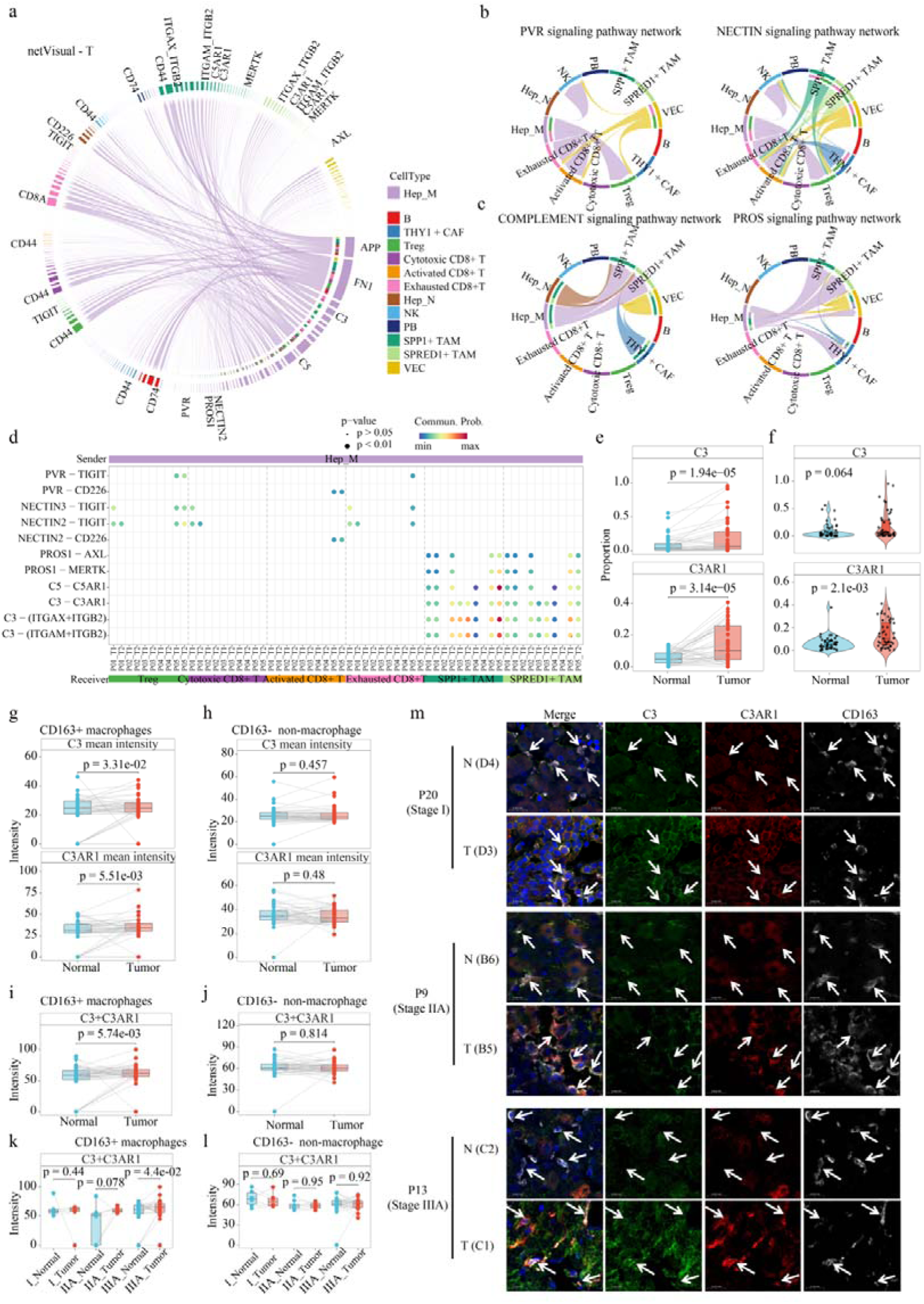
Malignant hepatocyte-centered signaling networks and macrophage-specific complement activation in multifocal HCC. **(a)** Chord diagram showing ligand–receptor interactions through which Hep_M cells communicate with other cell populations in tumor tissues. **(b,c)** Chord diagrams illustrating interactions between Hep_M cells and CD8+ T cells or TAMs mediated by PVR and NECTIN2 **(b)**, and C3/C5 COMPLEMENT and PROS **(c)** signaling pathways. **(d)** Bubble plot showing interaction strength of ligand–receptor pairs associated with PVR, NECTIN2, COMPLEMENT, and PROS signaling across tumor foci. **(e,f)** Quantification of C3- and C3AR1-positive cells in tumor and adjacent tissues. **(g,h)** Expression intensity of C3 and C3AR1 in CD163+ macrophages and CD163−non-macrophage populations. **(i,j)** Quantification of C3−C3AR1 signaling intensity in macrophage and non-macrophage compartments. **(k,l)** Stage-stratified analysis of C3–C3AR1 activation in CD163+ macrophages and CD163− populations. **(m)** Representative mIHC images showing C3 and C3AR1 expression in CD163+ macrophages across representative multifocal HCC cases. N: Adjacent normal, T: tumor, Scale bars, 20 um.

In contrast, Hep_M consistently engaged SPP1+/SPRED1+ TAMs through COMPLEMENT signaling (C3–C3AR1/ITGAX–ITGB2/ITGAM–ITGB2 and C5–C5AR1) and PROS signaling (PROS1–AXL/MERTK) across most patients and tumor foci, whereas these interactions were absent in Hep_N (**figure 7c,d, online supplemental figure 19a,c,d**). This finding extends previous evidence implicating C3/C5 complement signaling in M2-like macrophage polarization and suppression of antitumor T cell responses^26, 27^, and further reveals a tumor cell–derived complement mechanism whereby malignant hepatocytes drive SPP1+/SPRED1+ TAM reprogramming and establish a conserved immunosuppressive microenvironments across heterogeneous tumor foci. These interactions were independently validated by scATAC-seq, public scRNA-seq datasets, and spatial transcriptomics (**online supplemental figure 20a–d, online supplemental figure 21a–c**), demonstrating that, despite profound inter-focal heterogeneity, malignant hepatocytes converge on a shared complement-dependent program to remodel macrophage function and promote immune suppression.

To validate the Hep_M–TAM complement axis identified by single-cell and spatial analyses, we performed mIHC on a tissue microarray containing 48 paired multifocal HCC tumors and matched adjacent tissues (**online supplemental figure 22a**). Compared with adjacent tissues, tumor tissues exhibited significantly increased proportions of C3+ and C3AR1+ cells (**figure 7e,f**), whereas CD163+ macrophages were markedly reduced (**online supplemental figure 22b**). Importantly, elevated C3 and C3AR1 expression intensity was selectively observed within CD163+ macrophages in tumor tissues (**figure 7g)**, while CD163-non-macrophage populations showed no significant differences between tumor and adjacent tissues (**figure 7h**). Moreover, tumor-specific activation of C3–C3AR1 signaling in CD163+ macrophages was consistently observed across different clinical stages of multifocal HCC but was absent in adjacent tissues (**figure 7i–m, online supplemental figure 22c,d**), independent of patient age or sex (**online supplemental figure 22e,f**).

Together, these findings demonstrate that malignant hepatocytes activate conserved immunoregulatory programs, including the C3/C5 complement signaling, to reprogram macrophage states across evolutionarily divergent tumor foci. Large-scale mIHC validation further confirms tumor-specific activation of the C3–C3AR1 axis as a representative mechanism underlying TAM reprogramming. Thus, multifocal HCC exhibits a heterogeneity– convergence pattern, whereby diverse malignant lesions converge on shared macrophage-centered immunosuppressive microenvironments.

### A TMED2–ADAR1 RNA editing axis regulates tumor-intrinsic C3/C5 complement signaling in multifocal HCC

Previous studies have implicated a *NFAT1* (*NFATC2*)-dependent C3a/C3aR– Ca²⁺ feedback loop in TAM polarization^26^. However, *NFATC2* was downregulated in Hep_M cells and showed no differential expression in TAMs between tumor and adjacent tissues (**Figure 8a**), suggesting that the activated Hep_M–TAM C3/C3AR1 axis in multifocal HCC is driven by NFATC2-independent mechanisms.

**Figure 8.**
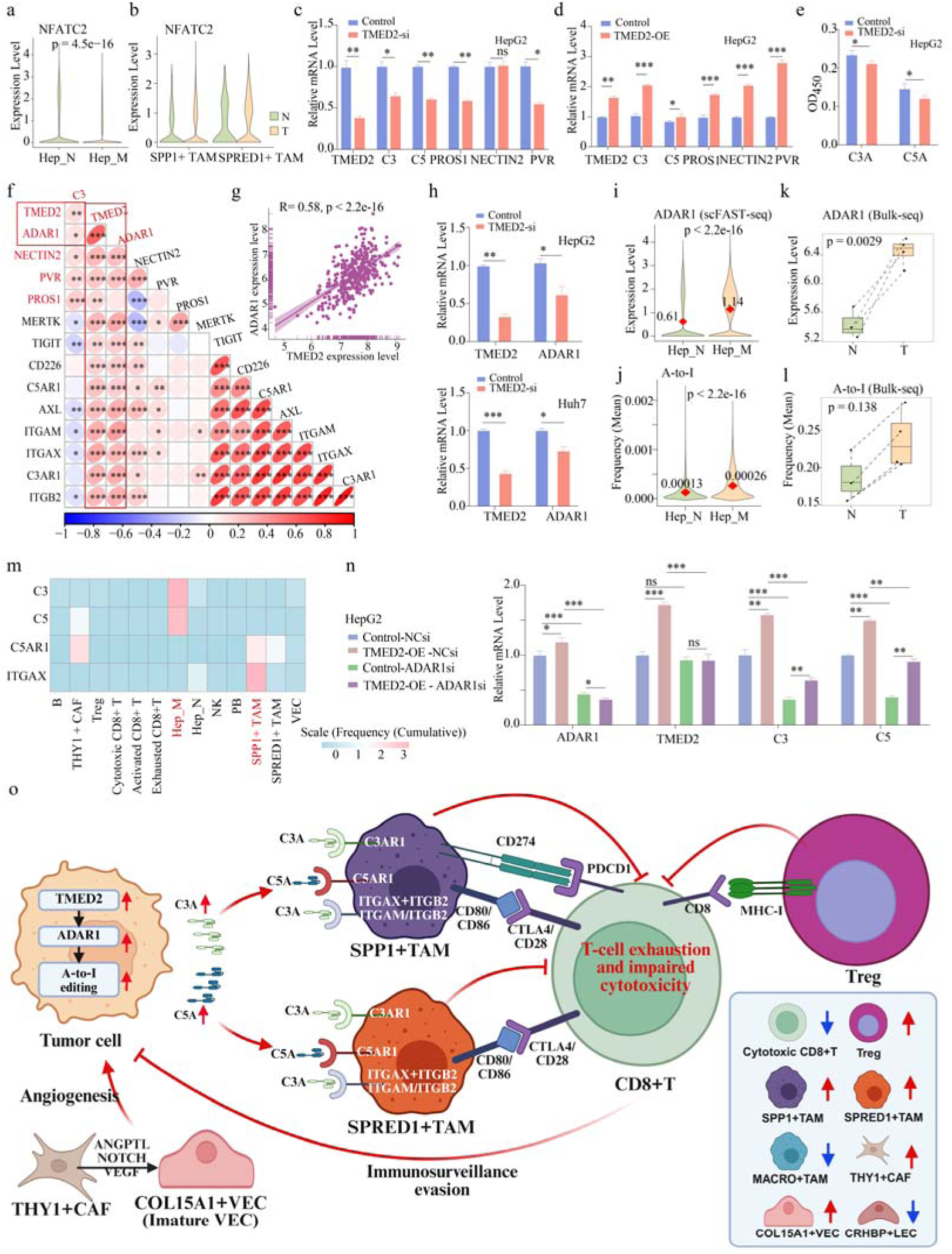
TMED2–ADAR1-mediated RNA editing shapes C3/C5 complement signaling axis linked to immunosuppressive TAMs in multifocal HCC. **(a,b)** Expression of *NFATC2* in malignant hepatocytes (Hep_M) and normal hepatocytes (Hep_N) **(a)**, and in TAM subsets **(b)** from multifocal HCC. **(c,d)** qPCR detecting the Hep_M–TAM communication molecules after *TMED2* knockdown **(c)** or overexpression **(d)** in HepG2 cells. **(d)** Secreted C3A/C5A following *TMED2* knockdown. **(f,g)** Correlation analysis of *TMED2* expression with *ADAR1* and Hep_M–TAM communication molecules in the TCGA-LIHC cohort. (h) *ADAR1* mRNA changes after *TMED2* knockdown in HepG2/Huh7 cells. (i-l) *ADAR1* expression and global A-to-I editing in Hep_M vs. Hep_N cells **(i,j)** and multifocal HCC vs. normal tissues **(k,l)**. **(m)** Heatmap of A-to-I editing frequencies across cell types. **(n)** Rescue experiments demonstrating that *ADAR1* depletion attenuates TMED2-induced activation of C3/C5 expression. **(o)** Proposed model illustrating the TMED2–ADAR1–complement axis in malignant hepatocytes and its role in TAM reprogramming and immune suppression in multifocal HCC.

Given that *TMED2* represents a conserved malignant programme across multifocal HCC lesions and is associated with poor survival (**Figure 2h–n**), we investigated whether it mediates tumour–immune communication. A pan-cancer analysis recently linked *TMED2* to immune features and prognosis^28^, although its immunoregulatory mechanisms remain unclear. *TMED2* knockdown in HepG2 cells significantly reduced the expression of immune-associated mediators, including *C3*, *C5*, *PROS1*, *PVR*, and *NECTIN2*, whereas *TMED2* overexpression restored their expression (**Figure 8b,c**). Consistently, depletion of *TMED2* markedly decreased C3a and C5a secretion (**Figure 8d**), identifying *TMED2* as a tumour-intrinsic regulator of complement activation and broader tumour–immune communication programmes.

Given the emerging role of A-to-I RNA editing in shaping HCC immunity^29^, we next examined whether *TMED2* regulates ADAR1-mediated RNA editing. TMED2 expression positively correlated with *ADAR1* levels in TCGA-LIHC (**figure 8e,f**), and *TMED2* knockdown reduced *ADAR1* expression (**figure 8g**). Both Hep_M cells and tumor tissues exhibited elevated *ADAR1* expression and increased global A-to-I RNA editing, with preferential editing events identified in *C3* and *C5* transcripts, whereas *C5AR1* and *ITGAX* editing occurred predominantly in TAMs (**figure 8h–m, online supplemental figure 23a–e)**. In contrast, no significant RNA editing was detected in *PROS1*, *NECTIN2*, or *PVR*. Importantly, *ADAR1* depletion abolished TMED2-induced C3/C5 upregulation, demonstrating that *TMED2* regulates complement activation through an ADAR1-dependent mechanism (**figure 8n**).

Collectively, these findings define a Hep_M-centered immunoregulatory circuit in multifocal HCC, in which *TMED2* activates ADAR1-mediated A-to-I RNA editing to enhance C3/C5 complement signaling, thereby reprogramming TAMs toward an immunosuppressive state and establishing a conserved immune microenvironments across heterogeneous tumor foci (**figure 8o**). The TMED2–ADAR1 axis therefore represents a potential therapeutic vulnerability for disrupting immune evasion in multifocal HCC.

## DISCUSSION

Multifocal HCC provides a unique opportunity to investigate how divergent malignant evolution shapes multicellular tumor ecosystems. Previous studies have demonstrated substantial molecular differences among spatially separated tumor foci, indicating that distinct evolutionary histories generate heterogeneous malignant states^8, 12^. However, whether such malignant divergence is accompanied by equally divergent tumour ecosystems remains unclear. By integrating single-cell and spatial multi-omics with mIHC and functional validation, we demonstrate that extensive inter-focal heterogeneity coexists with shared oncogenic programmes and a conserved immunosuppressive ecosystem. Although tumour foci acquired distinct genomic, transcriptional, epigenetic, and evolutionary states, they recurrently retained shared malignant programmes, exemplified by *TMED2*, and established similar immune, myeloid, stromal, and vascular states. These findings indicate that malignant evolution and tumour ecosystem organisation are not strictly coupled, suggesting that conserved ecological pressures may impose shared functional dependencies despite tumour-cell diversification.

Cancer evolution has traditionally been viewed as a process driven by genetic diversification, clonal selection, and acquisition of advantageous malignant traits^8, 13, 30^. Under this framework, molecularly distinct tumour populations would be expected to generate corresponding phenotypic and ecological differences. Our findings instead reveal that divergent malignant states can retain shared oncogenic programmes while independently establishing conserved tumour-supportive ecosystems. Spatially separated lesions displayed differences in chromatin accessibility, metabolic states, differentiation trajectories, and clonal evolution, yet repeatedly converged on immune suppression and microenvironmental remodelling. Thus, tumour evolution appears to involve both diversification and conservation, with heterogeneous malignant states retaining selected programmes that facilitate adaptation to a shared ecological context.

A defining feature of this conserved ecosystem was the recurrent establishment of immunosuppressive immune states across evolutionarily distinct tumour foci. Despite variation in T-cell composition among patients and lesions, tumour foci consistently exhibited Treg enrichment, reduced CD8+ T-cell infiltration and cytotoxicity, and recurrent interactions among Tregs, TAMs, and CD8+ T cells. SPP1+/SPRED1+ TAMs and Tregs formed complementary suppressive networks involving CD274–PDCD1, CD80/CD86–CTLA4/CD28, and MHC-I–CD8 interactions. Notably, MHC-I-associated interactions may extend beyond antigen presentation to participate in broader Treg–CD8+ T-cell regulatory networks. These findings suggest that immune suppression represents a conserved functional state rather than a fixed cellular composition, enabling distinct tumour foci to achieve similar impairment of antitumour immunity through shared communication programmes.

Conservation was also observed in the myeloid and stromal compartments. Although macrophage proportions varied among lesions, SPP1+ /SPRED1+ TAMs repeatedly emerged across evolutionarily distinct tumour foci and exhibited extensive communication with malignant hepatocytes and T cells. Their recurrent presence suggests that TAM states are shaped not only by cellular origin but also by tumour-derived ecological signals, enabling functional adaptation while maintaining immunosuppressive properties. Similarly, THY1+ CAFs and COL15A1+ VECs were consistently enriched across tumour foci, despite differences in their activating signals. In addition, Hep_M populations also communicated with VECs and CAFs through EPHA, TRAIL, and AGT signalling pathways, potentially contributing to angiogenesis, apoptosis regulation, and extracellular matrix remodelling (**online supplemental figure 24**). These findings indicate that distinct malignant states can engage diverse molecular routes while converging on conserved fibrovascular architectures that support tumour growth and immune suppression.

A central question arising from these observations is how evolutionarily distinct malignant populations establish similar tumour ecosystems. Our data identify malignant hepatocytes as active regulators of this process and reveal *TMED2* as a shared oncogenic programme linked to conserved tumour– macrophage communication. Integrated cell–cell communication analyses and functional experiments identified a tumour-intrinsic TMED2–ADAR1-dependent C3/C5 complement regulatory axis connecting malignant hepatocytes with SPP1+/SPRED1+ TAMs. This mechanism provides a potential explanation for how divergent tumour foci repeatedly establish similar macrophage-associated immunosuppressive niches and suggests that shared tumour-intrinsic programmes may act as ecological organisers across heterogeneous lesions.

Complement signalling has emerged as an important regulator of tumour immunity by shaping macrophage activation and inflammatory states^31, 32^. However, previous studies have mainly focused on macrophage-intrinsic or immune-cell-derived complement regulation^26, 27, 33, 34^. Our findings identify a tumour-originated complement regulatory mechanism in multifocal HCC, in which malignant hepatocytes acquire complement-producing capacity through a TMED2–ADAR1-dependent RNA editing programme. A-to-I RNA editing represents an epitranscriptomic layer linking shared malignant programmes to C3/C5 expression and secretion, thereby enabling tumour cells to directly influence macrophage states. These findings extend current understanding of complement regulation from an immune-cell-centred model toward a tumour-initiated intercellular communication mechanism and establish the TMED2– ADAR1–C3/C5 axis as a mechanistic link between malignant programmes and conserved tumour–macrophage interactions.

The identification of shared oncogenic programmes and conserved immunosuppressive ecosystems has important therapeutic implications. Current immunotherapies, including immune checkpoint blockade, remain limited by complex and heterogeneous immunosuppressive networks in HCC^35, 36^. Our findings suggest that targeting lesion-specific genetic alterations alone may not fully address the shared ecological dependencies sustaining multifocal disease. Instead, targeting conserved tumour–immune interactions may provide a complementary strategy across genetically distinct tumour foci. The recurrent activation of tumour-derived complement-associated macrophage remodelling raises the possibility that disrupting the TMED2–ADAR1–C3/C5 axis could interfere with cross-focal tumour– macrophage dependencies and enhance antitumour immunity.

Several limitations should be acknowledged. First, although integrated single-cell and spatial multi-omics, mIHC validation, and functional experiments support the involvement of the TMED2–ADAR1–C3/C5 axis in tumour–macrophage communication, its therapeutic relevance requires further validation *in vivo* using multifocal HCC models and patient-derived systems. Second, the upstream mechanism by which *TMED2* regulates *ADAR1* and the specific A-to-I RNA editing events through which ADAR1 controls *C3/C5* expression remain unclear. Given that *TMED2* is a Golgi– endoplasmic reticulum protein trafficking regulator^37^, future studies integrating trafficking analyses, *ADAR1* regulatory profiling, and site-specific RNA editing approaches will be required to define the complete TMED2–ADAR1–C3/C5 axis. Third, larger longitudinal cohorts are required to determine how shared oncogenic programmes and conserved tumour ecosystems evolve during disease progression and therapeutic selection.

In summary, our study demonstrates that malignant heterogeneity does not necessarily translate into functional heterogeneity of the tumour ecosystem in multifocal HCC. Spatially separated tumour foci undergo extensive genomic, transcriptional, epigenetic, and evolutionary diversification while retaining shared oncogenic programmes and repeatedly establishing conserved immunosuppressive ecosystems. The TMED2–ADAR1–C3/C5 complement axis provides a mechanistic link between shared malignant programmes and conserved tumour–macrophage interactions, revealing how divergent tumour foci achieve functional convergence. These findings establish a multi-omic and spatial framework for distinguishing lesion-specific malignant states from shared functional dependencies and highlight the TMED2–ADAR1–C3/C5 axis as a potential cross-focal therapeutic vulnerability in multifocal HCC.

## Contributors

W.L.Z. conceived and supervised the study. W.L.Z. and C.W. designed the study. C.W. analyzed the data. C.W., S.M., and K.S. prepared materials for scFAST-seq, scATAC-seq, Stereo-seq, and bulk RNA-seq. K.S. performed the experiments for validation. C.W., Z.H.J., D.Y.L., and H.P.Z. designed and constructed the website. C.W., W.L.Z. interpreted data. S.M., N.N.S., and W.L.L curated the clinical data. C.W. and W.L.Z. wrote and revise the manuscript. All authors read and approved the manuscript.

## Patient consent for publication

Not applicable

## Ethics approval

All patients provided written informed consent, and the study was approved by the Ethics Committee of Guangzhou Medical University (approval no. 202405020).

## Data availability statement

Raw scFAST-seq, scATAC-seq, single-cell spatial transcriptomic, and in-house bulk RNA-seq datasets have been deposited in the GSA database (accession numbers: HRA015013, HRA015779, HRA016973, and HRA017034, respectively). Processed scFAST-seq and scATAC-seq expression matrices are available via GEO (GSE205333 and GSE324115). Public scRNA-seq data HRA001748^16^ were downloaded from the Genome Sequence Archive (https://ngdc.cncb.ac.cn/gsa-human/browse/HRA001748). the full dataset can be explored interactively at http://www.biomedical-web.com/hccOmics/. Analysis code is available at GitHub (https://github.com/Wucheng-tech/HCC). Public bulk RNA-seq datasets included TCGA-LIHC (UCSC Xena, https://xena.ucsc.edu/) and OEP000321 (HCCDB v2.0^17^).

## References

[1] Singal AG, Kanwal F, Llovet JM. Global trends in hepatocellular carcinoma epidemiology: implications for screening, prevention and therapy. Nat Rev Clin Oncol 2023;20:864–84.

[2] Roayaie S, Jibara G, Tabrizian P, et al. The role of hepatic resection in the treatment of hepatocellular cancer. Hepatology 2015;62:440–51.

[3] Fukami Y, Kaneoka Y, Maeda A, et al. Liver Resection for Multiple Hepatocellular Carcinomas: A Japanese Nationwide Survey. Ann Surg 2020;272:145–54.

[4] Singal AG, Llovet JM, Yarchoan M, et al. AASLD Practice Guidance on prevention, diagnosis, and treatment of hepatocellular carcinoma. Hepatology 2023;78:1922–65.

[5] Furuta M, Ueno M, Fujimoto A, et al. Whole genome sequencing discriminates hepatocellular carcinoma with intrahepatic metastasis from multi-centric tumors. J Hepatol 2017;66:363–73.

[6] Torrecilla S, Sia D, Harrington AN, et al. Trunk mutational events present minimal intra- and inter-tumoral heterogeneity in hepatocellular carcinoma. J Hepatol 2017;67:1222–31.

[7] Miao R, Luo H, Zhou H, et al. Identification of prognostic biomarkers in hepatitis B virus-related hepatocellular carcinoma and stratification by integrative multi-omics analysis. J Hepatol 2014;61:840–9.

[8] Xu LX, He MH, Dai ZH, et al. Genomic and transcriptional heterogeneity of multifocal hepatocellular carcinoma. Ann Oncol 2019;30:990–7.

[9] Dong LQ, Peng LH, Ma LJ, et al. Heterogeneous immunogenomic features and distinct escape mechanisms in multifocal hepatocellular carcinoma. J Hepatol 2020;72:896–908.

[10] Boot GF, Haak F, Coto-Llerena M, et al. Multi-omics profiling reveals divergent biology and liver microenvironment in HCC of metastatic and de novo origin. Mol Cancer 2026;25.

[11] Lu Y, Xu L, Chen W, et al. Intrahepatic Microbial Heterogeneity in Multifocal Hepatocellular Carcinoma and Its Association with Host Genomic and Transcriptomic Alterations. Cancer Discov 2025;15:1630–48.

[12] Yang Y, Ni Q, Li H, et al. Genomic and the tumor microenvironment heterogeneity in multifocal hepatocellular carcinoma. Hepatology 2025;82:582–98.

[13] McGranahan N, Swanton C. Clonal Heterogeneity and Tumor Evolution: Past, Present, and the Future. Cell 2017;168:613–28.

[14] Hanahan D. Hallmarks of cancer-Then and now, and beyond. Cell 2026;189:2254–77.

[15] Seferbekova Z, Lomakin A, Yates LR, et al. Spatial biology of cancer evolution. Nat Rev Genet 2023;24:295–313.

[16] Xue R, Zhang Q, Cao Q, et al. Liver tumour immune microenvironment subtypes and neutrophil heterogeneity. Nature 2022;612:141–7.

[17] Jiang Z, Wu Y, Miao Y, et al. HCCDB v2.0: Decompose Expression Variations by Single-cell RNA-seq and Spatial Transcriptomics in HCC. Genomics Proteomics Bioinformatics 2024;22.

[18] Kang M, Armenteros JJA, Gulati GS, et al. Mapping single-cell developmental potential in health and disease with interpretable deep learning. bioRxiv 2024.

[19] Yin J, Wang S, Liu Y. TMED2 promotes thyroid cancer tumorigenesis by being involved in mTORC1-mediated fatty acid metabolism. Biochim Biophys Acta Gen Subj 2026;1870:130910.

[20] Xiao S, Jiang S, Wen C, et al. EMC2 promotes breast cancer progression and enhances sensitivity to PDK1/AKT inhibition by deubiquitinating ENO1. Int J Biol Sci 2025;21:2629–46.

[21] Wu X, Liu B, Liu Y, et al. Study on the role and mechanism of TMED2 in oral squamous cell carcinoma. Int J Biol Macromol 2025;289:138805.

[22] Dong X, Dai H, Yao L, et al. EMC2 promotes triple negative breast cancer growth by protecting FDFT1 from endoplasmic reticulum associated degradation to impair ferroptosis susceptibility. Oncogene 2025;44:3713–28.

[23] Wu X, Li T, Jiang R, et al. Targeting MHC-I molecules for cancer: function, mechanism, and therapeutic prospects. Mol Cancer 2023;22:194.

[24] Krenkel O, Tacke F. Liver macrophages in tissue homeostasis and disease. Nat Rev Immunol 2017;17:306–21.

[25] Prakash J, Shaked Y. The Interplay between Extracellular Matrix Remodeling and Cancer Therapeutics. Cancer Discov 2024;14:1375–88.

[26] Zhang Y, Song Y, Wang X, et al. An NFAT1-C3a-C3aR Positive Feedback Loop in Tumor-Associated Macrophages Promotes a Glioma Stem Cell Malignant Phenotype. Cancer Immunol Res 2024;12:363–76.

[27] Medler TR, Murugan D, Horton W, et al. Complement C5a Fosters Squamous Carcinogenesis and Limits T Cell Response to Chemotherapy. Cancer Cell 2018;34:561–78 e6.

[28] Wang Z, Sun C, Wang P, et al. Pan-cancer analysis of TMED2: unraveling potential immune characteristics and prognostic value in cancer therapy. Front Immunol 2025;16:1578627.

[29] Chen J, Zhang CH, Tao T, et al. A-to-I RNA co-editing predicts clinical outcomes and is associated with immune cells infiltration in hepatocellular carcinoma. Commun Biol 2024;7:838.

[30] Craig AJ, von Felden J, Garcia-Lezana T, et al. Tumour evolution in hepatocellular carcinoma. Nat Rev Gastroenterol Hepatol 2020;17:139–52.

[31] Artero MR, Minery A, Nedelcev L, et al. Complement and the hallmarks of cancer. Semin Immunol 2025;78:101950.

[32] Afshar-Kharghan V. The role of the complement system in cancer. J Clin Invest 2017;127:780–9.

[33] Zhang Y, Wang X, Gu Y, et al. Complement C3 of tumor-derived extracellular vesicles promotes metastasis of RCC via recruitment of immunosuppressive myeloid cells. Proc Natl Acad Sci U S A 2025;122:e2420005122.

[34] Niyonzima N, Rahman J, Kunz N, et al. Mitochondrial C5aR1 activity in macrophages controls IL-1beta production underlying sterile inflammation. Sci Immunol 2021;6:eabf2489.

[35] Donne R, Lujambio A. The liver cancer immune microenvironment: Therapeutic implications for hepatocellular carcinoma. Hepatology 2023;77:1773–96.

[36] Binnewies M, Roberts EW, Kersten K, et al. Understanding the tumor immune microenvironment (TIME) for effective therapy. Nat Med 2018;24:541–50.

[37] Dominguez M, Dejgaard K, Fullekrug J, et al. gp25L/emp24/p24 protein family members of the cis-Golgi network bind both COP I and II coatomer. J Cell Biol 1998;140:751–65.

