## Supplementary methods and figures for "Divergent malignant evolution converges on shared immunosuppressive ecosystems through tumor-intrinsic TMED2–ADAR1-dependent complement signalling in multifocal hepatocellular carcinoma"

### 1. Supplementary methods

#### Patient cohort and study design

All patients with primary multifocal HCC were enrolled in this study. All patients were newly diagnosed, harbored at least two anatomically distinct tumor foci, and had received no anticancer treatment before surgical resection. Hepatitis virus infection was not used as an inclusion or exclusion criterion. Each tumor focus and its matched adjacent non-tumor liver tissue were collected separately during surgery. Detailed clinicopathological characteristics of the patients are provided in **online supplemental Table 1**.

Integrated single-cell and spatial multi-omics profiling was performed on samples from five patients, including scFAST-seq (15 tumor/adjacent tissue samples), scATAC-seq (12 samples), and Stereo-seq (14 samples). For independent validation, a tissue microarray comprising 48 paired multifocal HCC tumor foci and matched adjacent liver tissues (**online supplemental Table 2**) was constructed by Bioaitech Co., Ltd. (China), and bulk RNA sequencing was performed on 13 samples from 10 patients (**online supplemental Table 1**).

Publicly available datasets were also analyzed, including scRNA-seq data from the HRA001748 cohort[1] (n = 88) and bulk RNA-seq data from TCGA-LIHC (n = 466, UCSC Xena, https://xena.ucsc.edu/) and OEP000321[2] (n = 316). All patients provided written informed consent, and the study was approved by the Ethics Committee of Guangzhou Medical University (approval no. 202405020).

#### scFAST-seq library construction and sequencing

Fresh tumor and adjacent normal tissues were dissociated using the Multi Tissue Dissociation Kit 2 (Miltenyi 130-110-203). After erythrocytes removal (Miltenyi 130-094-183), cell number and viability were estimated using

Fluorescence Cell Analyzer (Countstar® Rigel S2) with AO/PI reagent, followed by debris and dead cell depletion (Miltenyi 130-109-398/130-090-101). Fresh cells were washed in RPMI1640 and resuspended in 1× PBS with 0.04% BSA at $1\times10^6$ cells/mL.

Single-cell RNA-seq libraries were prepared using the SeekOne® Single Cell Whole Transcriptome Kit. Cells, reverse transcription reagents, Barcoded Hydrogel Beads (BHBs), and partitioning oil were loaded onto the SeekOne® DD Chip S3 and encapsulated into emulsion droplets using the SeekOne® Digital Droplet System. After barcode incorporation, cDNA was purified, ribosomal/mitochondrial transcripts were removed, and libraries were constructed and indexed. Sequencing was performed on Illumina NovaSeq 6000 with 150 paired-end reads.

Raw reads were trimmed using fastp (v 0.20.1) to remove adaptors and low-quality bases. Clean reads were processed with SeekSoul Tools (v1.2.0) to generate expression matrices. Cell barcodes and UMIs were extracted, corrected, and aligned to the reference genome (GRCh38) using STAR. Gene counting was done using featureCounts (v1.6.4), with chemistry-specific strand settings and exon/intron options. A Cell Ranger–like algorithm, similar to EmptyDrops, was used to identify valid cells and generate a filtered UMI count matrix. Detailed information on sequenced samples and the number of captured cells is provided in **online supplemental Table 3**.

**scATAC-seq library construction and sequencing**

Nuclei isolated from tumor and adjacent normal tissues were resuspended in Nuclei Buffer (10x Genomics, Chromium Single Cell ATAC Reagent Kit). Nuclei suspensions were incubated with a transposition mix containing Tn5 transposase, which preferentially fragments accessible chromatin and simultaneously tags DNA fragments with adapter sequences. Transposed nuclei were loaded onto a Chromium Chip E with barcoded Gel Beads,

Master Mix, and Partitioning Oil to generate Gel Bead-in-Emulsions (GEMs). Following GEM generation, silane magnetic beads were used to remove leftover biochemical reagents, and SPRI beads were applied to eliminate unused barcodes. P7, a sample index, and Read 2 sequence are added during library construction via PCR. Sequencing was performed on the Illumina NovaSeq 6000 platform. Raw reads were aligned to the reference genome (GRCh38) using the Cell Ranger ATAC pipeline (https://support.10xgenomics.com/single-cell-atac/software/overview/welcome), and peak-barcode count matrices were generated.

**Stereo-seq chip preparation and sequencing**

Tumor and adjacent normal tissues were embedded in OCT (Sakura, 4583), snap-frozen in liquid nitrogen-prechilled isopentane, and stored at -80°C. Cryosections (10 µm) were mounted onto Stereo-seq chips (BGI, China), incubated at 37°C for 3 minutes, fixed with methanol at −20°C for 40 minutes, and optionally stained with nucleic acid dye (Thermo Fisher, Q10212) for imaging. Sections were washed and permeabilized with 0.1% pepsin (Sigma, P7000) in 0.01 M HCl at 37°C for 5 minutes. RNA released from permeabilized tissues was captured by DNA nanoballs (DNBs) and reverse transcribed overnight at 42°C using SuperScript II (Invitrogen). After tissue digestion and cDNA release, cDNA was purified and amplified with KAPA HiFi Hotstart Ready Mix (Roche, KK2602). Amplified DNA (20 ng) was fragmented with in-house Tn5 transposase, followed by a second round of PCR and purification. Libraries were used for DNB generation and sequenced on the MGI DNBSEQ-Tx platform (performed by Beijing Novogene Technology Co., Ltd.).

Fastq files were processed using the SAW pipeline (https://github.com/BGIResearch/SAW). Cell barcodes (CID) and molecular barcodes (MID) were extracted from Read 1 (CID: bases 1–25, MID: bases 26–35), while Read 2 contained cDNA sequences. Reads with low-quality MIDs or invalid barcodes were filtered out. Clean reads were aligned to the reference genome (GRCh38) using STAR, and uniquely mapped reads (MAPQ >10) were assigned to genes. UMIs with the same CID and gene locus were collapsed to generate a spatial gene expression matrix.

**Bulk RNA extraction and sequencing**

Total RNA was isolated from HCC tumor and matched adjacent liver tissues. Ribosomal RNA was depleted to enrich for coding and non-coding transcripts. The remaining RNA was fragmented and used to generate strand-specific cDNA via first- and second-strand synthesis, incorporating dUTP to retain strand information. cDNA libraries were then end-repaired, A-tailed, adapter-ligated, PCR-amplified, and size-selected. Libraries were purified with magnetic beads (VAHTS DNA Clean Beads or AMPure XP beads), quantified, and sequenced on Illumina platforms according to the manufacturers' protocols.

**scFAST-seq data processing**

The UMI count matrix of each sample was processed using the Seurat (v5)[3] package in R. For each individual sample, low-quality cells were filtered out based on the following criteria: fewer than 200 detected genes, more than 6,000 genes, over 40% mitochondrial counts, or more than 50% ribosomal counts. Additionally, genes retained for downstream analysis were required to be expressed in at least three cells. After quality control, each sample was independently processed, including normalization (NormalizeData()), identification of highly variable genes (FindVariableFeatures()), scaling (ScaleData()), and principal component analysis (RunPCA()).To integrate multiple samples and correct for batch effects, we applied the Reciprocal PCA

(RPCA) integration method. The integrated dataset was then re-scaled (ScaleData()) and re-run through PCA (RunPCA()) for dimensionality reduction. Cell clustering was performed using FindNeighbors() and FindClusters(), with clustering resolutions set at multiple levels (e.g., 0.1, 0.6, and 1.2) to identify distinct cell populations. Finally, RunUMAP() was used to visualize the clusters in two-dimensional space.

**scATAC-seq data processing**

We analyzed the peak-barcode matrix using the Signac package[3]. A unified peak set was first generated using combined.peaks(). Fragment files for each sample were processed with CreateFragmentObject(), and peak-by-cell count matrices were constructed using FeatureMatrix(). These matrices were used to create individual Seurat objects via CreateChromatinAssay(). All Seurat objects were merged using merge(), ensuring consistency across datasets by using a shared peak set. The merged object retained the fragment references and cell name mappings, allowing efficient downstream analysis. We then applied RunTFIDF(), FindTopFeatures(), RunSVD(), and RunUMAP() for normalization, feature selection, dimensionality reduction, and visualization, respectively. Chromatin accessibility at specific genomic regions was assessed using CoveragePlot() to further validate cell identities and regulatory activity.

**Stereo-seq data processing**

We analyzed the spatial transcriptomics data using Stereopy (https://github.com/STOmics/Stereopy), a comprehensive toolkit tailored for the processing and visualization of Stereo-seq data. To address the low RNA capture efficiency at the single DNB resolution (~500 nm), the raw spatial expression matrix was aggregated into larger pseudo-spots using a convolution window of 50 × 50 DNBs (referred to as bin50), which corresponds to a spatial resolution of approximately 25 μm per spot. Therefore, we used the read_gef() to read the data with bin_size=50, and integrated multiple tissue sections using MSData(). The integrated data then underwent standard preprocessing steps, including total count normalization with normalize_total() and log-transformation via log1p(). We performed dimensionality reduction using pca(), followed by batch effect correction across sections with batches_integrate(). To identify spatial clusters, we constructed a neighborhood graph using neighbors() and applied the Leiden algorithm (leiden()) for clustering. Finally, we used umap() for visualization of the spatial domains, allowing us to explore transcriptional heterogeneity and spatial organization within the multifocal HCC samples.

**Cell type annotation**

For the scFAST-seq dataset, cell type annotation was performed by identifying marker genes using Seurat's FindAllMarkers(), which compares each cluster against all others using the Wilcoxon rank-sum test. Marker genes were retained if they were expressed in at least 25% of cells within a cluster and had a minimum log fold change of 0.25. Based on a manually curated list of known marker genes for major HCC-associated cell types, we annotated the primary cell populations, including hepatocytes, T cells, NK cells, B cells, fibroblasts, and endothelial cells. To further resolve immune and stromal heterogeneity, we conducted secondary clustering on relevant subsets and used well-established subtype-specific markers to define T cell subtypes, myeloid cell subtypes, B cell subtypes, as well as fibroblast and endothelial cell subtypes.

For the scATAC-seq dataset, cell type annotations were inferred by transferring labels from the annotated scFAST-seq dataset using Signac's FindTransferAnchors() and TransferData(). To ensure the reliability of the transferred identities, we applied an additional filtering step, retaining only those scATAC cells whose predicted labels were consistent with their de novo clustering results. Subtype annotations for immune and stromal populations in the scATAC dataset were similarly assigned via label transfer from the scFAST-derived subtypes, ensuring coherence and comparability across modalities.

For the Stereo-seq dataset, cell type annotation was performed using find_marker_genes() to identify marker genes for each spatial cluster. Based on the expression of known marker genes and the spatial distribution of clusters across tissue sections, we assigned putative cell identities. Given that the bin50 resolution may not correspond to single-cell precision, we categorized clusters into broader tissue-level cell types, including malignant hepatocytes, peri-tumoral normal hepatocytes, and stromal cell types.

**Identification of A-to-I RNA editing sites based on scFAST-seq data**

To identify A-to-I RNA editing events at single-cell resolution, we established a five-step analysis pipeline based on scFAST-seq data. First, raw paired-end FASTQ files were demultiplexed using a custom script that separated reads into cell-specific FASTQ files based on a predefined list of 17-bp barcodes. Second, the demultiplexed reads were individually aligned to the human reference genome (GRCh38) using STAR in two-pass mode to improve alignment accuracy and splice junction detection. Third, cell-specific BAM files were processed using REDItools 2.0[4] to identify RNA-DNA mismatches, with a focus on A-to-G substitutions indicative of A-to-I editing. Fourth, candidate editing sites within each cell were filtered using the following criteria: minimum read coverage of ≥2 and editing frequency ≥5%. Fifth, filtered sites were aggregated across cells within each sample, and only sites present in at least 10 individual cells were retained. This process yielded a high-confidence A-to-I editing matrix comprising 85,830 unique sites across 167,658 single cells, including 80,399 sites located on autosomes (chromosomes 1-22).

In parallel, we profiled A-to-I RNA editing using bulk RNA-seq data. For each sample, raw FASTQ files were aligned to the GRCh38 reference genome using STAR in two-pass mode, followed by mismatch detection using REDItools 2.0. Candidate editing sites were filtered using a minimum read coverage of 10 and an editing frequency threshold of ≥10%. Across all samples, we identified 23,925 A-to-I sites, 10,168 of which overlapped with those detected from the scFAST-seq data, supporting the reliability and reproducibility of our single-cell RNA editing pipeline.

To further exclude potential genomic variants, we leveraged matched DNA-seq data from five samples. BAM files were processed using REDItools 2.0 to identify DNA-level mismatches, with filtering criteria including a minimum read coverage of ≥10, editing frequency ≥10%, and bidirectional strand support. A total of 1,338,337 A-to-I substitutions were identified. After excluding DNA-originated sites, 42,745 A-to-I sites remained from the scFAST-seq dataset. To eliminate potential germline variants, we annotated the candidate A-to-G editing sites using ANNOVAR[5] with reference to the gnomAD (gnomad41_genome) and dbSNP (avsnp151) databases. Sites that were annotated in either database were removed. After this stringent filtering, 29,897 high-confidence A-to-I editing sites remained, which were subsequently used for downstream comparative and functional analyses.

**Identification of malignant cells based on InferCNV**

Copy number variation (CNV) analysis was performed using inferCNV (v1.18.1; https://github.com/broadinstitute/infercnv) on hepatocytes, with adjacent non-tumor hepatocytes as reference. The raw count matrix was analyzed using infercnv::run() with a cutoff of 0.1. Cells were clustered by CNV profiles using kmeans() to distinguish malignant from non-malignant hepatocytes. The same approach was applied to public scRNA-seq datasets for consistent identification of malignant cells.

##### Inference of CNVs from scATAC-seq

CNV profiles from scATAC-seq were inferred using epiAneufinder[6] (https://github.com/colomemaria/epiAneufinder). The epiAneufinder() function was applied to the peak matrix with hg38. Cells with <20,000 fragments were excluded (minFrags = 20000), and sex chromosomes and mitochondrial DNA were removed. CNV profiles were visualized using plot_single_cell_profile().

##### Hepatocyte differentiation potential using CytoTRACE2

CytoTRACE2[7] (https://github.com/digitalcytometry/cytotrace2) was used to estimate hepatocyte differentiation potential, assigning a potency score from 0 (differentiated) to 1 (totipotent). Analysis was performed on the raw counts matrix with batch_size = 10000 and smooth_batch_size = 1000, grouped by sample, and visualized using plotData().

##### Gene set enrichment analysis

Differentially expressed gene sets (DEGs) were analyzed for GO and KEGG pathway enrichment via clusterProfiler. T cell-related gene sets (cytotoxicity, exhaustion, proliferation; **online supplemental Table 4**) were assessed by first calculating differential expression per T cell subtype with wilcoxauc(), ranking gene sets by AUC, and computing enrichment scores using fgsea(). Gene sets with adjusted $P < 0.05$ were considered significant. Positive normalized enrichment score indicates preferential enrichment at the top of the ranked list.

##### Correlation analysis

For hepatocytes, subclonal distributions across samples were calculated in scFAST-seq and scATAC-seq datasets, and correlations between the two were assessed. To identify corresponding myeloid, fibroblast, and endothelial subtypes in public scRNA-seq datasets, average gene expression per subtype was computed using AverageExpression(), and Spearman correlations were calculated with corr.test().

#### Tumor clonal evolution

We applied inferCNV to identify CNV events and reconstructed phylogenetic trees using the uphyloplot2.py script (https://github.com/harbourlab/uphyloplot2)[8]. CNV amplifications and deletions were annotated with data from pred_cnv_regions.dat, and their frequencies were calculated. CNVs on the upper trunk indicate early clonal events, while those on lower branches reflect later subclonal evolution.

#### Deconvolution analysis

Immune cell composition was estimated using the CIBERSORT[9] with the LM22 signature. Gene expression matrices from TCGA-LIHC, OEP000321, and in-house multifocal HCC datasets were analyzed with 1,000 permutations and quantile normalization disabled (QN = FALSE). The estimated abundances across the three cohorts are provided in **online supplemental Table 5**.

#### Pseudo-time analysis

Hepatocyte differentiation trajectories across tumor and adjacent tissues were inferred using Monocle2[10] (https://github.com/cole-trapnell-lab/monocle2-rge-paper). Highly variable genes were selected, dimensionality reduced via reduceDimension(max_components = 2, method = "DDRTree"), and cells ordered with orderCells(). For macrophages, fibroblasts, and endothelial subtypes, Monocle3 was used. Data were preprocessed (preprocess_cds()), batch-corrected (align_cds()), and trajectories constructed with reduce_dimension(), cluster_cells(), and learn_graph().

**Cell-cell interactions**

Cell-cell communication was analyzed using CellChat (v2.1.2)[11], integrating gene expression with known ligand–receptor interactions. CellChat objects were created with createCellChat() and CellChatDB.human database. Tumor and adjacent samples were analyzed separately, then integrated with mergeCellChat() to compare signaling patterns. Interaction differences were visualized with netVisual_bubble() and netVisual_chord_gene().

**Survival analysis**

Patients were divided into high- and low-expression groups for each target gene, and Kaplan-Meier survival curves were generated using the survival (v3.7.0) and survminer (v0.4.9). Tumor size was analyzed similarly. For cell-type-specific signatures, top 20 marker genes per cluster were used to calculate GSVA (v1.50.0) enrichment scores, and patients were stratified into high- and low-score groups for survival analysis.

**Multiplex immunofluorescence staining and analysis**

Multiplex immunofluorescence (mIHC) was performed using a tyramide signal amplification (TSA)-based five-color kit (Recordbio, Shanghai, China) following the manufacturer's instructions. Tissue microarray sections and OCT-embedded frozen tumor and adjacent tissue sections were processed for multiplex immunohistochemistry, including fixation, antigen retrieval, blocking, and overnight incubation at 4 °C with primary antibodies (CD3, HUABIO, Cat. No. HA720082, 1:500; CD20, HUABIO, Cat. No. HA721138, 1:1500; α-SMA, Proteintech, Cat. No. 14395-1-AP, 1:4000; CD56, HUABIO, Cat. No. HA722755, 1:500; C3, Proteintech, Cat. No. 21337-1-AP, 1:100; C3AR1, Abcam, Cat. No. Ab317321, 1:1600; CD163, Abclonal, Cat. No. A25206, 1:1600), followed by secondary antibodies (ZSGB-BIO, Cat. No. ZB-

2301, 1:200) and fluorophores. Nuclei were counterstained with DAPI, and images acquired by fluorescence microscopy.

Acquired images were analyzed using the PhenoVision mIF AI analysis system (PhenoVision Bio Co., Ltd.), based on the Oncotopix Discovery platform (Visiopharm, version 4.5.6.5, Hoersholm, Denmark). An AI-based analysis workflow was trained to identify individual cells using DAPI staining and classify cellular phenotypes according to protein fluorescence signals. Quantitative analysis was performed for different cell populations, including the number and proportion of CD163+, CD163+C3+C3AR+, and $CD163^{-}$ C3+C3AR+ cells. The mean fluorescence intensity of C3, C3AR, and CD163 was further quantified in CD163+C3+C3AR+ and $CD163^{-}$ C3+C3AR+ cells.

**Overexpression and knockdown**

To generate stable *TMED2* or *EMC2* overexpressing cell lines, recombinant lentiviral vectors (pLV3-CMV--3×FLAG-mCherry-Puro, Miaoling Plasmid Platform, China, China) were packaged in HEK293T cells and used to transduce HepG2 cells, followed by puromycin selection. For gene knockdown, TMED2-, EMC2-, ADAR1- specific siRNAs (GENEWIZ, China) were transiently transfected into HepG2 or Huh7 cells using Lipofectamine® RNAiMAX (Thermo Fisher Scientific, USA), and knockdown efficiency was confirmed. The specific siRNA sequences are listed in **online supplemental Table 6**.

**Cell viability and migration assay**

Cells with *TMED2* or *EMC2* overexpression or knockdown were seeded in 96-well plates (8,000 cells/well, 5 replicates) for proliferation assays. At indicated time points, 20 μL CCK-8 reagent (Beyotime, China) was added, incubated 1.5 h, and absorbance measured at 450 nm. For migration assays, $1\times10^{5}$

cells were seeded in 24-well transwell inserts (CLS3415-48EA, Corning, USA) with 10% FBS medium in the lower chamber. After 24 h, migrated cells were fixed with 4% glutaraldehyde, stained with crystal violet (Beyotime, China), and counted in four random fields.

##### ELISA assay

C3A and C5A in cell culture supernatants were measured using Human C3A (Cat. No. EH2734) and C5A (Cat. No. EH0096), and ELISA kits (Finetest, China) following the manufacturer's instructions.

##### Quantitative PCR

Total RNA was extracted from cells using the Super FastPure Cell RNA Isolation Kit (Vazyme, China). Complementary DNA (cDNA) was synthesized using HiScript IV All-in-One Ultra RT SuperMix (Vazyme, China). qPCR was performed on a CFX96 system (Bio-Rad, USA) using SupRealQ Purple Universal SYBR qPCR Mix (U+) (Vazyme, China). All reactions were performed in technical triplicates. Gene-specific primers are listed in **online supplemental Table 7**. Relative expression levels were calculated using the $2^{(-\Delta\Delta Ct)}$ method, with *GAPDH* as the internal control.

#### 2. Supplementary figures S1 - S24

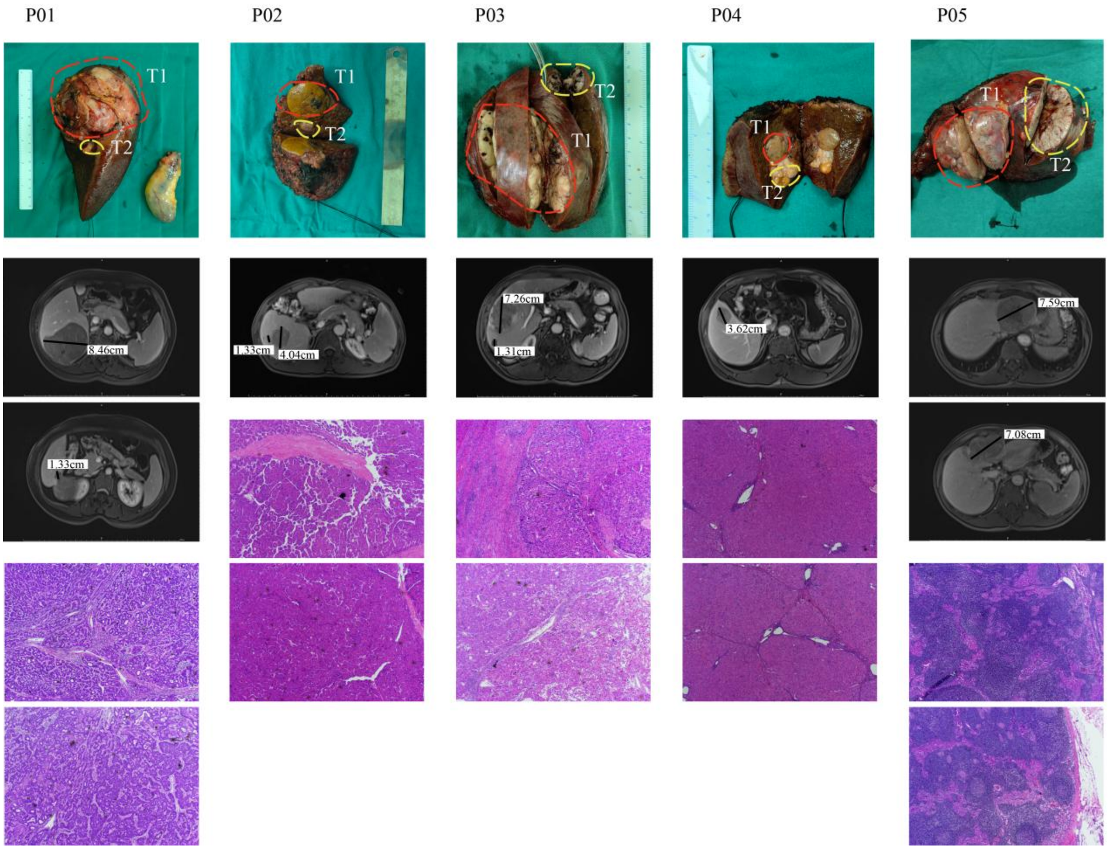

**Figure 1. Clinicopathologic characteristics of patients with multifocal hepatocellular carcinoma (HCC).**

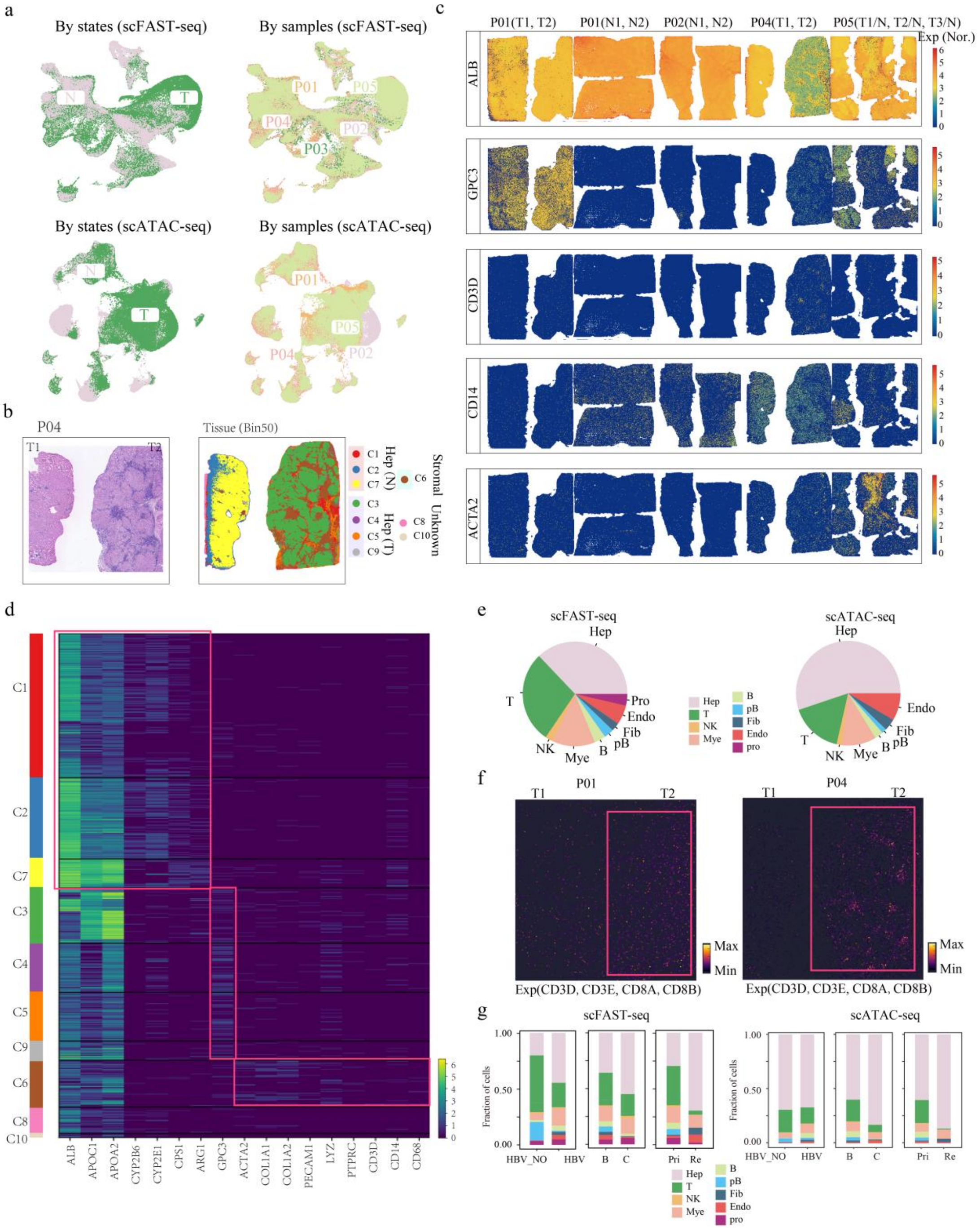

**Figure 2. Integrated single-cell and spatial multi-omics analysis reveals cellular heterogeneity, and spatial organization, and clinical subgroup-associated compositional differences across tumor foci. (a)** UMAP plots of cell lineages identified from scFAST-seq (top) and scATAC-seq (bottom), colored by cell state and patient. **(b)** H&E staining and spatial distribution of identified cell types across different tumor foci in patient P04, based on Stereo-seq. **(c)** Spatial expression patterns of selected marker genes on

tissue sections. **(d)** Heatmap displaying the expression of cell cluster–associated genes based on Stereo-seq data. **(e)** Pie charts showing the proportions of major cell types based on scFAST-seq (top) and scATAC-seq (bottom). **(f)** Expression patterns of T cell marker genes across different tumor foci in patients P01 and P04. **(g)** Stacked bar charts comparing cell type proportions across different clinical subgroups, including HBV vs. HBV_No, Type B vs. Type C, and Primary (Pri) vs. Recurrent tumors (Re), in both scRNA-seq (top) and scATAC-seq (bottom).

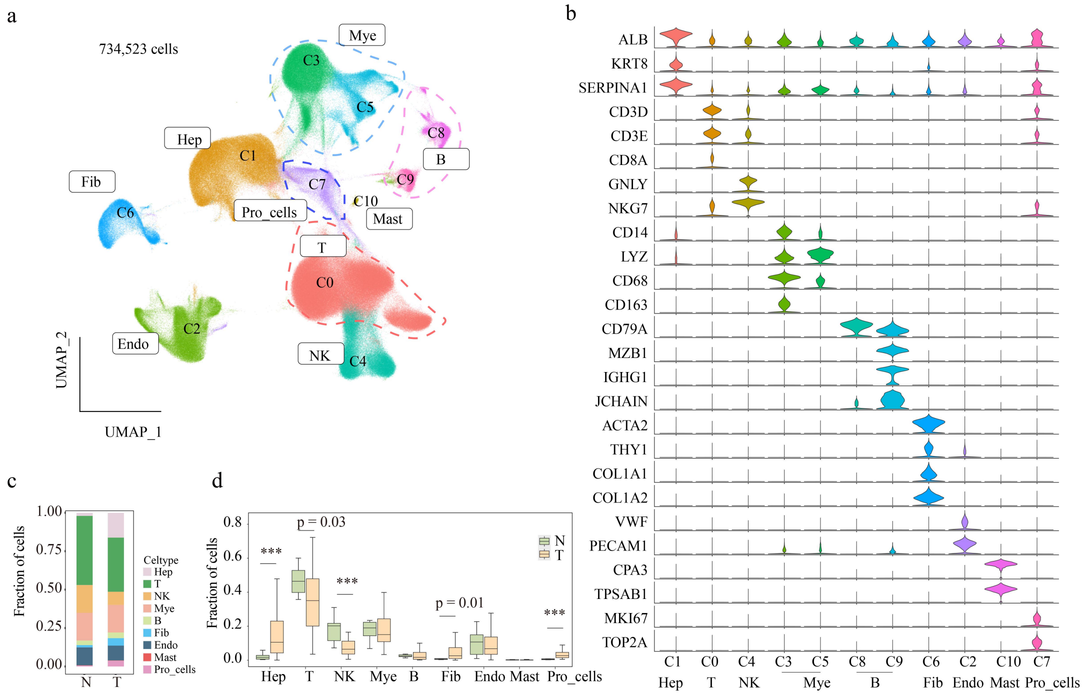

**Figure 3. Cell type identification, annotation, and compositional differences between adjacent non-tumor and tumor tissues based on public scRNA-seq analysis. (a)** UMAP plots of cell lineages identified from the public scRNA-seq dataset (HRA001748), colored by cell clusters. **(b)** Violin plots illustrating the expression levels of representative marker genes across different cell types. **(c)** Stacked bar charts showing the proportions of cell types in adjacent non-tumor (N) and tumor (T) tissues. **(d)** Boxplots comparing the abundance of specific cell types between adjacent non-tumor (N) and tumor (T) tissues.

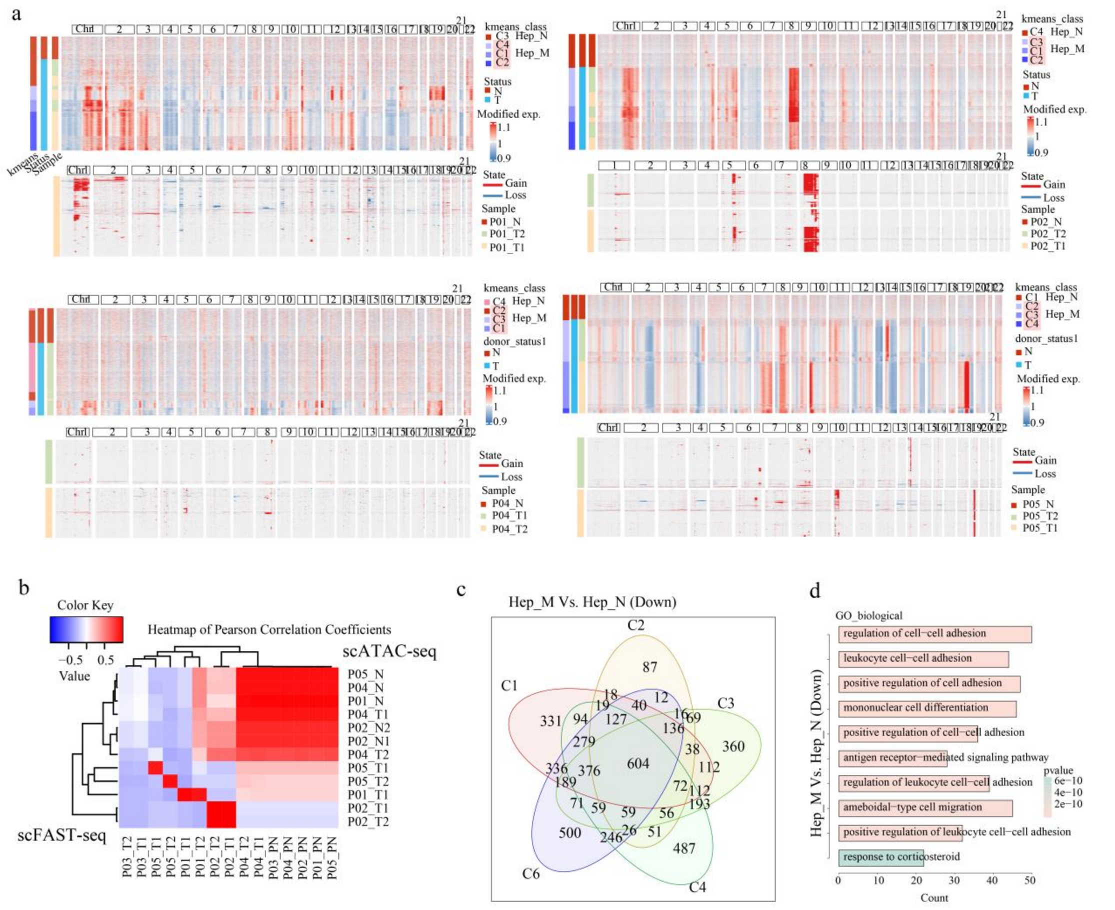

**Figure 4. Copy number variation landscapes, inter-assay concordance, and functional characterization of shared downregulated genes in hepatocyte subpopulations. (a)** Heatmap showing large-scale CNVs across individual hepatocytes (rows) from a single patient, inferred by scFAST-seq (top) and scATAC-seq (bottom). Red indicating amplifications and blue indicating deletions. **(b)** Correlation of CNV profiles between samples based on scFAST and scATAC data. **(c)** Venn diagram illustrating the overlap of downregulated genes among Hep_M clusters. **(d)** Functional enrichment analysis of the overlapping genes.

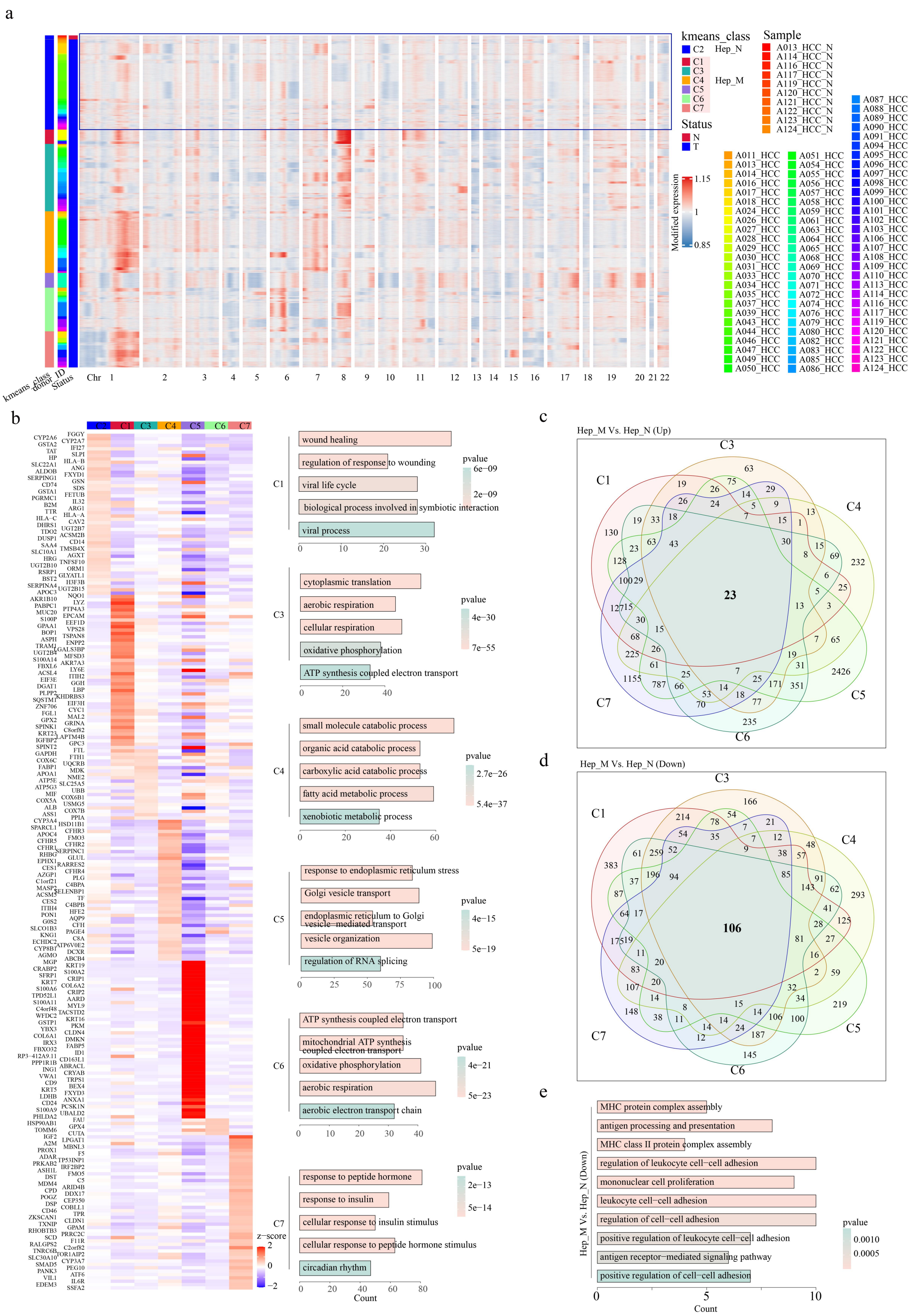

**Figure 5. Copy number variation inference, transcriptomic heterogeneity, and functional enrichment of differentially expressed genes in hepatocyte subpopulations based on public scRNA-seq analysis. (a)** Heatmap illustrating large-scale CNVs across individual hepatocytes (rows),

inferred from the public scRNA-seq data (HRA001748). **(b)** Heatmaps displaying differentially expressed genes across hepatocyte clusters, alongside corresponding functional enrichment analyses. **(c,d)** Venn diagrams showing the overlap of upregulated **(c)** and downregulated **(d)** genes identified by scRNA-seq. **(e)** Functional enrichment analysis of the overlapping downregulated genes.

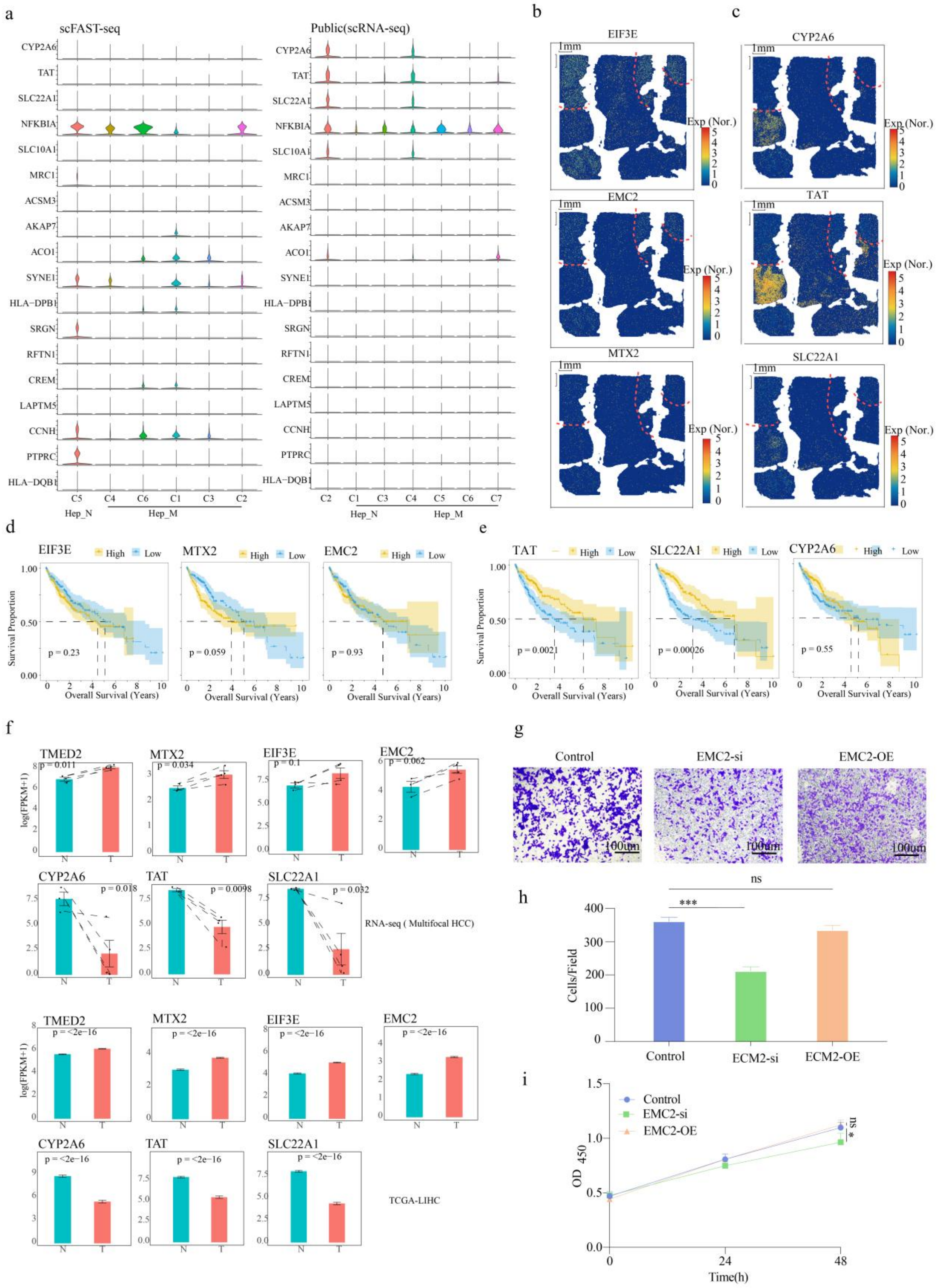

**Figure 6. Identification, spatial validation, clinical relevance, and functional characterization of key genes associated with malignant hepatocyte subtypes. (a)** Violin plots showing the expression of genes commonly downregulated across all malignant cell subtypes as identified by scFAST-seq (left) and public scRNA-seq (right). **(b,c)** Spatial expression

patterns of *EIF3E*, *EMC2*, and *MTX2* **(b)**, and *CYP2A6*, *TAT* and *SLC22A1* **(c)** in tissue sections from patient P05, profiled using Stereo-seq. **(d,e)** Kaplan-Meier survival analyses for *EIF3E*, *EMC2*, and *MTX2* **(d)**, *CYP2A6*, *TAT* and *SLC22A1* **(e)**. **(f)** Bar plots showing the expression changes of *TMED2*, *MTX2*, and other genes in our mutifocal HCC RNA-seq dataset and the TCGA-LIHC dataset. **(g)** Representative cell invasion results of HepG2 cells following *EMC2* knockdown (siRNA) or overexpression. **(h)** Quantification of *EMC2* knockdown and overexpression efficiency in HepG2 cells. **(i)** Functional assays assessing the effects of *EMC2* knockdown and overexpression in HepG2 cells.

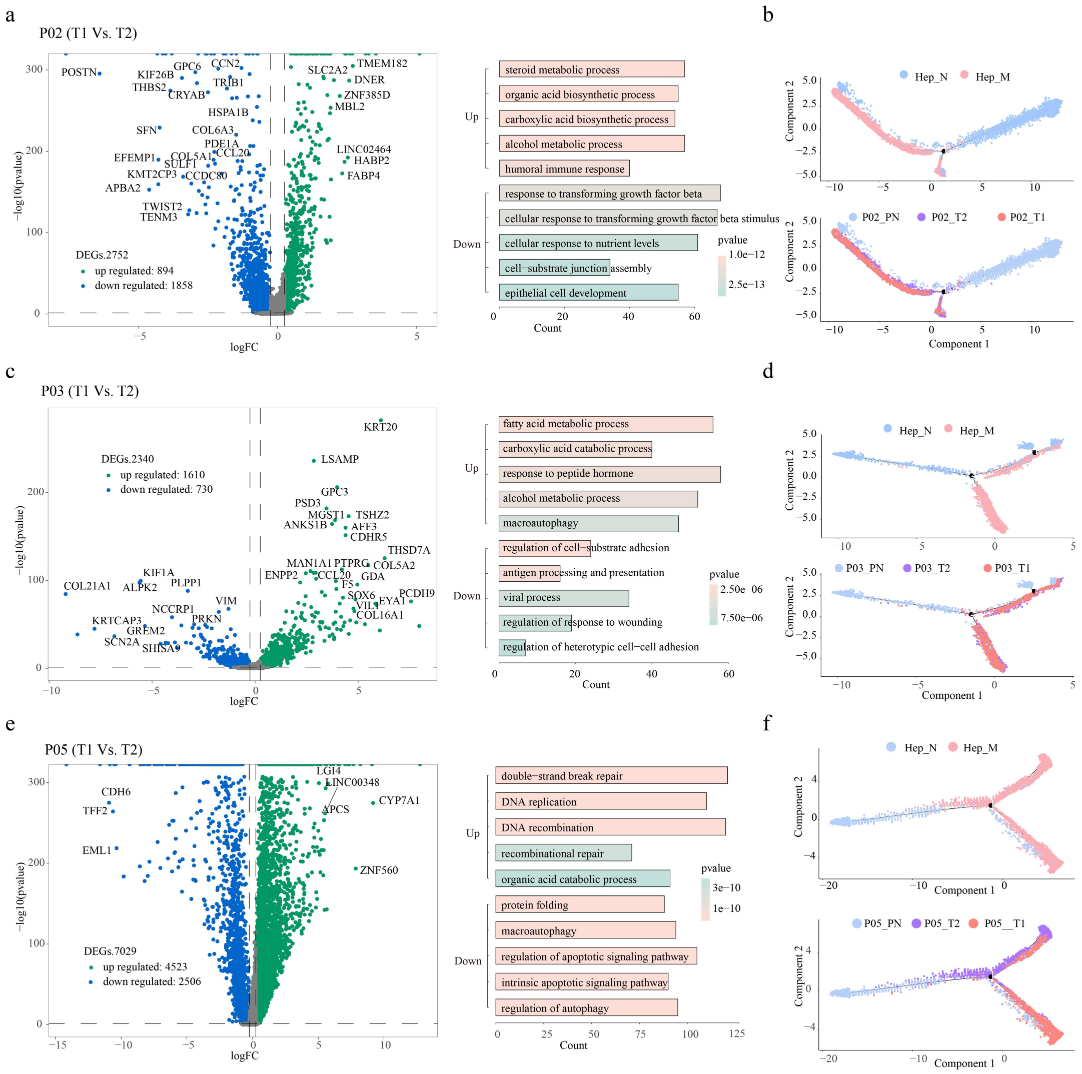

**Figure 7. Differential gene expression profiling, functional enrichment, and pseudotime trajectory analysis of malignant hepatocytes across multiple tumor foci. (a,c,e)** Volcano plots (left) illustrating differentially expressed genes between malignant cells from focus T1 and T2 in patient P02 **(a),** P03 **(c)**, and P05 **(e)**, alongside bar charts (right) summarizing the functional enrichment of upregulated and downregulated genes. **(b,d,f)** Developmental trajectories of hepatocytes in P02 **(b)**, P03 **(d)**, and P05 **(f)** inferred by Monocle2, with cells colored by cell state (top) and sample origin (bottom).

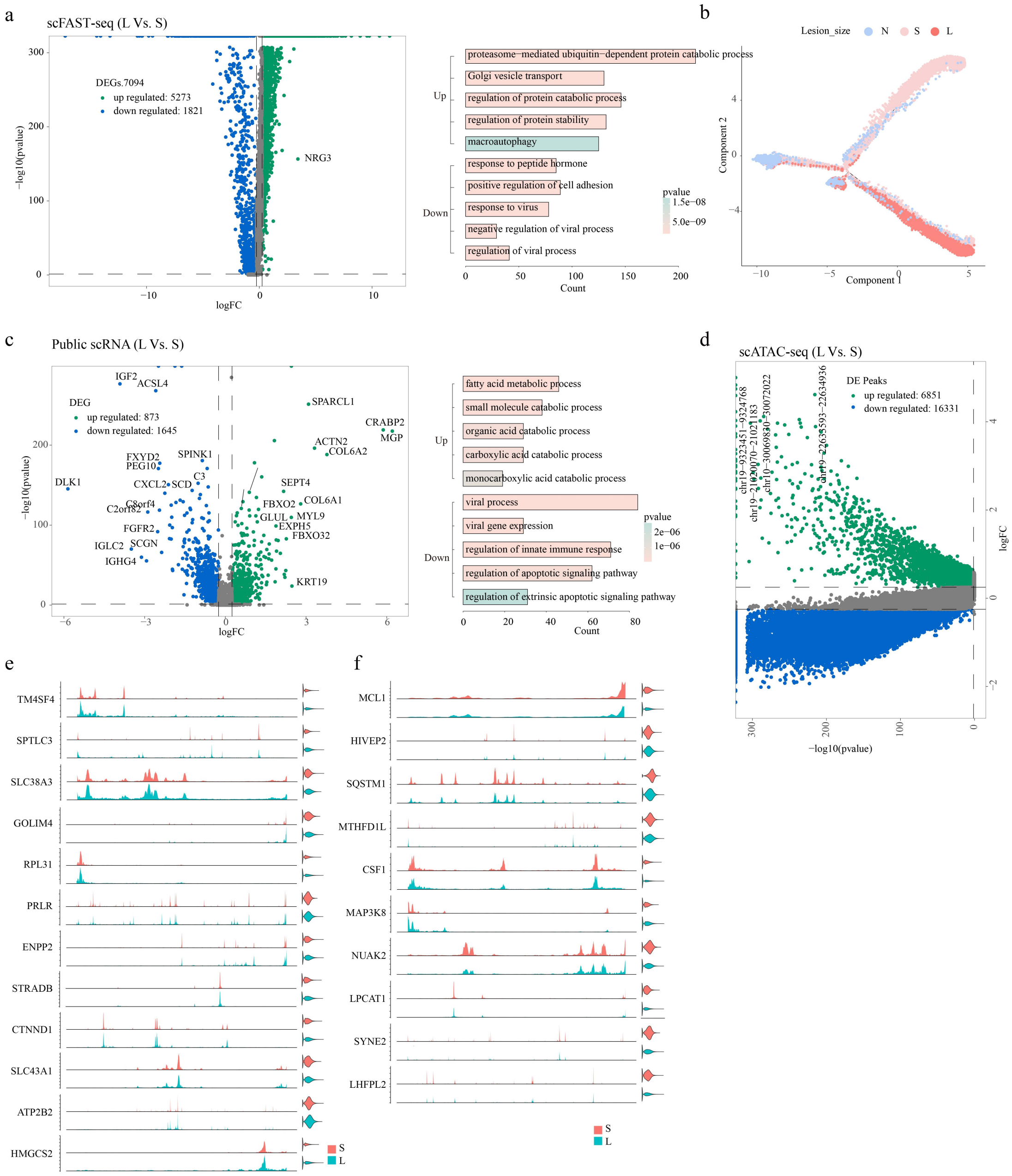

**Figure 8. Multi-omics comparison of malignant hepatocytes between large and small tumor foci reveals transcriptomic, epigenomic, and trajectory-level differences. (a,c)** Volcano plots (left) illustrating differentially expressed genes between malignant cells from large foci and small foci in the scFAST **(a)** and public scRNA-seq **(c)** datasets, alongside bar charts (right) summarizing the functional enrichment of upregulated and downregulated genes. **(b)** Developmental trajectories of hepatocytes from adjacent non-tumor tissue, small foci, and large foci, inferred by Monocle2. **(d)** Volcano plot illustrating differentially accessible chromatin peaks between malignant cells

from large foci and small foci in the scATAC-seq dataset. **(e,f)** Chromatin accessibility peaks of 13 upregulated **(e)** and 10 downregulated **(f)** genes in small and large tumor foci based on scATAC-seq data.

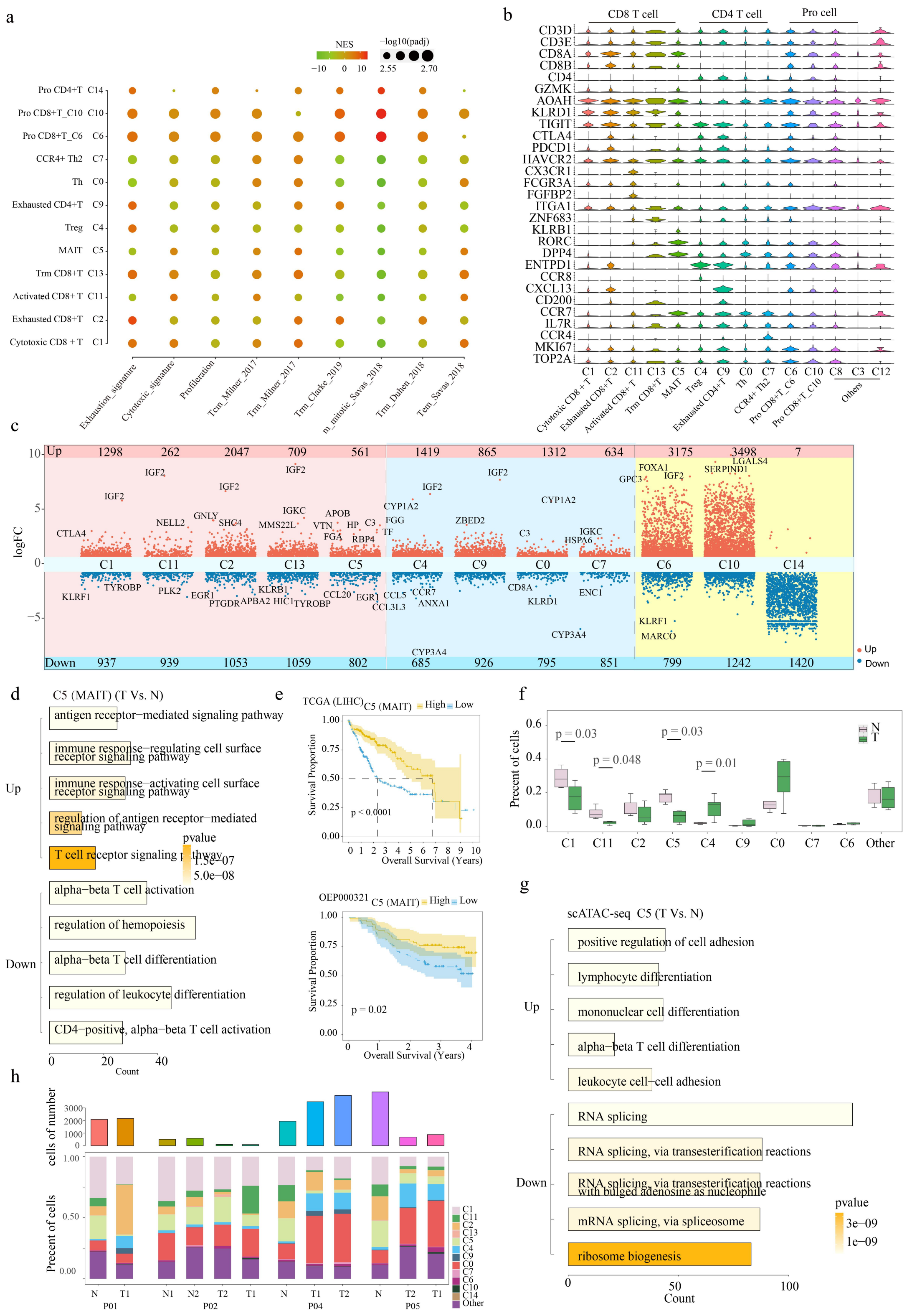

**Figure 9. T cell subtype landscape, differential activity, functional enrichment, and clinical relevance in tumor and adjacent non-tumor tissues. (a)** Bubble plot showing the enrichment of different T cell subtypes. **(b)** Violin plots displaying the expression of subtype-specific marker genes

across T cell subtypes, based on scATAC-seq data. **(c)** Scatter chart showing DEGs (dots) in T cell subtypes between adjacent non-tumor (N) and tumor (T) tissues. Red and blue dots indicate upregulated and downregulated genes, respectively; bar charts above and below represent the number of DEGs. **(d)** Functional enrichment analysis of upregulated and downregulated genes in MAIT (C5) within tumor tissues. **(e)** Kaplan-Meier survival analysis of MAIT (C5) cells in the TCGA-LIHC and OEP000321 datasets. **(f)** Boxplots showing the variation in T cell subtype proportions between adjacent non-tumor (N) and tumor (T) samples, based on scATAC-seq data. **(g)** Functional enrichment analysis of upregulated and downregulated genes in MAIT (C5) cell subtypes within tumor tissues, based on scATAC-seq data. (**h**) Stacked bar charts displaying the T cell subtype composition across different foci, based on scATAC-seq data.

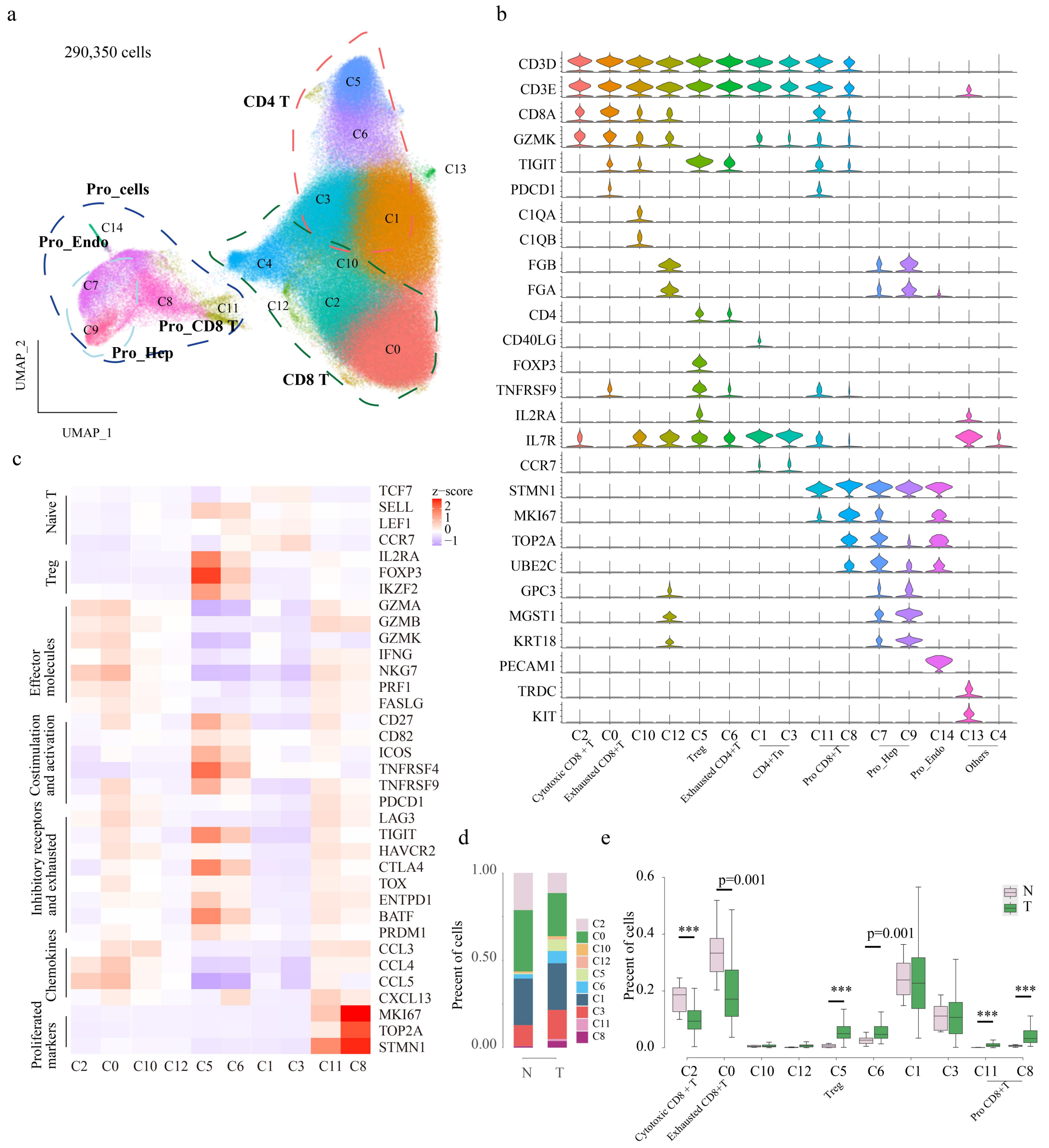

**Figure 10. Validation of T cell subtype landscape and compositional changes between adjacent non-tumor and tumor tissues using public scRNA-seq datasets. (a)** UMAP plots showing T cell lineages identified by the public scRNA-seq (HRA001748) analysis. **(b)** Violin plots displaying the expression of subtype-specific marker genes across T cell subtypes, based on the public scRNA-seq data. **(c)** Heatmap of scaled normalized gene expression within T cell clusters, based on the public scRNA-seq data. **(d)** Stacked bar charts showing the proportion of T cell subtypes in adjacent non-tumor and tumor samples, based on the public scRNA-seq data. **(e)** Boxplots

showing the variation in T cell subtype proportions between adjacent non-tumor and tumor samples, based on the public scRNA-seq data.

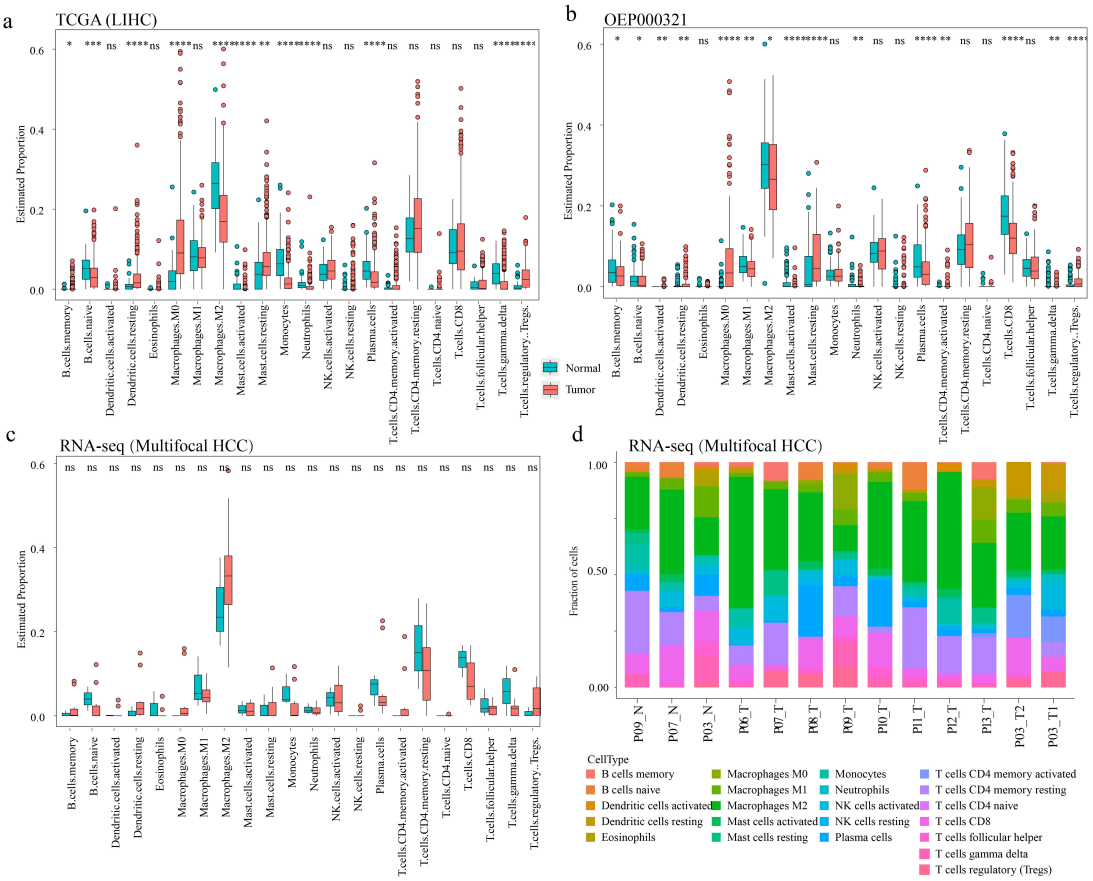

**Figure 11. Immune cell deconvolution reveals immune landscape differences between tumor and non-tumor tissues across TCGA-LIHC, OEP000321, and our multifocal HCC cohorts based on bulk RNA-seq data. (a,b)** Estimated proportions of 22 immune cell types in tumor and adjacent normal tissues in the TCGA-LIHC cohort **(a)** and OEP000321 cohort **(b)** using CIBERSORT deconvolution. **(c)** Immune cell composition in multifocal HCC tumors and matched non-tumor samples from bulk RNA-seq data. **(d)** Stacked bar plots showing the relative fractions of immune cell types across tumor and peritumoral normal samples in multifocal HCC from RNA-seq data. Cell types were inferred using the LM22 signature.

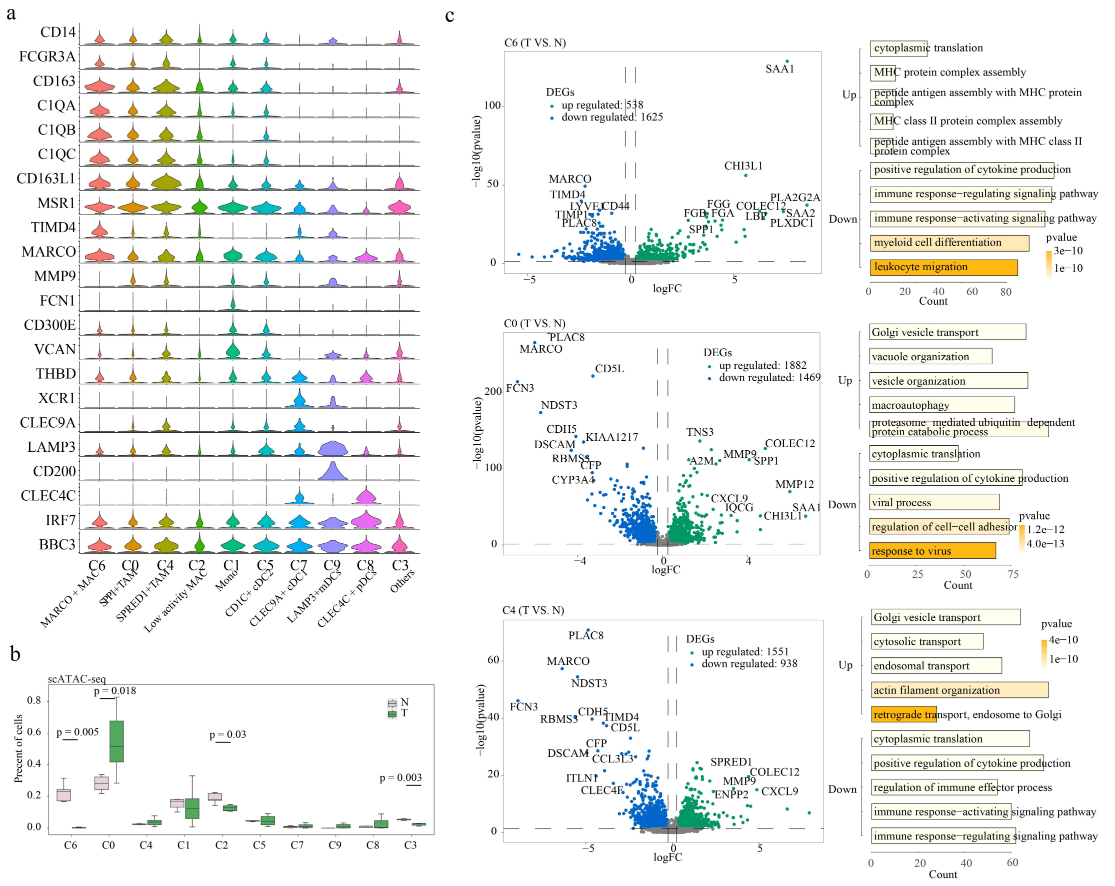

**Figure 12. Single-cell epigenomic characterization of myeloid cell subtypes and differential gene expression between tumor and adjacent non-tumor tissues. (a)** Violin plots displaying the expression of subtype-specific marker genes across Mye cell subtypes, based on scATAC-seq data. **(b)** Boxplots showing the variation in Mye cell subtype proportions between adjacent non-tumor (N) and tumor (T) samples, based on scATAC-seq data. **(c)** Volcano plots showing differentially expressed genes between adjacent non-tumor (N) and tumor (T) tissues for the C6, C0, and C4 clusters (left), and functional enrichment analysis of upregulated and downregulated genes (right).

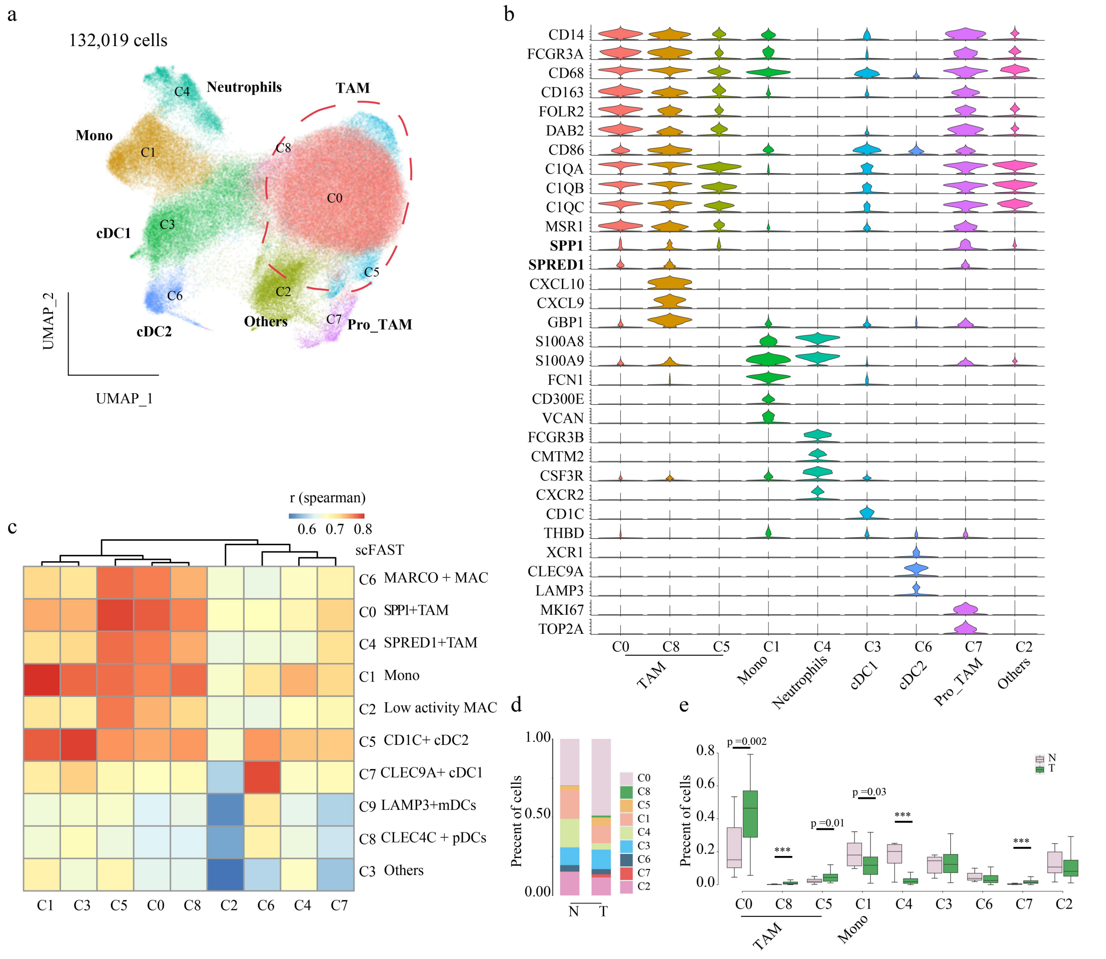

**Figure 13. Validation of myeloid cell subtype landscape and compositional differences between tumor and adjacent non-tumor tissues using public scRNA-seq data. (a)** UMAP plots showing Mye cell lineages identified by the public scRNA-seq (HRA001748) analysis. **(b)** Violin plots displaying the expression of subtype-specific marker genes across Mye cell subtypes, based on the public scRNA-seq data. **(c)** Correlation analysis between cell clusters based on the public scRNA-seq and our scFAST-seq data. **(d)** Stacked bar charts showing the proportion of Mye cell subtypes in adjacent non-tumor (N) and tumor (T) samples, based on the public scRNA-seq data. **(e)** Boxplots showing the variation in Mye cell subtype proportions between adjacent non-tumor (N) and tumor (T) samples, based on the public scRNA-seq data.

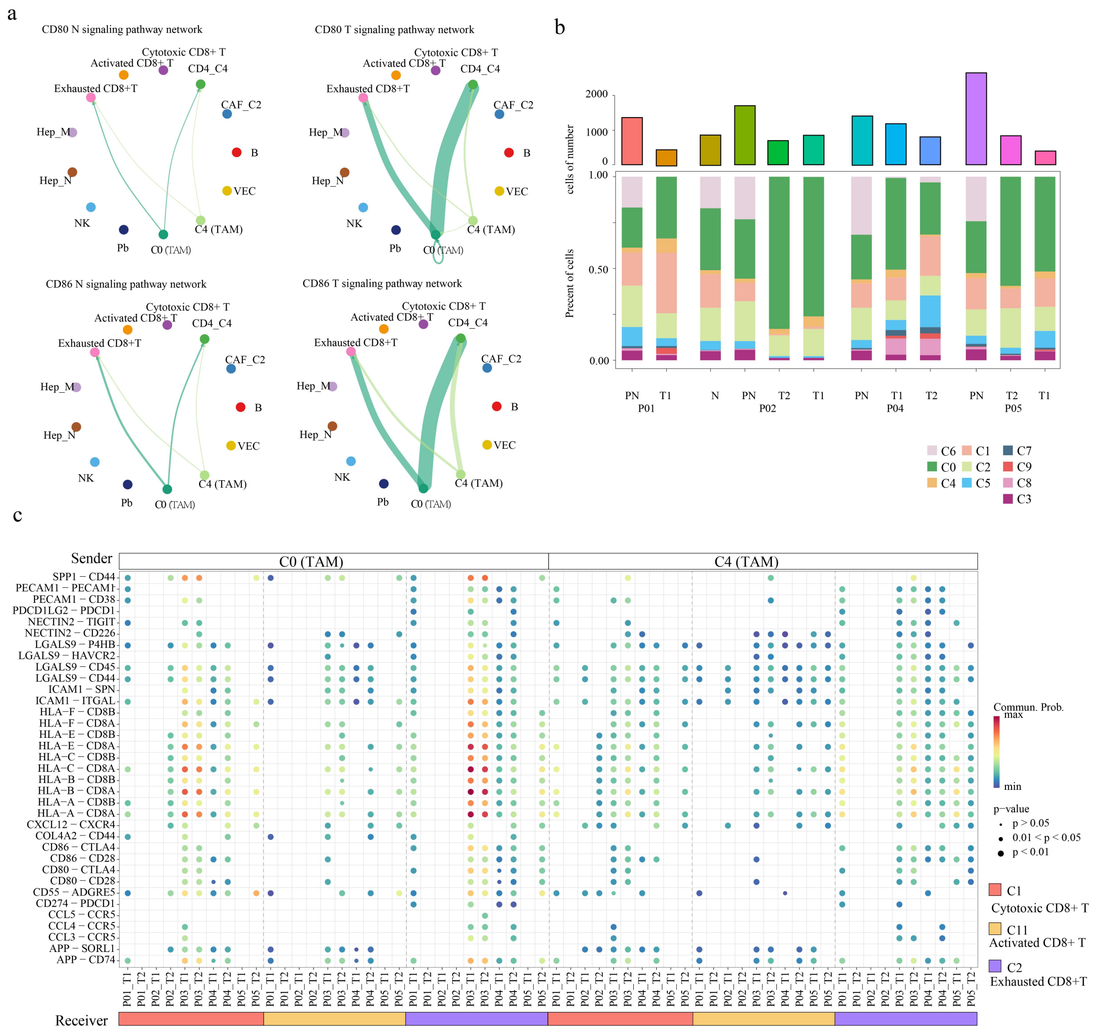

**Figure 14. Myeloid-T cell interaction landscape and tumor focus-specific immune cell composition across multifocal HCC. (a)** Chord diagram showing interactions between TAM cells and CD8+ T cells, mediated by CD80 and CD86 signaling pathways, in adjacent non-tumor (N, left) and tumor (T, right) tissues. **(b)** Stacked bar charts displaying the Mye cell subtype composition across different foci, based on scATAC-seq data. **(c)** Bubble plot indicating the interaction strength of ligand–receptor pairs between SPP1+ TAMs (C0) / SPRED1+ TAMs (C4) and CD8+ T cells across tumor foci.

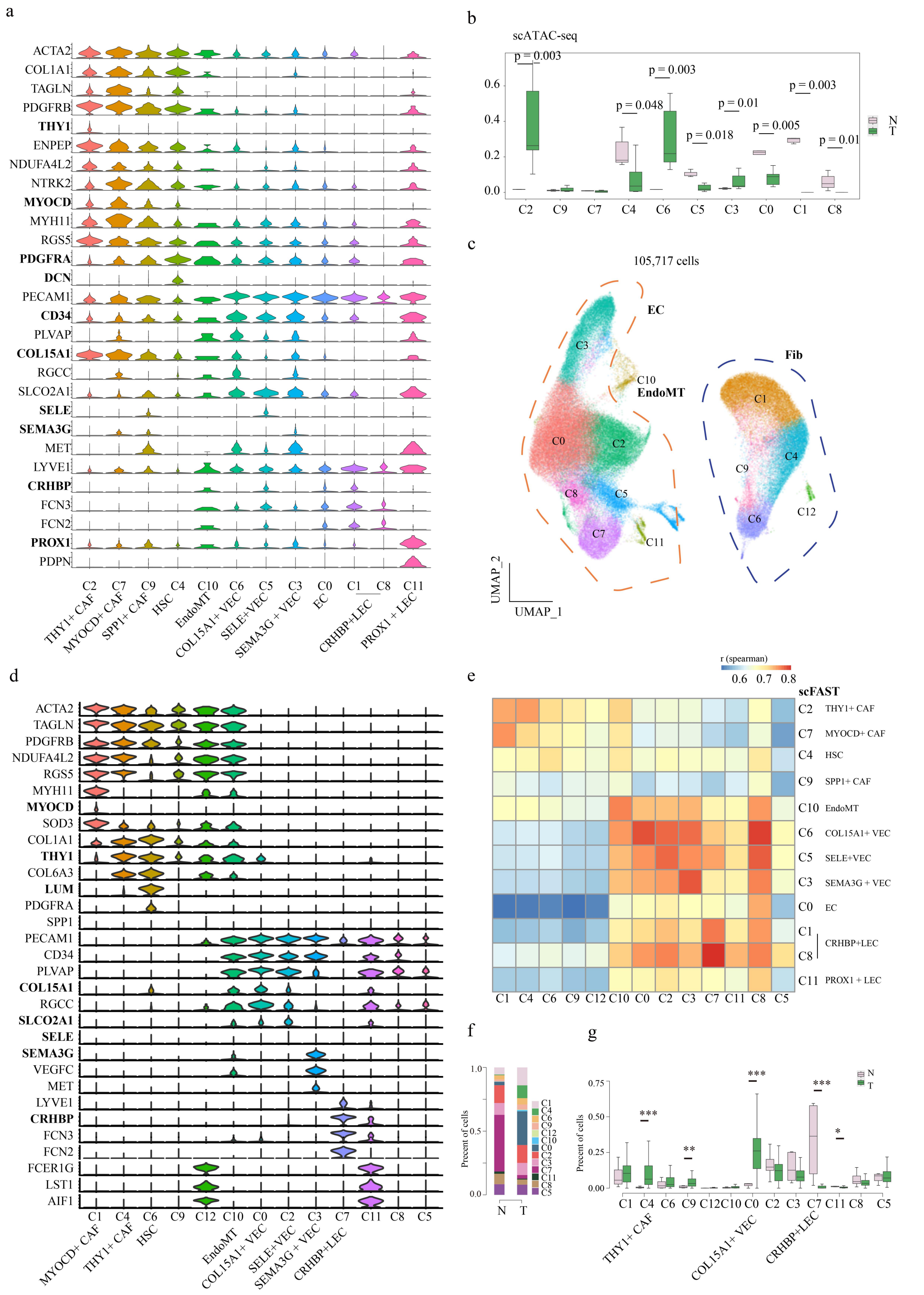

**Figure 15. Integrated scATAC-seq and scFAST-seq analysis, together with validation using public scRNA-seq datasets, reveals stromal cell subtype landscapes and their compositional changes between tumor and adjacent non-tumor tissues. (a)** Violin plots displaying the expression

of subtype-specific marker genes across stromal cell subtypes, based on scATAC-seq data. **(b)** Boxplots showing the variation in stromal cell subtype proportions between adjacent non-tumor (N) and tumor (T) samples, based on scATAC-seq data. **(c)** UMAP plots showing stromal cell lineages identified by the public scRNA-seq (HRA001748). **(d)** Violin plots displaying the expression of subtype-specific marker genes across stromal cell subtypes, based on the public scRNA-seq data. **(e)** Correlation analysis between stromal cell subtypes based on the public scRNA-seq and our scFAST-seq data. **(f)** Stacked bar charts showing the proportion of stromal cell subtypes in adjacent non-tumor and tumor samples, based on the public scRNA-seq data. **(g)** Boxplots showing the variation in stromal cell subtype proportions between adjacent non-tumor (N) and tumor (T) samples, based on public scRNA-seq data.

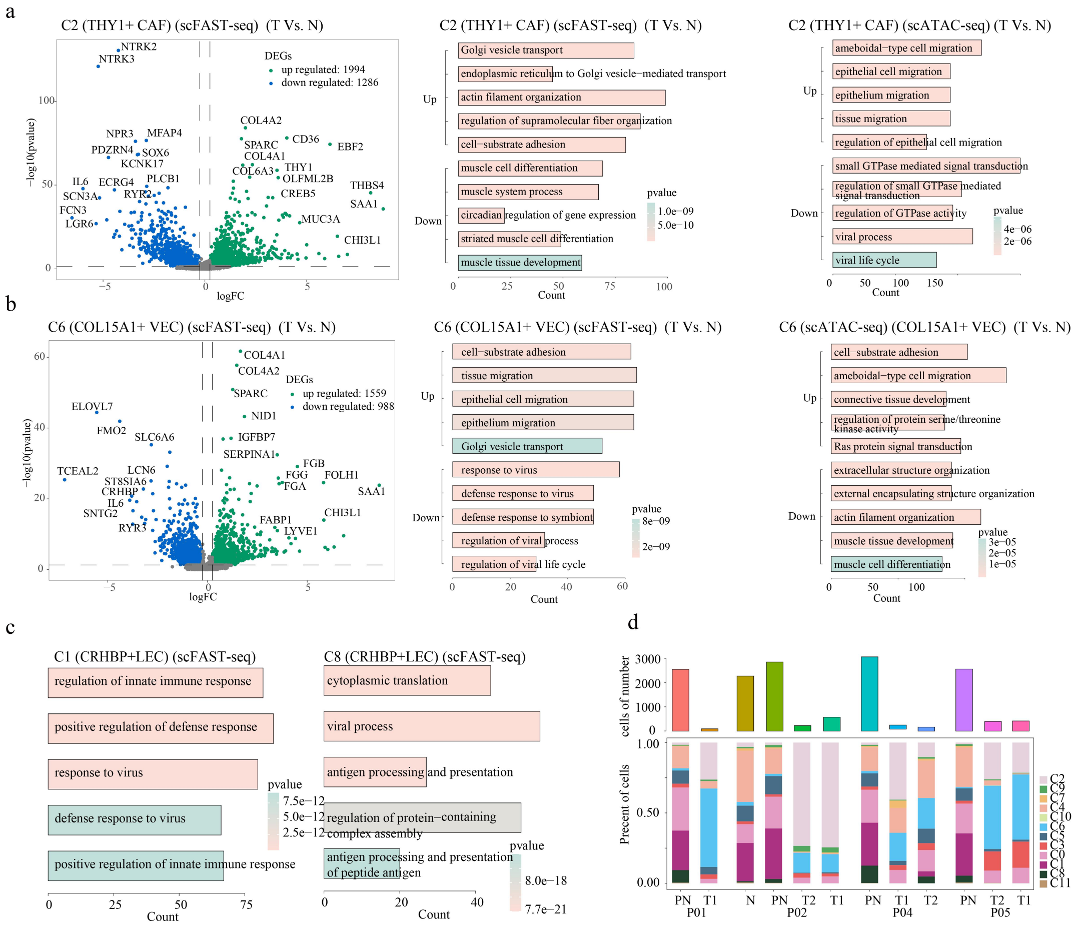

**Figure 16. Transcriptomic and epigenomic characterization of stromal cell subtypes reveals differential gene expression, chromatin accessibility, and compositional changes across tumor foci. (a,b)** Volcano plots (left) showing differentially expressed genes between adjacent non-tumor (N) and tumor (T) tissues for C2 (THY1+CAFs) **(a)** and C6 (COL15A1+ VECs) **(b)** , functional enrichment analysis of upregulated and downregulated genes based on scFAST-seq data (middle), and functional enrichment of genes associated with up- and downregulated peaks from scATAC-seq (right). **(c)** Functional enrichment analysis of highly expressed genes in C1 and C8 (CRHBP+ LECs). **(d)** Stacked bar charts displaying the stromal cell subtype composition across different foci, based on scATAC-seq data.

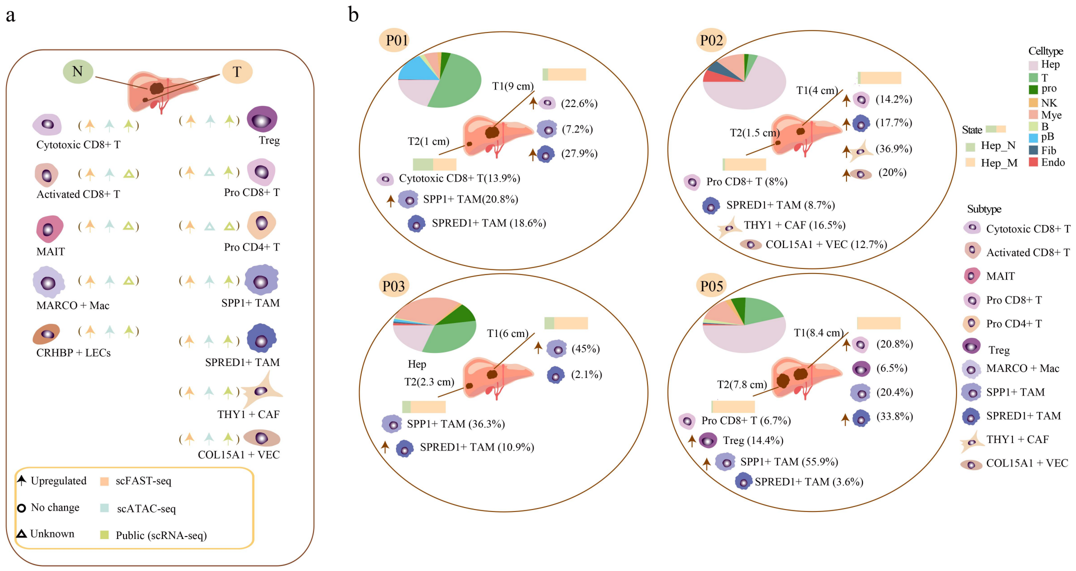

**Figure 17. Overview of dynamic alterations in immune and stromal cell populations across adjacent non-tumor and tumor tissues in multifocal hepatocellular carcinoma. (a)** Overview of the dynamic changes in key immune and stromal cell clusters between adjacent non-tumor and tumor tissues in multifocal HCC. **(b)** Changes in the proportions of cell subtypes across different foci in individual patients. Pie charts represent the distribution of major cell types per patient.

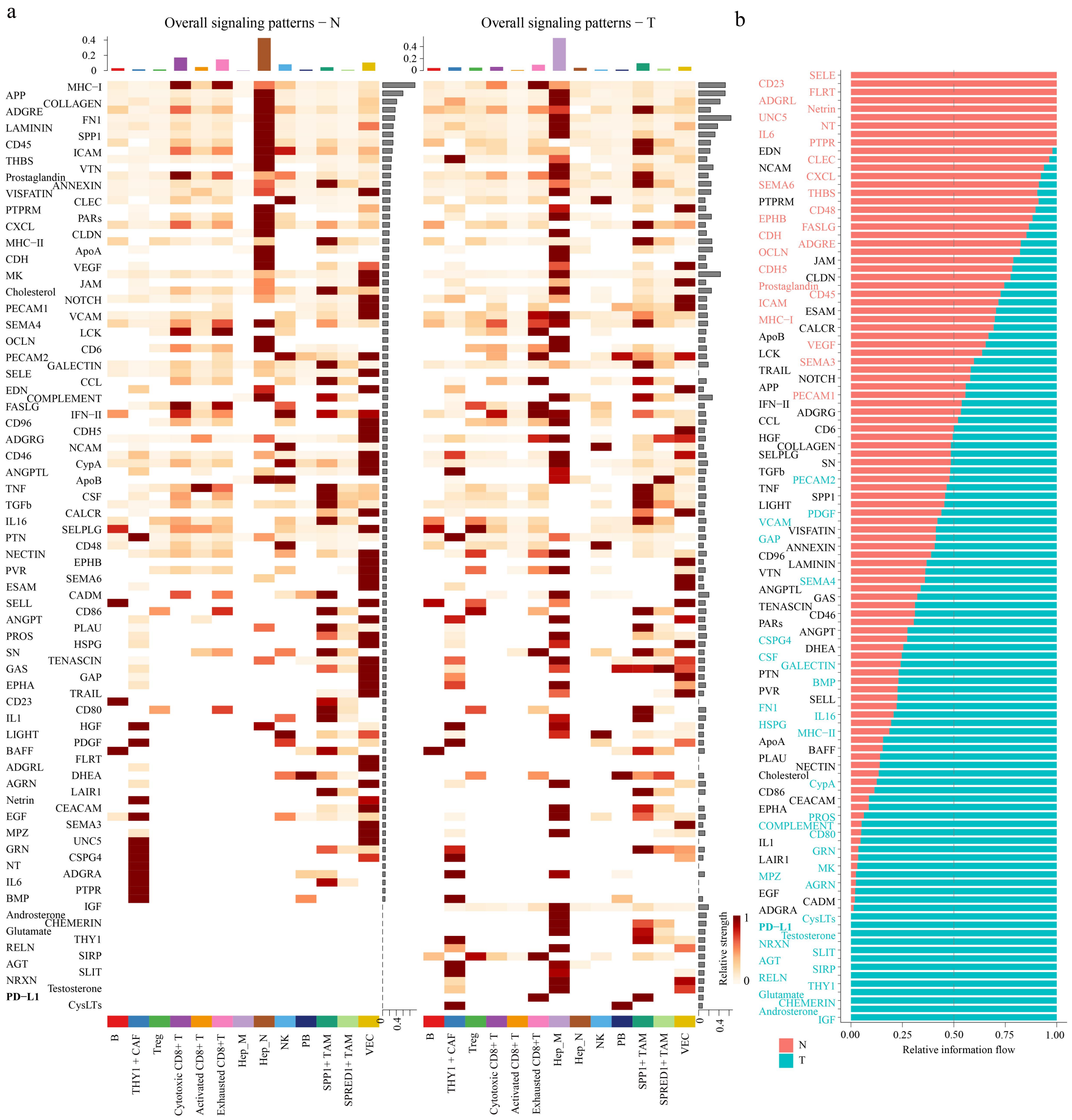

**Figure 18. Comparative analysis of outgoing and incoming intercellular signaling pathway strengths across cell types and between tumor and adjacent non-tumor tissues. (a)** Heatmap showing the relative strength of outgoing signaling pathways (left) and incoming signaling pathways (right) across cell types. The color represents the relative signaling strength. The right gray bar plot summarizes the total signaling strength of each pathway by aggregating all cell types displayed in the heatmap. **(b)** Relative signaling strength of different pathways in adjacent non-tumor (N) and tumor (T) tissues.

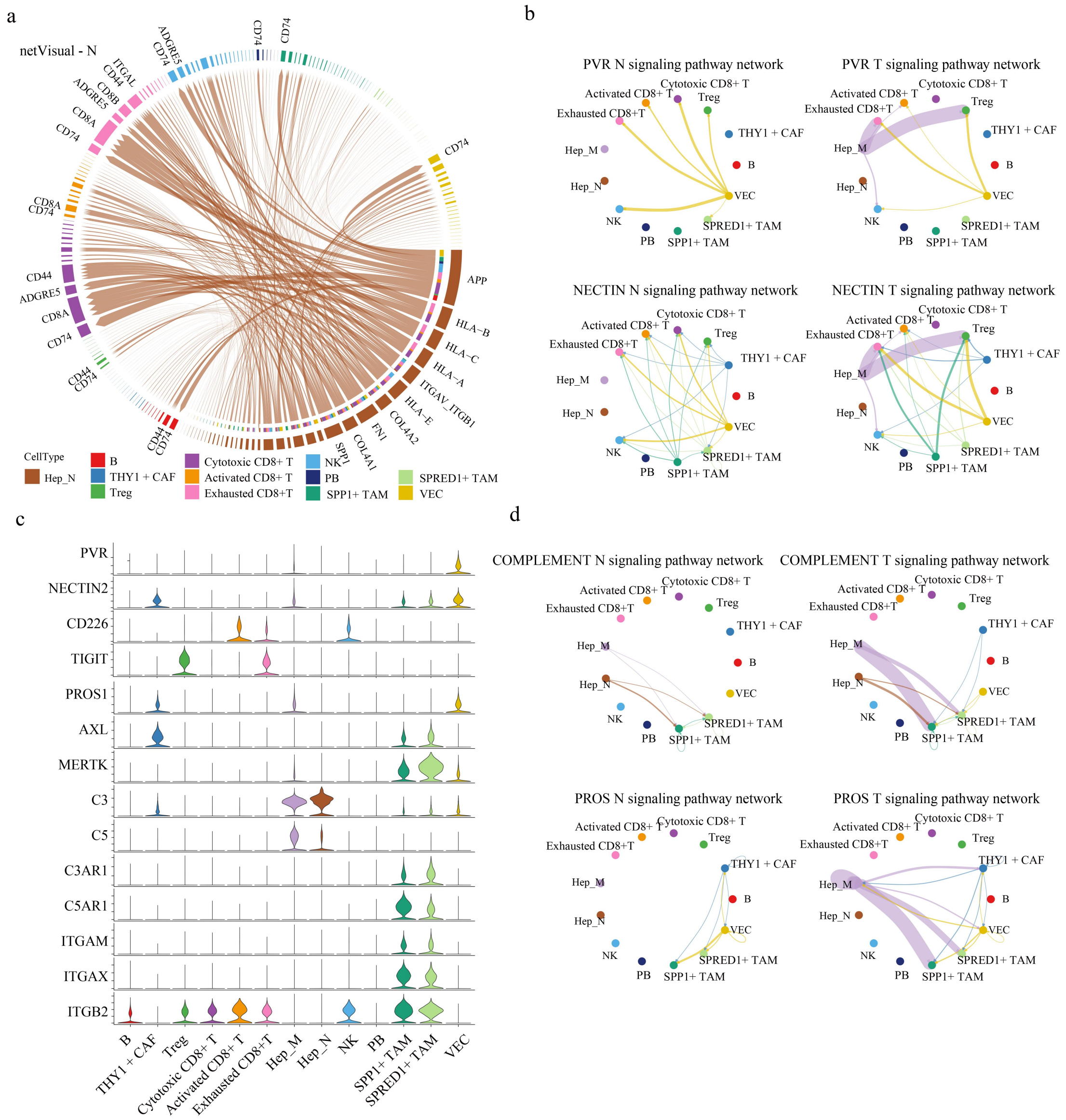

**Figure 19. Cell-cell communication networks reveal key ligand-receptor interactions and signaling pathways in adjacent non-tumor and tumor tissues. (a)** Chord diagram showing that Hep_N cells regulate other cell populations in adjacent non-tumor samples via ligand-receptor interactions. **(b)** Chord diagram depicting interactions mediated by PVR and NECTIN2 signaling pathways in adjacent non-tumor (N,left) and tumor (T, right) tissues. **(c)** Expression profiles of PVR, NECTIN2, COMPLEMENT, and PROS-related ligand and receptor genes across different cell types. **(d)** Chord diagram depicting interactions mediated by COMPLEMENT and PROS signaling pathways in adjacent non-tumor (N,left) and tumor (T, right) tissues.

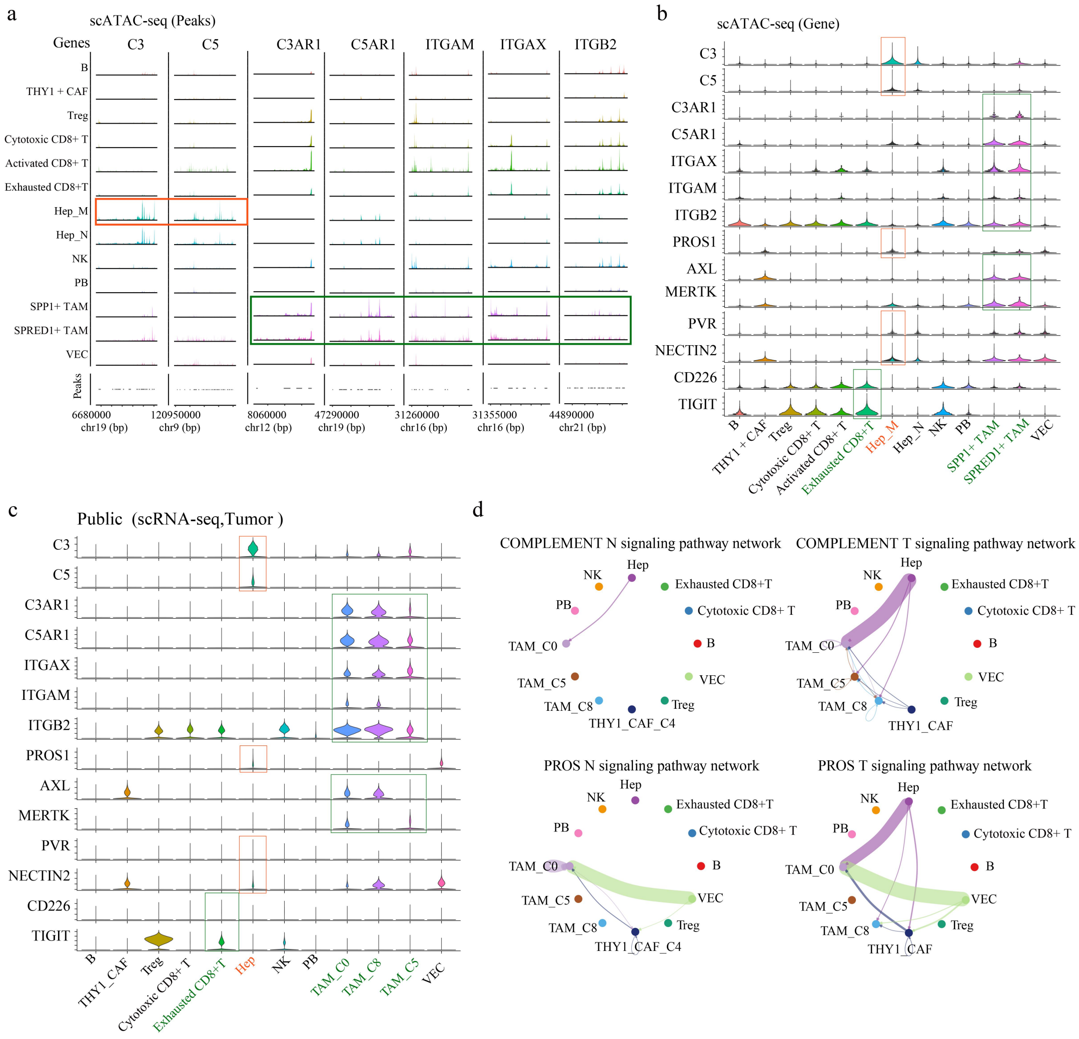

**Figure 20. Integrated scATAC-seq and scRNA-seq analysis of PVR, NECTIN2, COMPLEMENT, and PROS signaling pathways reveals cell-type-specific ligand-receptor activity and intercellular communication networks. (a)** Peaks of COMPLEMENT-related ligands and receptors across different cell types, based on scATAC-seq data. **(b)** Violin plots showing the expression of PVR, NECTIN2, COMPLEMENT, and PROS-related ligand and receptor genes, based on scATAC-seq data. **(c)** Violin plots showing the expression of PVR, NECTIN2, COMPLEMENT, and PROS-related ligand and receptor genes, based on the public scRNA-seq data. **(d)** Chord diagram showing the COMPLEMENT and PROS signaling pathways in adjacent non-tumor (N,left) and tumor (T, right) tissues, based on the public scRNA-seq data.

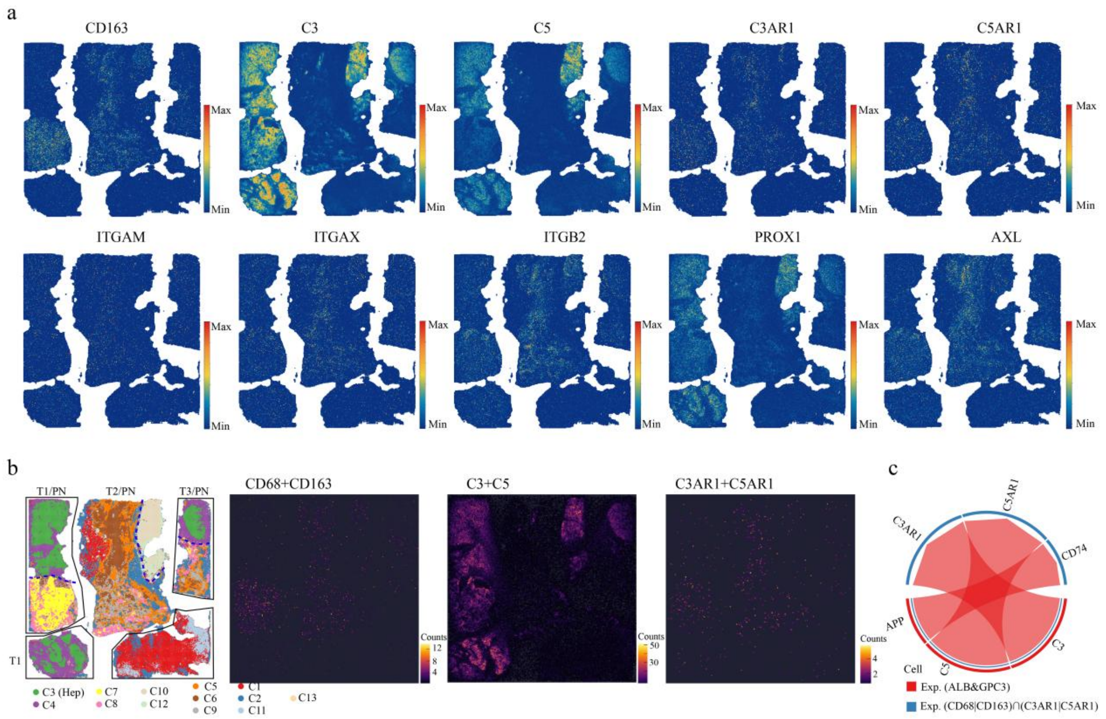

**Figure 21. Spatial transcriptomic analysis reveals expression patterns and spatial activation of COMPLEMENT signaling in tumor tissue sections. (a)** Spatial expression patterns of COMPLEMENT-related ligand and receptor genes in P05 tissue sections, based on Stereo-seq. **(b)** Spatial mapping of COMPLEMENT-related genes in P05 patient slices based on Stereo-seq data. **(c)** Chord diagram illustrating the spatial activation patterns of the COMPLEMENT signal.

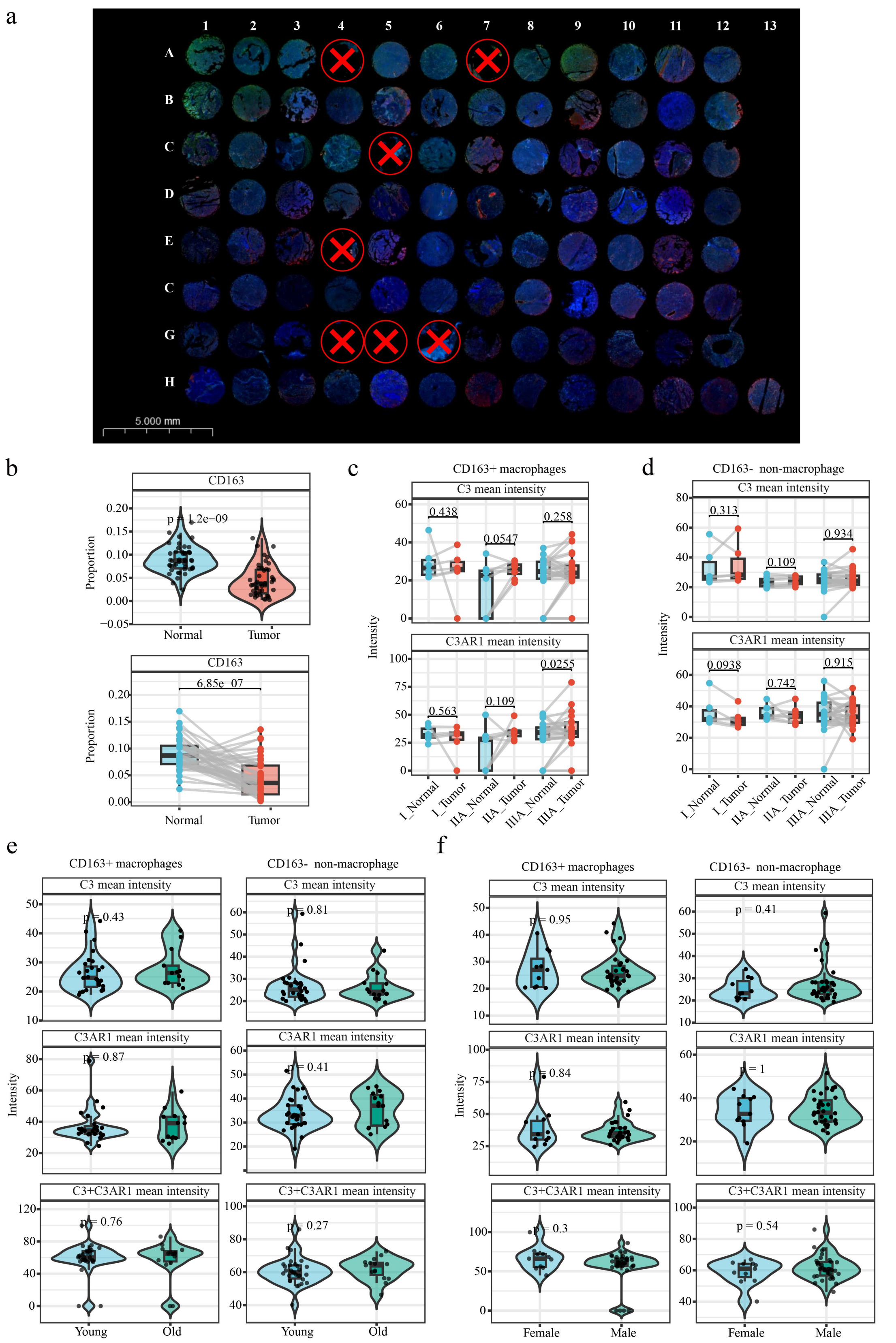

**Figure 22. Validation of tumor-associated C3–C3AR1 activation in CD163+ macrophages by multiplex immunohistochemistry. (a)** Representative image of the tissue microarray containing 48 paired multifocal

HCC tumor and adjacent tissue specimens. **(b)** Quantification of CD163+ macrophage proportions in tumor and adjacent normal tissues. **(c,d)** C3 and C3AR1 expression intensity in CD163+ macrophages **(c)** and CD163− non-macrophage populations **(d)** across different tumor stages. **(e,f)** C3 and C3AR1 expression intensity in CD163+ macrophages and CD163− non-macrophage populations according to patient age **(e)** and sex **(f)**.

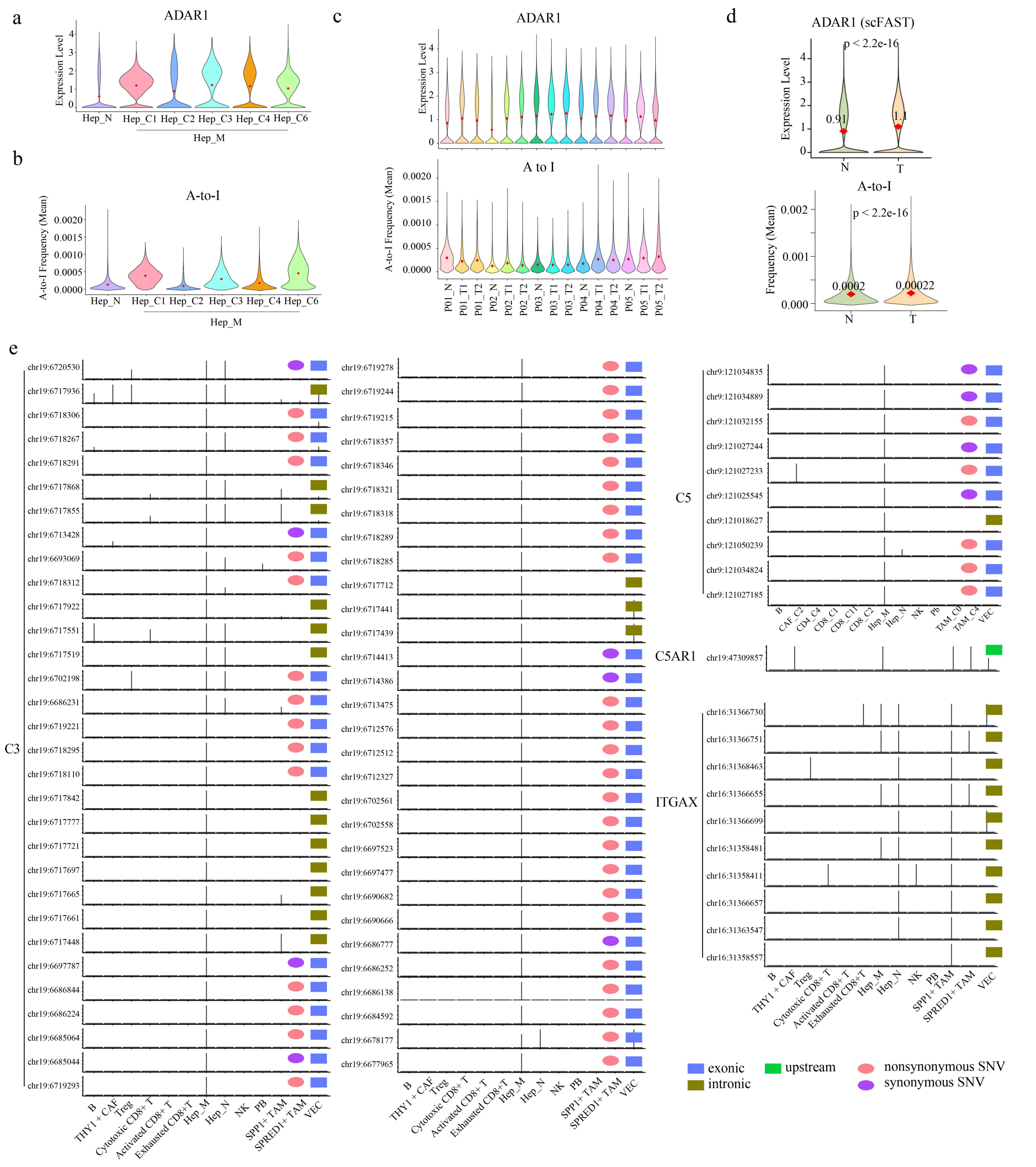

**Figure 23. ADAR-associated RNA editing landscape links malignant hepatocytes to complement signaling regulation. (a–c)** *ADAR* expression levels and global A-to-I RNA editing frequencies across different Hep subtypes **(a, b)** and across individual samples **(c)**. **(d)** *ADAR* expression and global A-to-I editing in tumor (T) vs. Adjacent normal (N) tissues by scFAST-seq. **(e)** Bar plots showing the distribution of A-to-I RNA editing sites in selected complement signaling genes (*C3*, *C5*, *C5AR1*, and *ITGAX*) across different cell types.

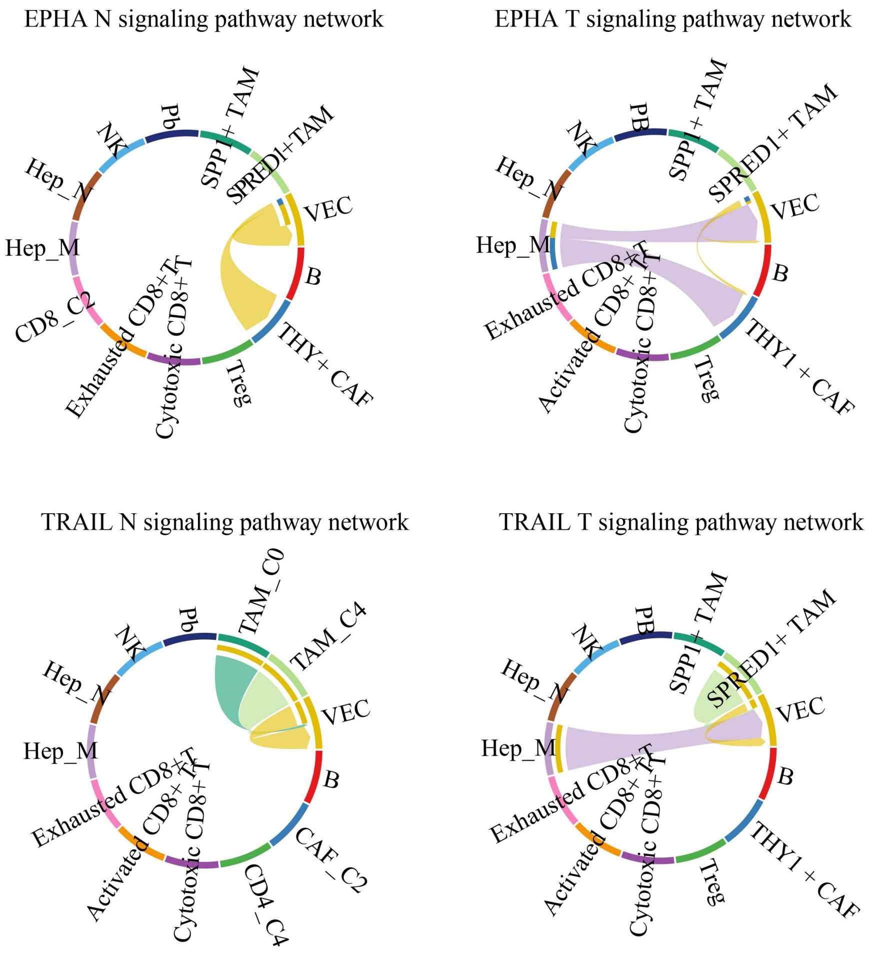

**Figure 24. EPHA and TRAIL signaling-mediated cell-cell interaction networks in adjacent non-tumor (N) and tumor (T) tissues.**

##### 3. Supplementary references

[1] Xue R, Zhang Q, Cao Q, et al. Liver tumour immune microenvironment subtypes and neutrophil heterogeneity. Nature 2022;612:141-147.

[2] Jiang Z, Wu Y, Miao Y, et al. HCCDB v2.0: Decompose Expression Variations by Single-cell RNA-seq and Spatial Transcriptomics in HCC. Genomics Proteomics Bioinformatics 2024;22.

[3] Hao Y, Stuart T, Kowalski MH, et al. Dictionary learning for integrative, multimodal and scalable single-cell analysis. Nat Biotechnol 2024;42:293-304.

[4] Mansi L, Tangaro MA, Lo Giudice C, et al. REDIportal: millions of novel A-to-I RNA editing events from thousands of RNAseq experiments. Nucleic Acids Res 2021;49:D1012-D1019.

[5] Wang K, Li M, Hakonarson H. ANNOVAR: functional annotation of genetic variants from high-throughput sequencing data. Nucleic Acids Res 2010;38:e164.

[6] Ramakrishnan A, Symeonidi A, Hanel P, et al. epiAneufinder identifies copy number alterations from single-cell ATAC-seq data. Nat Commun 2023;14:5846.

[7] Kang M, Armenteros JJA, Gulati GS, et al. Mapping single-cell developmental potential in health and disease with interpretable deep learning. bioRxiv 2024.

[8] Kurtenbach S, Cruz AM, Rodriguez DA, et al. Uphyloplot2: visualizing phylogenetic trees from single-cell RNA-seq data. BMC Genomics 2021;22:419.

[9] Chen B, Khodadoust MS, Liu CL, et al. Profiling Tumor Infiltrating Immune Cells with CIBERSORT. Methods Mol Biol 2018;1711:243-259.

[10] Trapnell C, Cacchiarelli D, Grimsby J, et al. The dynamics and regulators of cell fate decisions are revealed by pseudotemporal ordering of single cells. Nat Biotechnol 2014;32:381-386.

[11] Jin S, Plikus MV, Nie Q. CellChat for systematic analysis of cell-cell communication from single-cell transcriptomics. Nat Protoc 2025;20:180-219.
